# High-Spatial-Resolution Multi-Omics Atlas Sequencing of Mouse Embryos via Deterministic Barcoding in Tissue

**DOI:** 10.1101/788992

**Authors:** Yang Liu, Mingyu Yang, Yanxiang Deng, Graham Su, Archibald Enninful, Cindy C. Guo, Toma Tebaldi, Di Zhang, Dongjoo Kim, Zhiliang Bai, Eileen Norris, Alisia Pan, Jiatong Li, Yang Xiao, Stephanie Halene, Rong Fan

## Abstract

We present DBiT-seq –Deterministic Barcoding in Tissue for spatial omics sequencing – for co-mapping of mRNAs and proteins in a formaldehyde-fixed tissue slide via NGS sequencing. Parallel microfluidic channels were used to deliver DNA barcodes to the surface of a tissue slide and crossflow of two sets of barcodes A1-50 and B1-50 followed by ligation *in situ* yielded a 2D mosaic of tissue pixels, each containing a unique full barcode AB. Application to mouse embryos revealed major tissue types in early organogenesis as well as fine features like microvasculature in a brain and pigmented epithelium in an eye field. Gene expression profiles in 10μm pixels conformed into the clusters of single-cell transcriptomes, allowing for rapid identification of cell types and spatial distributions. DBiT-seq can be adopted by researchers with no experience in microfluidics and may find applications in a range of fields including developmental biology, cancer biology, neuroscience, and clinical pathology.

**In Brief:** Microfluidic deterministic barcoding of mRNAs and proteins in tissue slides followed by high-throughput sequencing enables the construction of a high-spatial-resolution multi-omics atlas at the genome scale. Application to mouse embryos (E10-12) identified major tissue types in early organogenesis and revealed fine tissue features such as retinal pigmented epithelium and endothelial microvasculature at the cellular level.

**Highlights:** - Deterministic barcoding in tissue enables NGS-based spatial multi-omics mapping.
- DBiT-seq identified spatial patterning of major tissue types in mouse embryos.
- DBiT-seq revealed fine features such as retinal pigmented epithelium and microvascular endothelium at the cellular level.
- Direct integration with scRNA-seq data allows for rapid cell type identification.

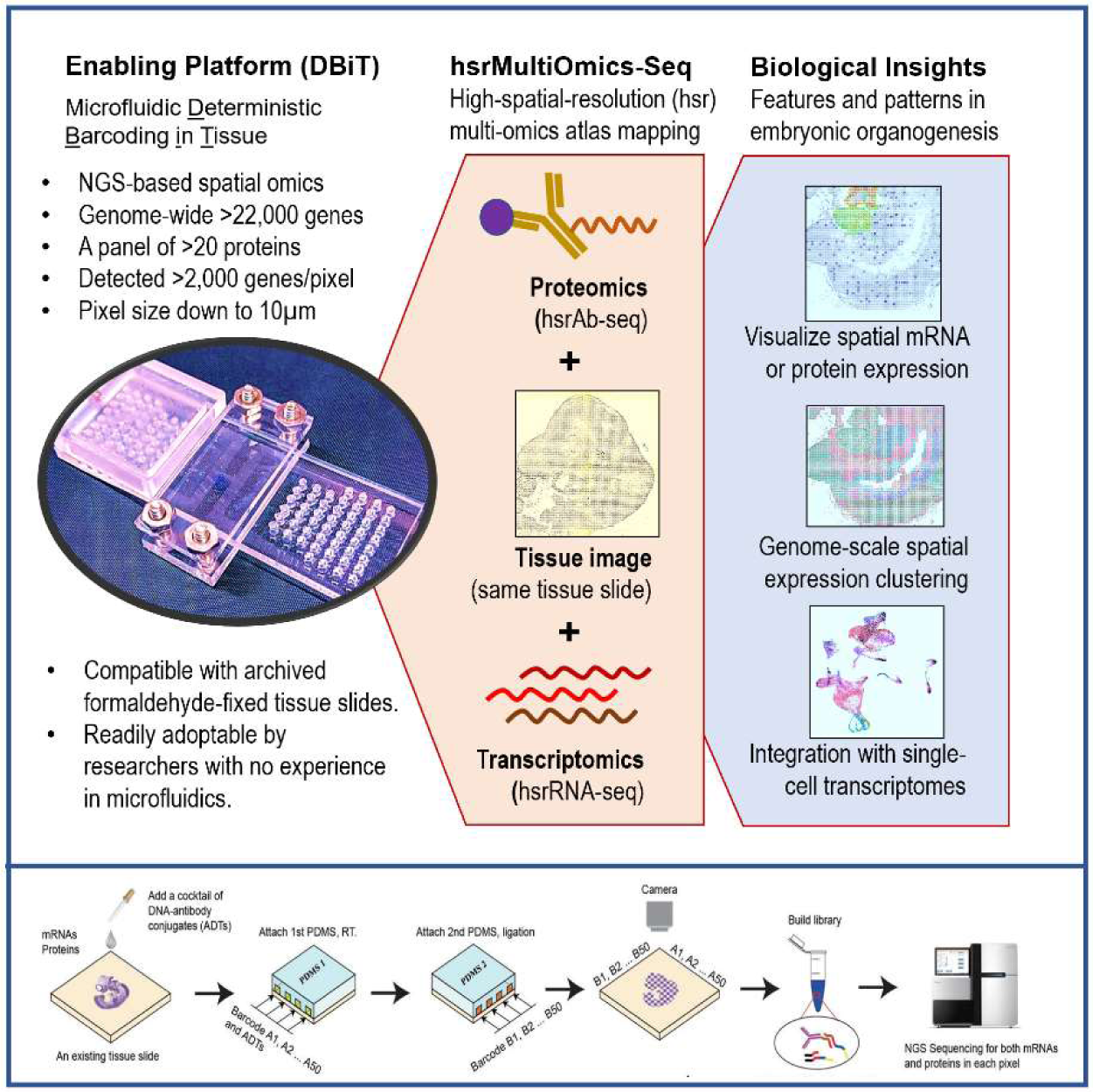

## INTRODUCTION

In multicellular systems, cells do not function in isolation but are strongly influenced by spatial location and surroundings(Knipple et al., 1985; Scadden, 2014; van Vliet et al., 2018). Spatial gene expression heterogeneity plays an essential role in a range of biological, physiological and pathological processes(de Bruin et al., 2014; Fuchs et al., 2004; Yudushkin et al., 2007). For example, how stem cells differentiate and give rise to diverse tissue types is a spatially regulated process which controls the development of different tissue types and organs(Ivanovs et al., 2017; Slack, 2008). Mouse embryonic organogenesis begins at the end of the first week, follows gastrulation, and continues through birth(Mitiku and Baker, 2007). When and how exactly different organs emerge in an early embryo is still inadequately understood due to highly dynamic spatial organization of tissues and cells at this stage. An embryonic organ could differ substantially in anatomical and molecular definitions as compared to the adult counterpart. In order to dissect the initiation of early organogenesis at the whole embryo scale, it is desirable to not only measure genome-wide molecular profiles for cell type identification but also interrogate spatial organization in the tissue context at a high resolution.

Despite the latest advent of massively parallel single-cell RNA-sequencing (scRNA-seq) (Klein et al., 2015; Macosko et al., 2015) that revealed astonishing cellular heterogeneity in many tissue types, including the dissection of all major cell types in developing mouse embryos from E9 to E14(Cao et al., 2019; Pijuan-Sala et al., 2019), the spatial information in the tissue context is missing in scRNA-seq. Spatial transcriptomics emerged to address this problem(Burgess, 2019). Early attempts were all based on multiplexed single-molecule fluorescent *in situ* hybridization (smFISH) via spectral barcoding and/or sequential imaging(Pichon et al., 2018; Trcek et al., 2017). It evolved rapidly over the past years from detecting a handful of genes to hundreds or thousands (e.g., seqFISH, MERFISH) (Chen et al., 2015; Lubeck et al., 2014), and recently to the whole transcriptome level (e.g, SeqFISH+) (Eng et al., 2019). However, these methods were technically demanding, requiring high-sensitivity single-molecule fluorescence imaging systems, sophisticated image analysis processes, and a lengthy repeated imaging workflow to achieve high multiplexing (Perkel, 2019). Moreover, they were all based upon a finite panel of probes that hybridize to known mRNA sequences, limiting their potential to discover new sequences and variants. Fluorescent *in situ* sequencing methods (e.g., FISSEQ, STARmap) (Lee et al., 2015; Wang et al., 2018) were additionally reported but the number of detectable genes was limited, and their workflow resembled sequential FISH, again requiring a lengthy, repeated, and technically demanding imaging process.

It is highly desirable to develop new methods for high-spatial-resolution, unbiased, genome-scale molecular mapping in intact tissues at cellular level, which does not require sophisticated imaging but capitalize on the power of Next Generation Sequencing (NGS) to achieve higher sample throughput and cost efficiency. Spatial transcriptome mapping at cellular level (spot size ∼10µm) was demonstrated with Slide-seq that utilized a self-assembled monolayer of DNA-barcoded beads on a glass slide to capture mRNAs released from a frozen tissue section placed on top (Rodriques et al., 2019). A similar method, called high-definition spatial transcriptome (HDST), used 2µm beads in a microwell array chip to further increase resolution (Vickovic et al., 2019). However, these emergent methods are limited by low number of detected genes (∼150 genes per pixel in Slide-seq), incompatibility with fixed tissues, potential lateral diffusion of release mRNAs, and sophisticated process for bead decoding. Moreover, they are all limited to spatial transcriptomes and yet to realize multi-omics spatial sequencing.

We sought to develop a completely different approach, which was to spatially barcode biomolecules in tissues rather than to capture them on a solid-phase substrate. Previously, we developed microfluidic channel-guided patterning of DNAs or antibodies on a glass slide for multiplexed protein assay(Lu et al., 2013; Lu et al., 2015). We speculated that a microfluidics-confined delivery of molecular barcodes to a tissue section could enable high-spatial-resolution barcoding of mRNAs or proteins directly in tissue. Herein, we report on Deterministic Barcoding in Tissue for spatial omics sequencing (DBiT-seq). A microfluidic chip with parallel channels (10, 25 or 50μm in width) was placed directly against a fixed tissue slide to introduce oligo-dT tagged DNA barcodes A1-A50 that annealed to mRNAs to initiate *in situ* reverse transcription. This step resulted in stripes of barcoded cDNAs inside the tissue. Afterwards, the first microfluidic chip was removed and another chip was placed on the same tissue slide with the microchannels perpendicular to the first flow direction to introduce a second set of DNA barcodes B1-B50, which were subsequently ligated at the intersections to form a 2D mosaic of tissue pixels, each containing a distinct combination of barcodes Ai and Bj (i=1-50, j=1-50). Then, the tissue was digested to recover spatially barcoded cDNAs that were collected to an Eppendorf tube, template-switched, PCR amplified, and tagmentated to prepare a library for NGS sequencing. Proteins could be co-measured by applying a cocktail of antibody-derived DNA tags (ADTs) to the fixed tissue slide prior to flow barcoding, similar to Ab-seq or CITE-seq(Shahi et al., 2017; Stoeckius et al., 2017). We demonstrated high-spatial-resolution co-mapping of whole transcriptome and a panel of 22 proteins in mouse embryos (E10-12). DBiT-seq faithfully detected all major tissue types in early organogenesis and identified the fine features such as brain microvascular networks and a single-cell-layer of melanocytes lining an optical vesicle. We found that the gene expression profiles of 10µm tissue pixels were dominated by single-cell transcriptomes and an integrated analysis allowed for rapid identification of cell types in relation to spatial distribution. Besides, the microfluidic chip was directly clamped onto the tissue slide and the reagent dispensing was performed by directly pipetting into the inlet holes, requiring no prior experience in microfluidic control. Thus, DBiT-seq could be readily adopted by researchers from a wide range of fields in biological and biomedical research.

## RESULTS

### DBiT-seq workflow

The workflow of DBiT-seq is described in Figure 1A (also see Figure S1). A tissue section pre-fixed with formaldehyde on a standard aminated glass slide was used. A polydimethylsiloxane (PDMS) microfluidic chip (Figure S2) containing 50 parallel microchannels (down to 10μm in width) was placed on the tissue slide to introduce a set of DNA barcode A solutions. To assist the assembly, an acrylic clamp was used to hold the PDMS chip firmly against the tissue slide (Figure 1B). The inlet holes were ∼2mm in diameter and ∼4mm in depth allowing for ∼5μL reagents to be directly pipetted into the inlets. The outlet holes were roofed with a global cover connected to a house vacuum to pull the reagents all the way from the inlets to the outlets through the tissue surface, which took several seconds for a 50μm chip and up to 3min for a 10μm chip. Barcode A is composed of an oligo-dT sequence for binding mRNAs, a distinct spatial barcode Ai (i=1-50, 8mer), and a ligation linker (15mer). Reverse transcription was conducted during the first flow for *in situ* synthesis of first strand cDNAs that immediately incorporate barcode A. Then, the first PDMS chip was removed and another PDMS chip with the microchannels perpendicular to those in the first flow barcoding was placed on the same tissue to introduce a second set of barcodes Bj (j=1-50), each containing a ligation linker(15mer), a distinct spatial barcode Bj (j=1-50, 8mer), a unique molecular identifier (UMI), and a PCR handle (22mer) functionalized with biotin, which was used later to perform cDNA purification with streptavidin-coated magnetic beads. Also added to the barcode B reagents were T4 ligase and a complementary ligation linker to perform *in situ* ligation at the intersections, resulting in a mosaic of tissue pixels, each containing a distinct combination of barcodes Ai and Bj (i=1-50, j= 1-50). The tissue slide being processed could be imaged during each flow or afterwards such that the tissue morphology can be correlated with spatial omics map. To co-measure proteins and mRNAs, the tissue slide was stained with a cocktail of 22 antibody-derived DNA tags (ADTs) (Stoeckius et al., 2017) (see Table S1) prior to microfluidic flow barcoding. Each of the ADTs contains a distinct barcode (15mer) and a polyadenylated tail that allowed for protein detection using a workflow similar to that for mRNAs detection. After forming a spatially barcoded tissue mosaic, cDNAs were collected, template-switched, and PCR amplified to make a sequencing library. Using a paired-end (2×100) NGS sequencing, we detected spatial barcodes (AiBj, i=1-50, j=1-50) from one end and the corresponding transcripts and protein barcodes from the other end to computationally reconstruct a spatial expression map. It is worth noting that unlike other methods, DBiT-seq permits the same tissue slide being imaged during or after the flow barcoding (Figure S3) to precisely locate the pixels and perform correlative analysis of tissue morphology and spatial omics maps at high precision. The design of DNA barcodes and the chemistry to perform DBiT-seq are further detailed in Figure S1. The sequences of ADTs and spatial barcodes are summarized in Tables S1&2, respectively.

**Figure 1.**
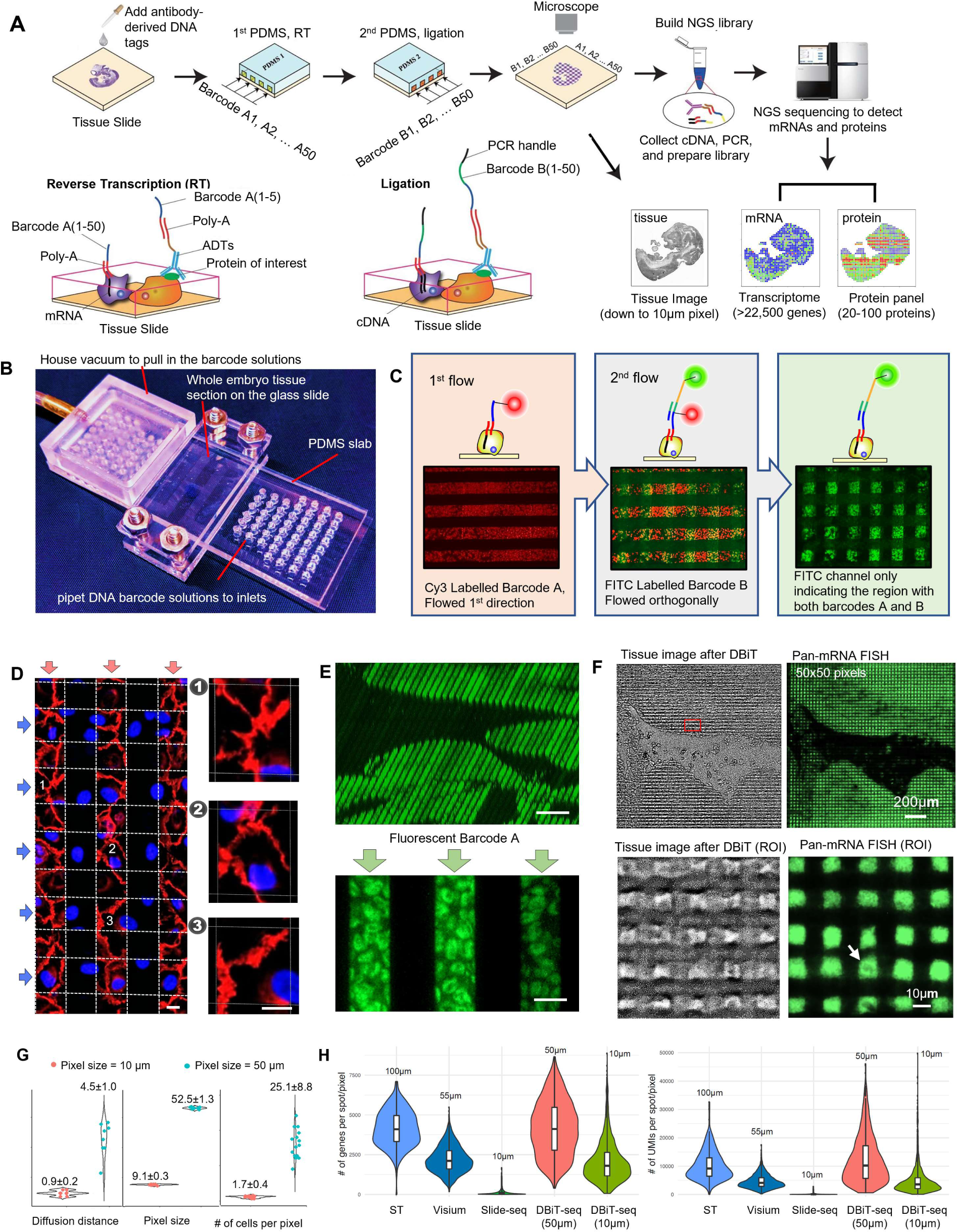
Design and validation of DBiT-seq. (A) Schematic workflow. A standard fixed tissue slide is used as the starting material, which is incubated with a cocktail of antibody-DNA tags that recognize a panel of proteins of interest. A custom-designed PDMS microfluidic chip with 50 parallel microchannels in the center is aligned and placed on the tissue slide to introduce the 1^st^ set of barcodes A1:A50. Each barcode is tethered with a ligation linker and a poly-T tail for binding mRNAs or the DNA tag for protein detection. Then, reverse transcription (RT) is conducted *in situ* to yield cDNAs which are covalently linked to barcodes A1:A50. Afterwards, this microfluidic chip is removed and another microfluidic chip with 50 parallel microchannels perpendicular to those in the first microfluidic chip is placed on the tissue slide to introduce the 2^nd^ set of DNA barcodes B1:B50. These barcodes contain a ligation linker, a unique molecular identifier (UMI) and a PCR handle. After introducing the barcodes B1:B50 and the universal complementary ligation linker through the second microfluidic chip, the barcodes A and B are joined through *in situ* ligation and then the intersection region of the two microfluidic channels defines a distinct pixel with a unique combination of A and B, giving rise to a 2D array of spatially addressable barcodes AiBj (i=1-50, j=1-50). Afterwards, the second PDMS chip is removed and the tissue remains intact while spatially barcoded for all mRNAs and the proteins of interest. The barcoded tissue is imaged under an optical or fluorescence microscope to visualize individual pixels. Finally, cDNAs are extracted from the tissue slide, template switched to add a PCR handle to the other side and amplified by PCR for making the sequencing via tagmentation. A paired end high-throughput sequencing is performed to read the spatial barcodes (AiBj) and cDNA sequences from either mRNAs or antibody DNA tags. In silicon reconstruction of a spatial mRNA or protein expression map is realized by matching the spatial barcodes AiBj to the corresponding cDNA reads. The reconstructed spatial omics atlas can be correlated to the tissue image taken after DBiT-seq to identify the exact spatial location of individual pixels. (B) Microfluidic device used in DBiT-seq. A series of microfluidic chips were fabricated with 50 parallel microfluidic channels in the center that are 50μm, 25μm, and 10μm in width, respectively. The PDMS chip containing 50 parallel channels is placed directly on a tissue slide and, if needed, the center region is clamped using two acrylic plates and screws. All 50 inlets are open holes ∼2mm in diameter and capable of holding ∼13μL of solution. Different barcode solutions are pipetted to these inlets and drawn into the microchannels via a global vacuum applied to the outlets situated on the opposite side of the PDMS chip. Basically, this device is a universal approach to realize spatially defined delivery of DNA barcodes to the tissue surface at a resolution of down to 10μm or even better. (C) Validation of spatial barcoding using fluorescent DNA probes. The images show parallel lines of Cy3-labelled barcode A (red, left panel) on the tissue slide defined by the first flow, the square pixels of FITC-labeled barcode B (green, right panel) defined as the intersection of the first and the second flows, and the overlay of both fluorescence colors (middle). Because barcode B is ligated to the immobilized barcode A in an orthogonal direction, it occurs only at the intersection of the first and second flows. Microfluidic channel width = 50μm. (D) Validation of leak-free flow barcoding with 3T3 cells culturing on glass slides. Cells were stained by flowing DAPI (blue) through the 1^st^ microfluidic chip and phalloidin (red) through the 2^nd^ microfluidic chip. As shown in the enlarged figures, the staining was strictly confined inside the channel. (E) Confocal microscope imaging of stained tissue using fluorescent DNA barcode A. The 3D stack image shows no leakage between two channels. (F) Validation of spatial barcoding for 10μm pixels. A tissue slide was subjected to spatial barcoding or DBiT-seq and the resultant pixels were visualized by optical imaging (upper left) of the tissue post-DBiT-seq and fluorescent imaging (upper right) of the same tissue sample using FITC-labeled barcode B. Pressing microfluidic channels against the tissue slide resulted in a small plastic deformation of the tissue matrix, which enabled us to directly visualize all individual pixels created by DBiT-seq, which turned out to be in excellent agreement with the fluorescent image of all pixels using FITC-labelled barcode B. Enlarged views (low panels) further show discrete pixels down to a 10-μm level. (G) Qualification of the diffusion “leak-out” distance, the measured size of pixels, and the number of cells per pixel. Quantitative analysis of the line profile revealed the barcode diffusion through the dense tissue matrix is small (0.9μm, obtained with 10μm microchannels fixed by clamps). The measured pixel size agreed with the microchannel size. Using a cell nuclear staining DAPI, the number of cells in a pixel can be visualized. The average cell number is 1.7 per a 10μm pixel and 25.1 per a 50μm pixel. Therefore, the pixel size is approaching the single cell level. (H) Gene and UMI counts assessment and comparison. A high-resolution spatial transcriptome sequencing data was obtained using a 10μm microfluidic barcoding approach. The number of genes detected per pixel was >2000. In contrast, Slide-seq only detected ∼150 genes per pixel (10μm), which is insufficient for spatial visualization of individual genes in a meaningful way. The low-resolution ST method yielded a similar number of genes per pixel, but the pixel size is ∼100-150μm, which is almost 100 times larger in area. Similarly, Visium shows comparable genes per spot as DBiT-seq, but with lower resolution.

### Evaluation of the flow barcoding process

Although no obvious leakage was observed by imaging the flow of fluorescent molecules in the microchannels on the tissue surface (Figures S4&5), we designed fluorescently labelled barcodes A and B to further evaluate spatially confined binding of barcodes in tissue using fluorescence microscopy (Figure 1C, Figure S1C). We conjugated barcodes A(i=1-50) with fluorophore Cy3 and barcodes B(j=1-50) with fluorophore FITC. The first flow gave rise to stripes of Cy3 signal (red) corresponding to barcodes A immobilized by hybridization to mRNAs fixed in tissue. The second flow added barcodes B only to the regions where barcodes A were immobilized, yielding isolated square pixels of FITC signal (green) (Figure 1C). We also used a layer of NIH3T3 cells grown on a glass slide and fixed with formaldehyde to mimic a thin “tissue” section (Figure 1D), which had a higher surface roughness and served as a stringent model to evaluate the leakage across microchannels. Small molecule dyes DAPI (DNA stain, blue) and phalloidin (actin stain, red) with higher diffusivity as compared to DNA oligomers were used in the first and the second flow, respectively. When a microchannel wall cut through one cell or one nucleus, fluorescence signal was observed only in the half within the microchannel (Figure 1D and Figure S6). To evaluate the possibility of DNA diffusion through the tissue matrix underneath the microchannel wall, a 3D fluorescence confocal image was collected, which confirmed negligible leakage signal throughout the tissue section thickness (Figure 1E). A 10μm chip was used to perform the DBiT workflow with FITC-tagged barcodes B, which yielded a square lattice of green fluorescence pixels (Figure 1F and Figure S5). Interestingly, we found that compressing the PDMS microchip against the tissue section led to plastic deformation of the tissue underneath the microchannel walls, which allowed for imaging the topography of tissue pixels and compare to fluorescence. The decrease of fluorescence from the microchannel edge to half of the peak intensity was used to estimate the “diffusion” distance (Figure 1G). It was found to be 0.9±0.2μm for 10μm channels operated with a clamp and 4.5±1μm for 50μm channels without a clamp, which validated spatially confined delivery and binding of DNA barcodes to mRNAs in tissue using microfluidics.

### Evaluation of DBiT-seq data quality

The PCR amplicons were analyzed for cDNA size distribution that showed the peaks around 900-1100bp (Figure S7). We conducted paired-end (2×100) sequencing to identify spatial barcodes and the expression of mRNAs on each pixel. The reads were first processed using a custom python script to extract UMIs, barcodes A and barcodes B. The processed reads were demultiplexed, trimmed, and mapped against the mouse genome (GRCh38, Gencode M11 annotation) using the ST pipeline reported previously (Navarro et al., 2017). Total number of UMIs per pixel and detected genes were calculated (Figure 1H and Figure S8). In a 10μm pixel DBiT-seq experiment, we detected 22,969 genes in total and an average of 2,068 genes per pixel. In contrast, Slide-seq (Rodriques et al., 2019), which has the same pixel size (10μm), detected ∼150 genes per pixel(spot). This improvement in data quality allowed DBiT-seq to directly visualize the expression pattern of individual genes at cellular level. The number of UMIs or genes per pixel detected by the low-resolution Spatial Transcriptomics (ST) method (Stahl et al., 2016) was similar to that from DBiT-seq, but the spot size in ST was ∼100μm, two orders of magnitude larger in area. The commercialized Visium system (10X genomics) reduced the spot size to 55μm and the performance is still comparable to our 10 μm resolution chip. We also compared the saturation curves of DBiT-seq at different resolutions (10 μm and 50 μm) that were found to be nearly identical (Figure S9), demonstrating technical consistency and low variability. Compared with the ST method (Stahl et al., 2016), the saturation curves showed a similar trend but DBiT-seq was able to reach a higher number of total identified genes. There existed a detection bias dependent on the gene length similar to that observed in ST (Figure S10).

DBiT-seq was further validated with single molecule fluorescence in situ hybridization (smFISH) for a panel of specific genes (Trf, Ttn, Dlk1 and Strp2), which were measured on the same mouse embryo but different tissue sections (Figure S11A). The spatial expression patterns obtained by smFISH were similar to those from DBiT-seq. The differences in quantitative line profiles were mainly due to the fact that smFISH and DBiT-seq were conducted on different tissue sections, especially for Ttnm. Single molecule counts in smFISH were quantified and compared side-by-side to the transcript counts detected by DBiT-seq (Figure S11B). We estimated that DBiT-seq detected an average of ∼15.5% of total mRNA transcripts defined by smFISH, which was 1-2 orders of magnitude higher than Slide-seq.

### Spatial multi-omics mapping of whole mouse embryos

The dynamics of embryonic development and early organogenesis is intricately controlled spatiotemporally. A range of techniques such as FISH, immunohistochemistry (IHC), and RNAseq were used to yield a comprehensive database – eMouseAtlas data (Armit et al., 2017), which can be utilized to validate new spatial technology. We applied DBiT-seq to an E10 whole mouse embryo tissue slide at a pixel size of 50μm to computationally construct a spatial multi-omics atlas. The tissue histology image from an adjacent section was stained for Hemotoxylin and Eosin (H&E) (Figure 2A left). The spatial map of UMI counts (Figure 2A middle and right) showed co-detection of 12,314 UMIs for mRNAs and ∼3,038 UMIs for 22 proteins per pixel. It yielded an average of 4,170 genes detected per pixel. To benchmark DBiT-seq data, we aggregated the mRNA expression profiles of all pixels for each E10 embryo sample to generate “pseudo-bulk” data, which were compared to the “pseudo-bulk” data generated from scRNA-seq of E9.5 to E13.5 mouse embryos (Cao et al., 2019) using un-supervised clustering (Figure 2B). We observed consistent temporal developmental classification visualized in UMAP with four E10 DBiT-seq samples localized between E9.5 and E10.5 data from the reference (Cao et al., 2019). To further compare DBiT-seq to known spatial patterning during development, we examined a set of genes from the *Hox* gene family that plays an essential role in developmental specification along the anterior–posterior embryonic axis. *Hox1-3* genes are expressed throughout the neural tube, extending to the hindbrain, whereas *Hox8-9* are enriched in the lumbar and sacral (tail) region, (Deschamps and Duboule, 2017), which is consistent with our observations, for example, in *Hoxa2* vs *Hoxb9* (Figure S12). Unsupervised clustering of spatial pixel transcriptomes revealed eleven major clusters (Figure 2C), which correlated with telencephalon (forebrain), mesencephalon (midbrain), rhombencephalon (hindbrain), branchial arches, spinal neural tube, heart, limb bud, and ventral and dorsal side of main body for early internal organ development, based on Gene Ontology (GO) enrichment analysis (top ranked genes in Figure 2D). Using the literature database and the classical Kaufman’s Atlas of Mouse Development (Baldock and Armit, 2017), a manual anatomical annotation was performed to reveal 13 major tissue types(Figure 2E). Among them, 9 were consistent with the tissue types identified by GO analysis of DBiT-seq. Since a panel of 22 proteins were co-measured in this experiment, we attempted a head-to-head comparison between proteins and mRNAs (Figure S13). Notch signaling plays a crucial role in regulating a vast array of embryonic developmental processes. Notch1 protein was found highly expressed throughout the whole embryo, which is consistent with extensive *Notch1* mRNA expression. CD63 is an essential player in controlling cell development, growth, proliferation, and motility. The mRNA of CD63 was indeed expressed extensively in the whole embryo but a higher expression level in hindbrain and heart. Pan-EC-Antigen (PECA) or MECA-32, as a pan-endothelial marker, was expressed in multiple tissue regions containing microvasculature. The expression of EpCAM, a pan-epithelial marker, was localized in highly specific regions as seen for both mRNA and protein. Integrin subunit alpha 4 (ITGA4), known to be critical in epicardial development, was indeed highly expressed in heart but also observed in other tissue regions but the cognate protein was observed throughout the whole embryo. Certain genes, such as NPR1, showed strong discordance between mRNA and protein. A pan-leukocyte protein marker CD45 was observed in dorsal aorta and brain although the expression level of its cognate mRNA *Ptprc* was low. A chart was generated to show the expression of 8 mRNA/protein pairs in 13 anatomically annotated tissue regions (Figure 2F). We calculated the correlations across all 15 detected pairs (Figure S14A) and the average Pearson correlation coefficient (∼0.28) was low but as expected according to literature (de Sousa Abreu et al., 2009; Vogel and Marcotte, 2012) (Figure S14B). Next, to further validate the DBiT-seq protein expression, immunofluorescence staining was performed on the same embryo (different tissue sections) using antibodies of P2RY12 (microglia in central nerve system), PECA (endothelium), and EpCAM (epithelium). We observed a consistent pattern of EpCAM between immunostaining and DBiT-seq (Figure S15). It is worth noting that pan-mRNA UMI count map showed horizontal bands due to the variability in microfluidic flow, which was mechanistically similar to DNA barcode density variability on beads observed in scRNA-seq and normalization should be performed to correct this effect (Figure S16).

**Figure 2.**
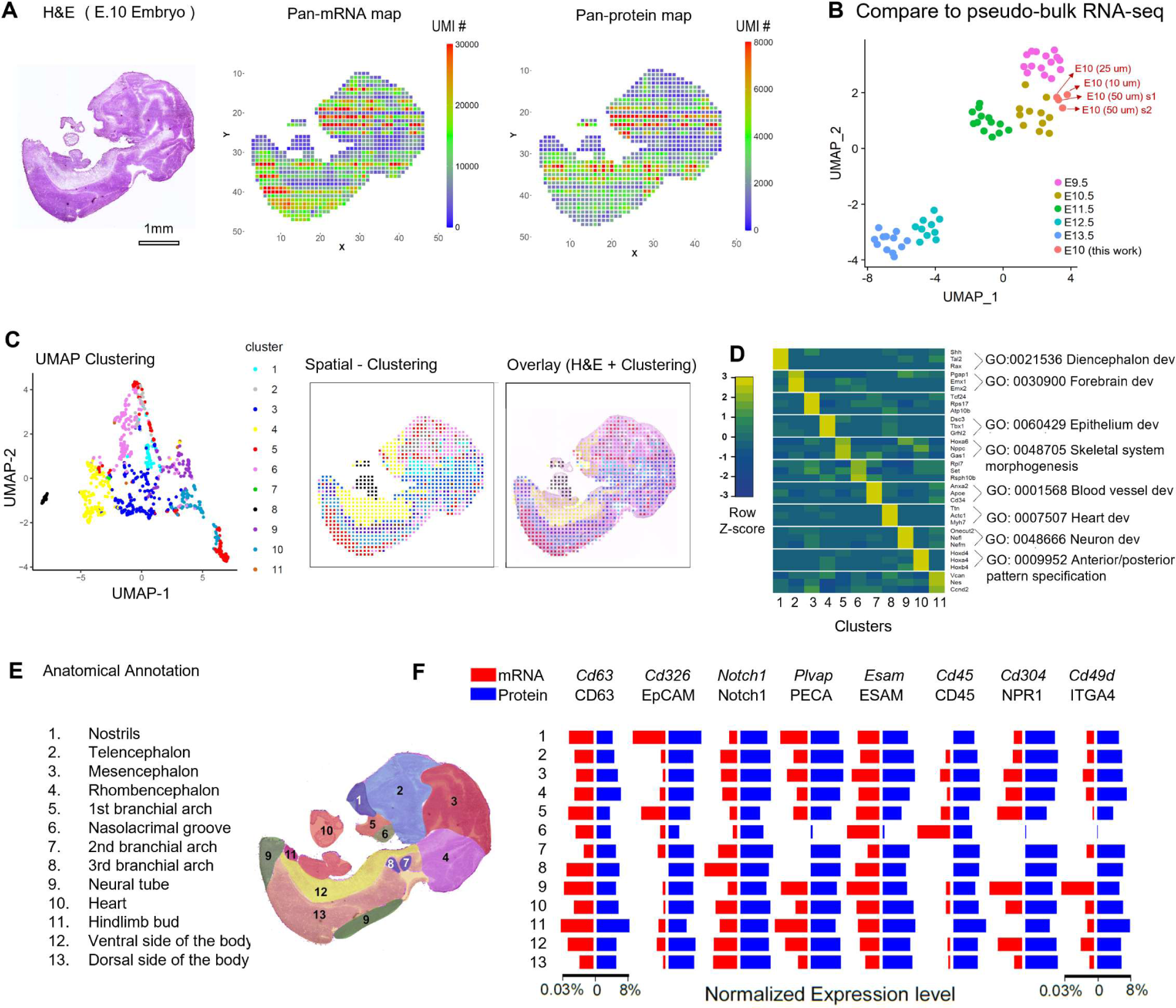
Spatial multi-omics mapping of whole mouse embryos. (A) Pan-mRNA and pan-protein panel spatial expression maps (pixel size 50μm) reconstructed from DBiT-seq, together with the H&E image of an adjacent tissue section. Whole transcriptome signal correlated with anatomic tissue morphology and density. (B) Comparison to “pseudo bulk” RNA-seq data. The aggregated transcriptome profiles of four E10 embryo samples sequenced using DBiT-seq conformed well to the correct stage (E10) of a single-cell RNA-seq data UMAP generated from mouse embryos ranging from E9.5-E13.5(Cao et al., 2019). (C) Unsupervised clustering of all pixels and mapping back to the spatial atlas. The results were shown in a UMAP plot and plotted back to a spatial map, which was further compared to the anatomical structure via overlay with the H&E image. Since the H&E stained tissue is not exactly the same as the slide we used for DBiT-seq, there are some minor inconsistencies observed between the H&E image and the DBiT-seq heatmap. (D) GO analysis of the 11 clusters identified. The results agree strongly with anatomical annotation. (E) Anatomic annotation of tissue (sub)types. To facilitate the examination of tissue morphology in correlation with the spatial gene expression atlas, an adjacent tissue section was stained with H&E and overlaid with the unsupervised clustering map. The clustering analysis revealed 11 major tissue (sub)types, which are in general agreement with anatomically annotated tissue regions (1-13). (F) Correlation between mRNAs and proteins in anatomically annotated tissue regions. The average expression levels of individual mRNAs and cognate proteins in each of the thirteen anatomically annotated tissue regions were compared.

### Spatial multi-omics mapping of an embryonic brain

We conducted a higher resolution (pixel size = 25μm) DBiT-seq to analyze the brain region of an E10 mouse embryo (Figure 3). As compared to the 50μm experiment (Figure 2), pan-mRNA and pan-protein UMI count maps (Figure 3C) showed finer structures that correlated with tissue morphology (Figure 3B). We surveyed all 22 individual proteins (Figure S17) and observed distinct expression patterns in at least 12 proteins with four shown in Figure 3D. CD63 was expressed extensively except in a portion of the forebrain. PECA, a pan-endothelial cell marker, was unambiguously detected in brain microvasculature, which was not readily distinguishable in tissue histology. EpCAM was localized in highly defined regions as thin as a single line of pixels (∼25μm) with high signal-to-noise ratio. MAdCAM was differentially expressed in a sub-region of the forebrain. To validate these observations, we performed immunofluorescence staining using nearby tissue sections from the same embryo to detect EpCAM and PECA. Spatial expression maps obtained by DBiT-seq and immunofluorescence staining were superimposed onto a H&E image and their line profiles were drawn for quantitative comparison (Figure 3E). The major peaks agreed with each other although some discordance in exact peak positions was observed because different tissue sections were used for DBiT-seq and immunofluorescence. Finally, we performed unsupervised clustering of all the pixels using their mRNA expression profiles and identified 10 distinct clusters, characterized by specific marker genes (Figure 3F). We then plotted the spatial distribution of pixels in four representative clusters against the H&E image (Figure 3G). Pathway analysis of marker genes revealed that cluster 1 was mainly involved in telencephalon development, cluster 2 associated with erythrocytes in blood vessels, clusters 3 implicated in axonogenesis, and clusters 4 corresponding to cardiac muscle development, in good agreement with anatomical annotations. Cluster 2, enriched for hemoglobulin genes in red blood cells, coincided with PECA protein expression that delineated endothelial microvasculature. We further demonstrated that high-quality spatial protein mapping data can be used to guide genome-wide spatial gene expression analysis.

**Figure 3.**
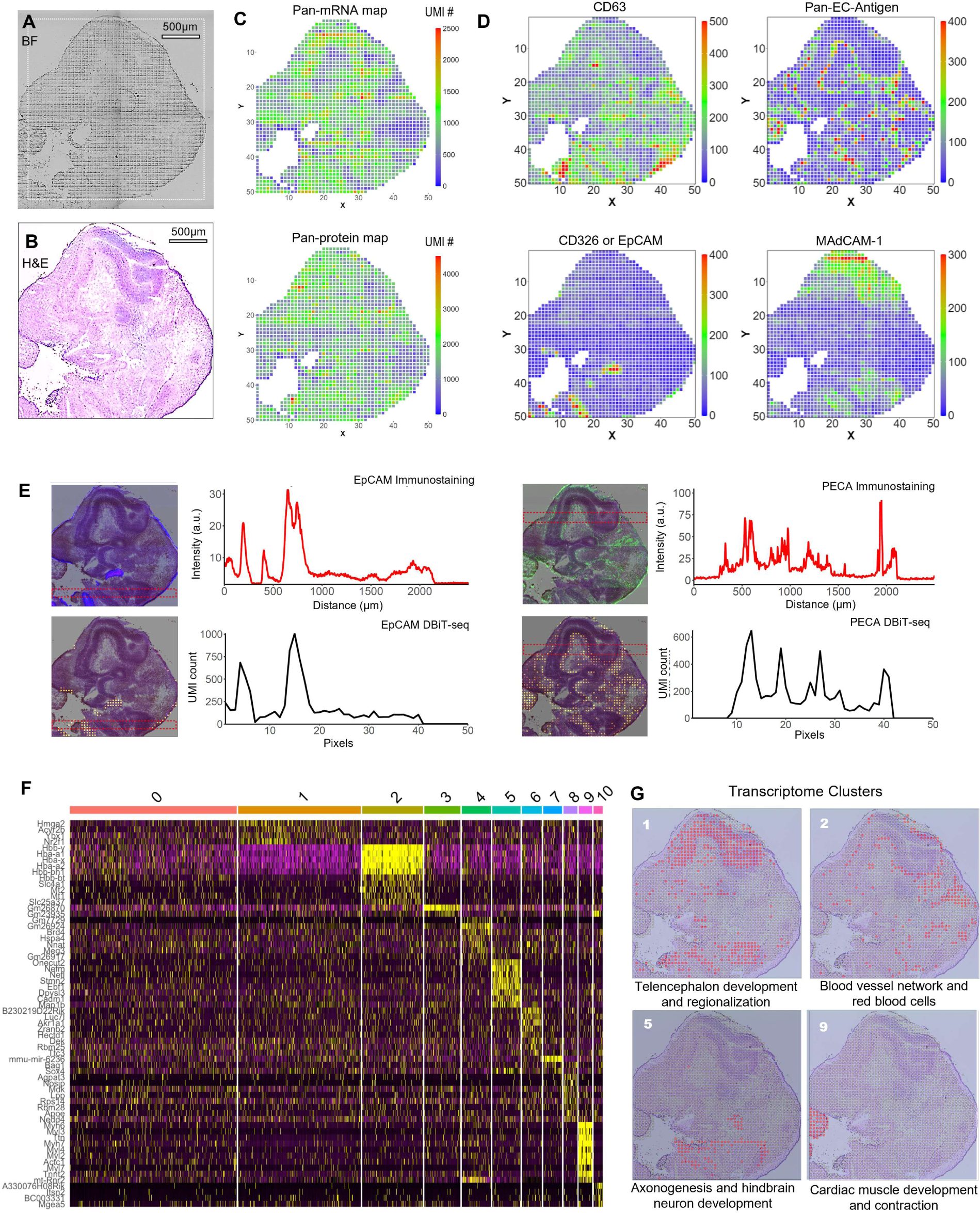
Spatial multi-omics mapping of an embryonic mouse brain. (A) Bright field image of mouse embryo E10 head region. (B) H&E stained mouse embryo E10 head region. H&E was performed on an adjacent tissue section. (C) Pan-mRNA and pan-protein-panel spatial expression maps of the brain region of a mouse embryo (E10) obtained at a higher resolution (pixel size 25μm) in comparison with the H&E image taken from an adjacent tissue section. The spatial pattern of whole transcriptome signal correlated with cell density and tissue morphology. (D) Spatial expression of four individual proteins: CD63, Pan-endothelial-Antigen, EpCAM (CD326) and MAdCAM-1. Spatial protein expression heatmaps explicitly showed brain region-specific expression and the network structure of brain microvasculature. (E) Validation by immunofluorescence staining. Spatial expression of EpCAM and PECA reconstructed from DBiT-seq and the immunofluorescence staining of the same proteins were superimposed on the H&E image for comparison. A highly localized expression pattern of EpCAM is in strong correlation with immunostaining as seen by the line profile. The network of microvasculature revealed by PECA in DBiT-seq is strongly correlated with the immunostaining signal. (F) Gene expression heatmap of the 10 clusters (showing top 4 genes for each cluster). (G) Spatial patterns of cluster 1, 2, 5 and 9. Go analysis identified the main biological process within each cluster, which matches with histological annotation.

### High-resolution mapping of early eye development

We conducted further spatial transcriptome mapping of the developing eye field in a E10 mouse embryo using 10μm microfluidic channels and the resultant pan-mRNA UMI heatmap (red) was superimposed onto the whole mouse embryo tissue image (Figure 4A). An enlarged view of the mapped region showed the imprinted morphology and individual pixels. An adjacent tissue section was stained for H&E (Figure 4B). At this stage (E10), the eye development likely reaches a late optic vesicle stage. Four genes were identified within the optic vesicle with distinct but spatially correlated expression patterns (Figure 4C). *Pax6* was expressed in the optic vesicle and stalk (Heavner and Pevny, 2012; Smith et al., 2009). *Pmel*, a pigment cell-specific gene(Kwon et al., 1991) involved in developing fibrillar sheets, was observed around the optic vesicle. Six6, a gene known for specification and proliferation of retinal cells in vertebrate embryos, was mainly localized within the optical vesicle but not the optic stalk (Heavner and Pevny, 2012). *Trpm1* lined the optic vesicle showing minimal overlap with Six6. It is known that the retinal pigment epithelium (RPE) consists of a single-cell-layer of melabocytes lining around an optic vesicle, which was successfully detected by DBiT-seq with markers like *Pmel* and *Trpm1* (Mort et al., 2015). We further performed GO analysis to identify major pathways and signature genes (Figure S20). Eye development and melanin pathways emerged as the two major categories. Additionally, we performed 10µm DBiT-seq on an E11 mouse embryo and compared it with E10 side-by-side for the eye field region (Figure 4D). The expression patterns of *Pmel, Pax6* and *Six6* around the eye were similar between E10 and E11 embryo, but showed spatial changes as the optic cup started to form in E11(Yun et al., 2009). Additionally, we analyzed other genes known to be involved in early eye formation (Figure 4E, 4F and 4G). *Aldh1a1*, a gene encoding Aldehyde Dehydrogenase 1 Family Member A1, was observed in the dorsal retina whereas *Aldh1a3* was mainly located at the ventral side and RPE. The spatial patterning of *Aldh1a1* and *Aldh1a3* within the eye field and the changes from E10 to E11 were in agreement with literature, showing that the *Aldh1a* family genes differentially control the dorsal-ventral polarization in embryonic eye development (Matt et al., 2005). We noticed that *Msx1*, a gene highly expressed in both ciliary muscle and ciliary epithelium as the structural support of eye (Zhao et al., 2002), was mainly surrounding the eye field in both E10 and E11 embryos. *Gata3*, a gene pivotal for eye closure, was enriched at the front end of the eye field to control the shape of eye during development. Our data allowed for high-spatial-resolution visualization of genome-wide gene expression in early stage eye field development.

**Figure 4.**
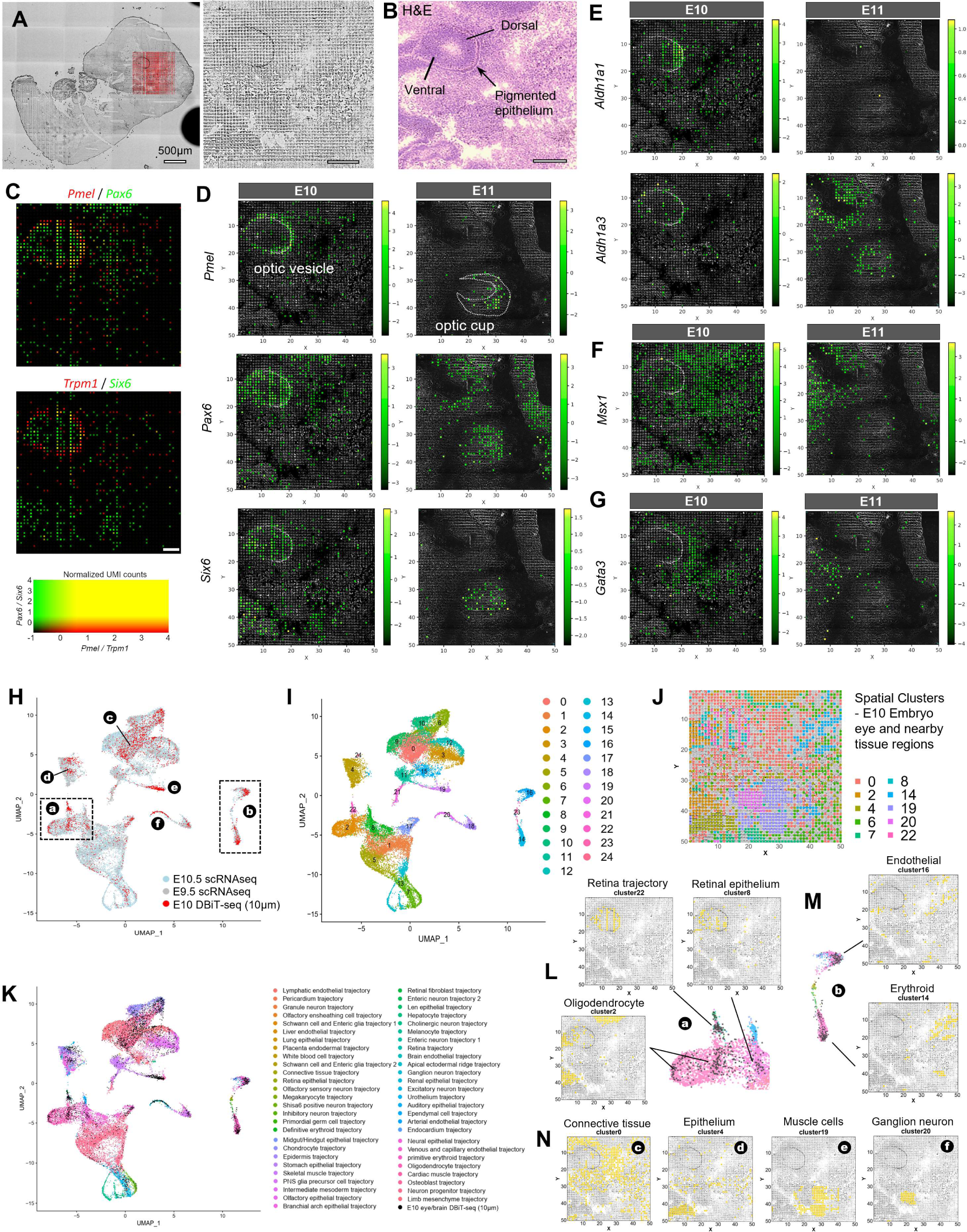
High-resolution mapping of early eye development. (A) Bright field image of whole mouse embryo E10 and enlarged region of interest, profiled by OBiT-seq (10 1-1m). (B) H&E stained mouse embryo E10 eye region. H&E was performed at an adjacent tissue section. (C) The overlay of selected genes revealed the relative spatial relationship at ultra-high resolution. For example, *Pax6* is expressed in whole optical vesicle including one layer of melanoblasts defined by *Pmel* and the optical nerve fiber bundle on the left. Six6 is expressed in the eye vesicle but does not overlap with the melanoblast layer even though they are in close spatial proximity to each other. (D) *Pmel*, *Pax6* and *Six6* spatial expression superimposed on the darkfield image of embryo E10 and E11 (pixel size 10μm). They are all associated with early stage embryonic eye development but mark different fine structures. *Pmel* was expressed in a layer of melanoblast cells lining the optical vesicle. *Pax6* and *Six6* were mainly expressed inside the optical vesicle but also seen in other regions mapped in this data. (E) Spatial expression pattern of gene *Aldh1a1* and *Aldh1a3*. *Aldh1a1* is expressed in dorsal retina of early embryo, and meanwhile, Adlh1a3 is mainly expressed in melanoblast cells and in ventral retina. (F) Spatial expression pattern of gene *Msx1*. *Msx1* is mainly enriched in ciliary body of the eye, including the ciliary muscle, and the ciliary epithelium, which produces the aqueous humor. (G) Spatial expression pattern of gene *Gata3*. *Gata3* is essential for lens development and mainly expressed in posterior lens fiber cells during embryogenesis. (H) Integration of scRNA-seq data (Cao et al., 2019) with DBiT-seq (10 µm) data. scRNA-seq data was integrated with DBiT-seq data, and UMAP clustering showed a strong agreement between these two sets of data. (I) Clustering of the combined datasets of scRNA-seq and DBiT-seq. (J) Clustering of DBiT-seq data alone using Seurat package. (K) UMAP plot with pixels colored by cell type. (L), (M) & (N) Cell type annotation of DBiT-seq data using scRNA-seq reference data. Totally 9 cell types were identified in this region.

### Direct integration with single-cell RNA sequencing data

We observed additional tissue features based on the spatial expression pattern of 19 top ranked genes (Figure S21) but the cell types could not be readily identified. Since the pixel size (10µm) in this experiment was approaching cellular level, we speculated that it is possible to directly integrate data from scRNA-seq and DBiT-seq to infer cell types and visualize spatial distribution. scRNA-seq data from E9.5 and E10.5 mouse embryos (Cao et al., 2019) were combined with high-resolution DBiT-seq (10µm) data from an E10 mouse embryo to perform unsupervised clustering (Figure 4H). We found that the spatial pixels (red) conformed well into single cell transcriptomes (blue and gray) and together identified 24 clusters in the combined dataset (Figure 4I). Each cluster was mapped back to its spatial distribution in tissue (8 clusters are shown in Figure 4J). We further used scRNA-seq data as a reference for cell type annotation (Figure 4K) and the reported 53 cell types were directly compared to DBiT-seq data (black) in UMAP, allowing for detecting the dominant cell type in each pixel (10µm). Then, we could link scRNA-seq-annotated cell types to corresponding spatial pixels and visualize cell type distribution on the tissue. First, we examined spatial pixels in clusters 2, 8 and 22 (see a in Figure 4H) and the dominant cell types were found to be retina trajectory, retina epithelium, and oligodendrocyte. Mapping cell type-annotated pixels to the tissue image showed that retina trajectory and retina epithelium cells were indeed localized within the optic vesicle while oligodendrocytes were localized in three tissue regions with one corresponding to optic stalk right next to optic vesicle, in agreement with the observation that multiple sub-clusters of oligodendrocyte pixels were present(Figure 4L). Second, spatial pixels in the region b of Figure 4H were detected only in clusters 14 and 16, which were found to be dominated by erythroid and endothelial cells. Mapping them back to the tissue image revealed microvessels (endothelial) and blood clots (erythroid) at the upper right corner (Figure 4M). Third, we also analyzed spatial pixels in c-f of Figure 4H and the corresponding clusters 0, 4, 19, and 20, respectively. Linking spatial pixels to cell types revealed (c) connective tissues as the structural support of eye formation, (d) epithelial cells forming the pituitary gland, muscle cells (e) surrounding the trigeminal sensory nerve for facial touch sensing, and ganglion neurons (f) in the trigeminal sensor itself(Figure 4N). Thus, high-resolution (10µm) DBiT-seq data can be directly integrated with scRNA-seq to infer cell types and visualize spatial distribution in the tissue context.

### Clustering analysis of 11 embryo samples across different stages (E10-12)

To further understand the early development of mouse embryo over time, we integrated the DBiT-seq data of 11 mouse embryo tissue samples from three stages, E10, E11 and E12 (Figure 5, detailed information in Table S4) and conducted unsupervised clustering, which showed 20 clusters visualized by t-distributed stochastic neighbor embedding (t-SNE) (Figure 5A&5B) and the top differentially expressed genes (Figure 5C). Cluster 2 was associated with muscle system processes with the *Myl* gene family preferentially expressed and the pixels in this cluster were mainly from three E11 tail samples (see blue in Figure 5A). Although the pixels from the same sample were clustered together without batch normalization, some samples like “E11 Tail (25µm) 1” showed multiple distant clusters (Figure 5D left panel), indicating significant difference of tissue types in this sample. The large pixels (50µm) tend to locate away from the origin of the UMAP presumably because they covered many more cells and possessed a higher degree of cell diversity within a pixel. In contrast, the 10µm pixels were clustered around the center of the UMAP, indicating a convergence to single-cell-level gene expression. E10, E11 and E12 pixels were spaced out along the same trajectory (left to right) consistent with the development stages although these samples were hugely different, so that they were mapped for different tissue regions (head vs tail) and at different resolutions (pixel size 10, 25 vs 50 µm) (Figure 5D right panel).

**Figure 5.**
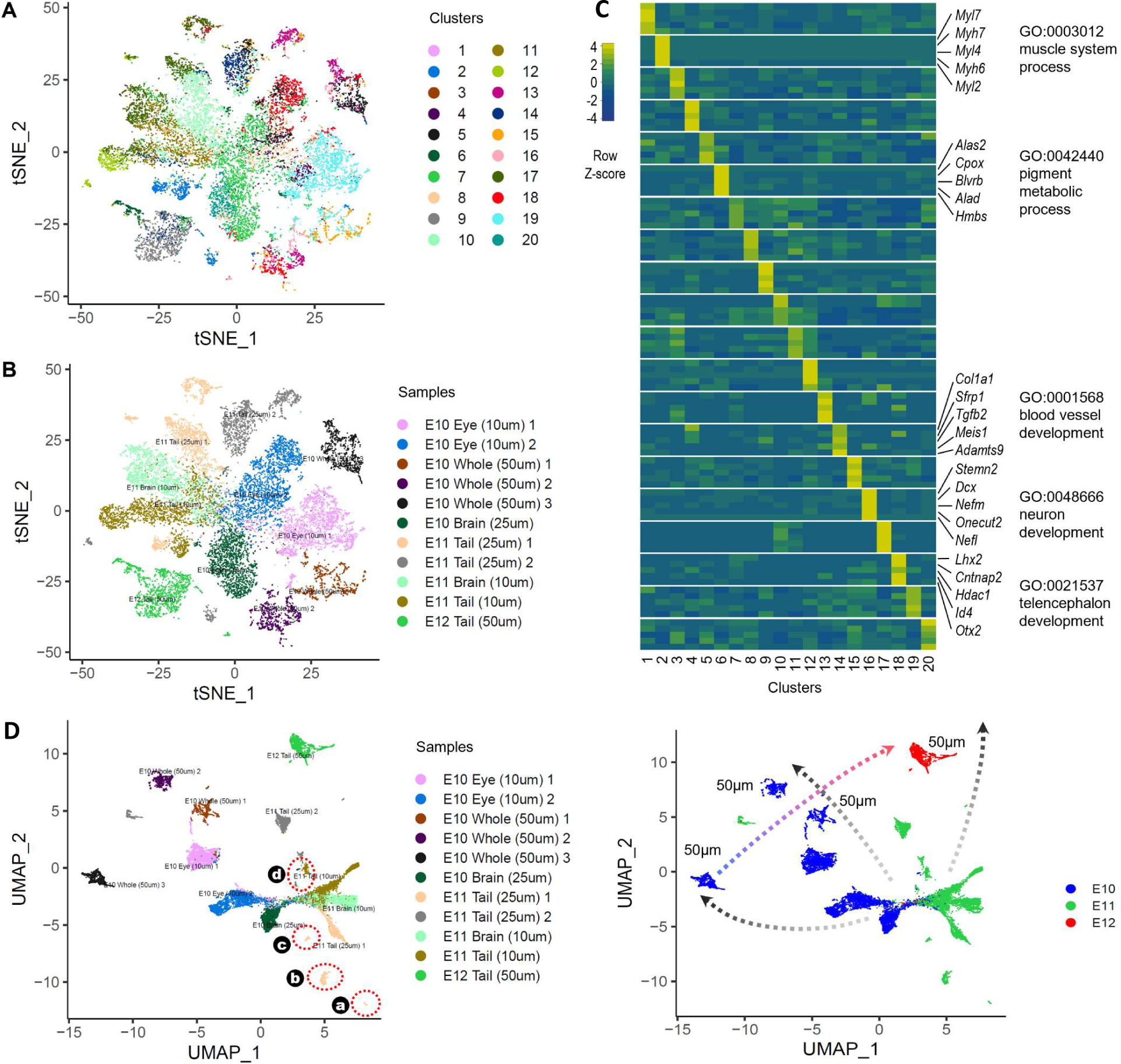
Global clustering analysis of 11 mouse embryos from E10, E11 to E12. (A) TSNE plot of integrated DBiT-seq datasets. 11 different samples were integrated and plotted within one TSNE plot. There are a total of 20 clusters identified with optimized clustering parameter settings. (B) TSNE plot labelled with sample names. (C) GO analysis and differentially expressed genes of all 20 clusters. Some of the interesting clusters are muscle system, pigment metabolic system, blood vessel development, neuron development and telencephalon development. (D) UMAP plot of integrated DBiT-seq datasets. Plots were labelled with sample names (left) or embryonic age (right).

### Spatial mapping of internal organ development

Sample “E11 Tail (25µm) 1” showed multiple distinct sub-clusters in the global UMAP (Figure 5D left panel) which made us wonder what cell types constitute these clusters (see enlarged view in Figure 6A). Four subclusters (a, b, c and d) were mapped back to the tissue image, which revealed distinct spatial patterns for all of them (Figure 6B). Clustering analysis of all pixels in this sample identified 13 clusters visualized in both UMAP (Figure 6C) and spatial map (Figure 6D). To unveil the identities of these spatial patterns, we again use scRNA-seq as reference(Cao et al., 2019) to perform automated cell type annotations (Figure 6E) with SingleR (Aran et al., 2019). The dominant cell types in these spatial clusters (a, b, c, and d) were associated with different internal organs such as liver (cluster a), neutral tube (cluster b), heart (cluster c), and blood vessels containing coagulated erythrocytes (cluster d) (Figure 6G). We further visualized the spatial expression of 8 representative marker genes (Figure 6F). *Myh6*, a gene encoding Myosin heavy chain α, was highly expressed in atria, while *Myh7* (encoding myosin heavy chain β) was the predominant isoform expressed in ventricular muscle, allowing for not only detecting cardiac muscle cells but also differentiating between atria vs ventricle of an embryonic heart. *Pax6* was expressed in region-specific neural progenitors in the neural tube. *Car3*, which encodes carbonic anhydrase III and expressed in slow twitch skeletal muscles, specifically delineated the formation of notochord. Apoa2, which encodes apolipoprotein E, is liver specific. Hemoglobin α encoding gene, *Hba.a2*, normally found in red blood cells, indicated the coagulated erythrocytes in both large vessels like dorsal aorta and microvessels in multiple organs. It was also found in the blood clots inside atria. *Col4a1*, which encodes a specific collagen, the type IV alpha1, produced by endothelial cells to form the basement membrane, precisely lined the inner surface of the dorsal aorta, which supposedly consisted of a single layer of endothelial cells. It was also expressed in heart presumably at endocardium and coronary arties. *Actb*, which encodes β-actin, a widely used reference or housekeeping gene, was expressed extensively throughout the embryo but showed lower expression in, for example, nervous tissues. We also compiled the “pseudo bulk” expression data by aggregating pixels in three major organs (heart, liver and neutral tube) and compared with the ENCODE bulk RNA-seq data side-by-side, which revealed excellent concordance (Pearson Correlation Coefficient = ∼0.8) (Figure S22).

**Figure 6.**
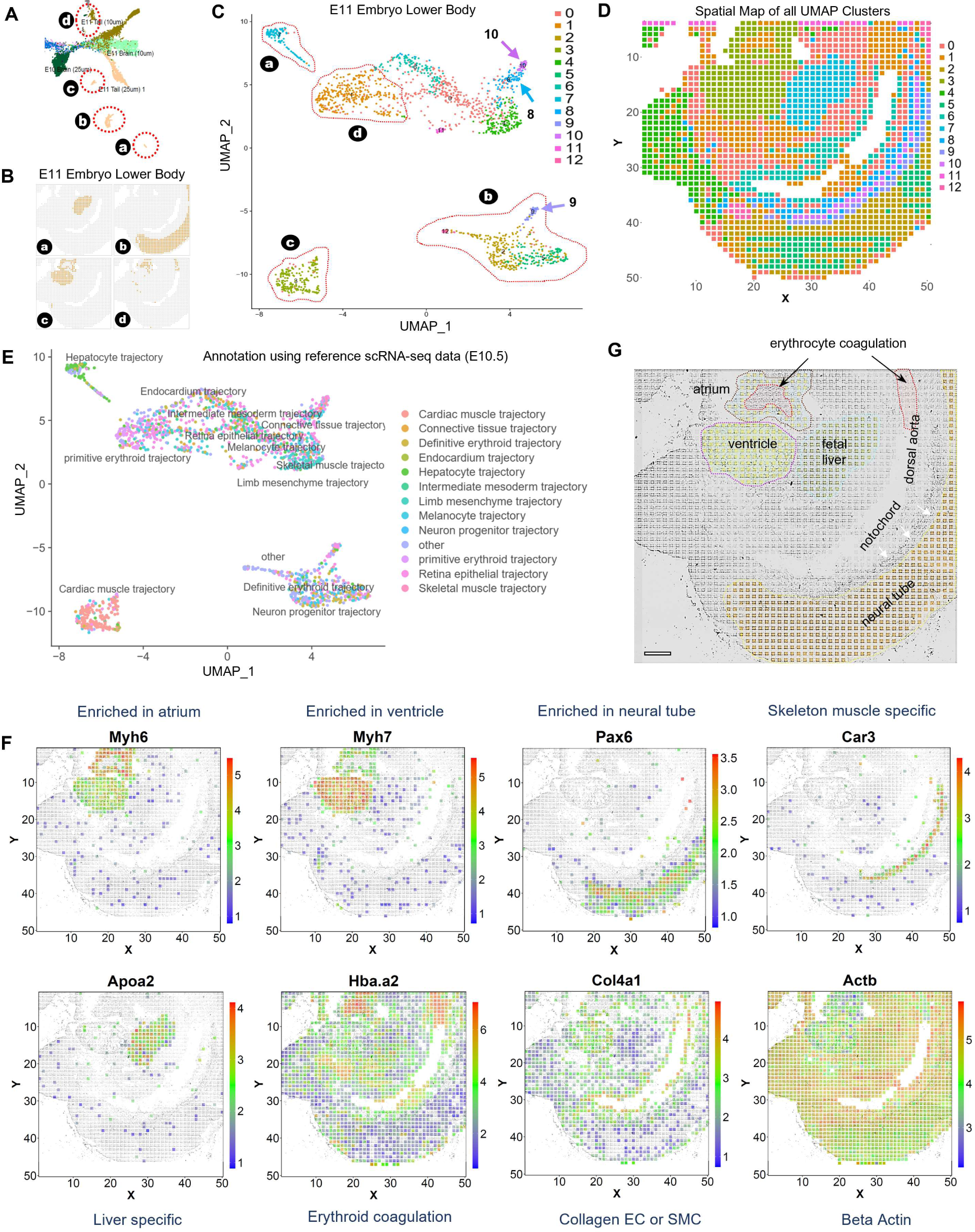
Mapping internal organs in a E11 mouse embryo. (A) Enlarged view of UMAP clustering of Figure 4A with a focus on E11 embryo lower body sample. (B) Mapping the four labelled clusters in the Figure 6A to their original tissue location. Each of the four clusters can be mapped back to a distinct organ. Cluster a is mapped to liver, cluster b to neutral tube region, cluster c to heart and cluster d to erythrocyte coagulates. (C) UMAP clustering of E11 embryo lower body sample. The four distinct clusters shown in Figure 6A were also labelled. (D) Spatial map of all the UMAP clusters of E11 embryo lower body sample. (E) Automatic cell annotation using SingleR package with reference scRNA-seq data of E10.5 embryo (Cao et al., 2019). (F) Spatial mapping of individual genes. (G) Anatomical annotations of a mouse embryo. Major organs including heart, liver and neutral tube are quite clear, whereas the erythrocyte coagulation can only be identified with the assistance of DBiT-seq data.

**Figure 7.**
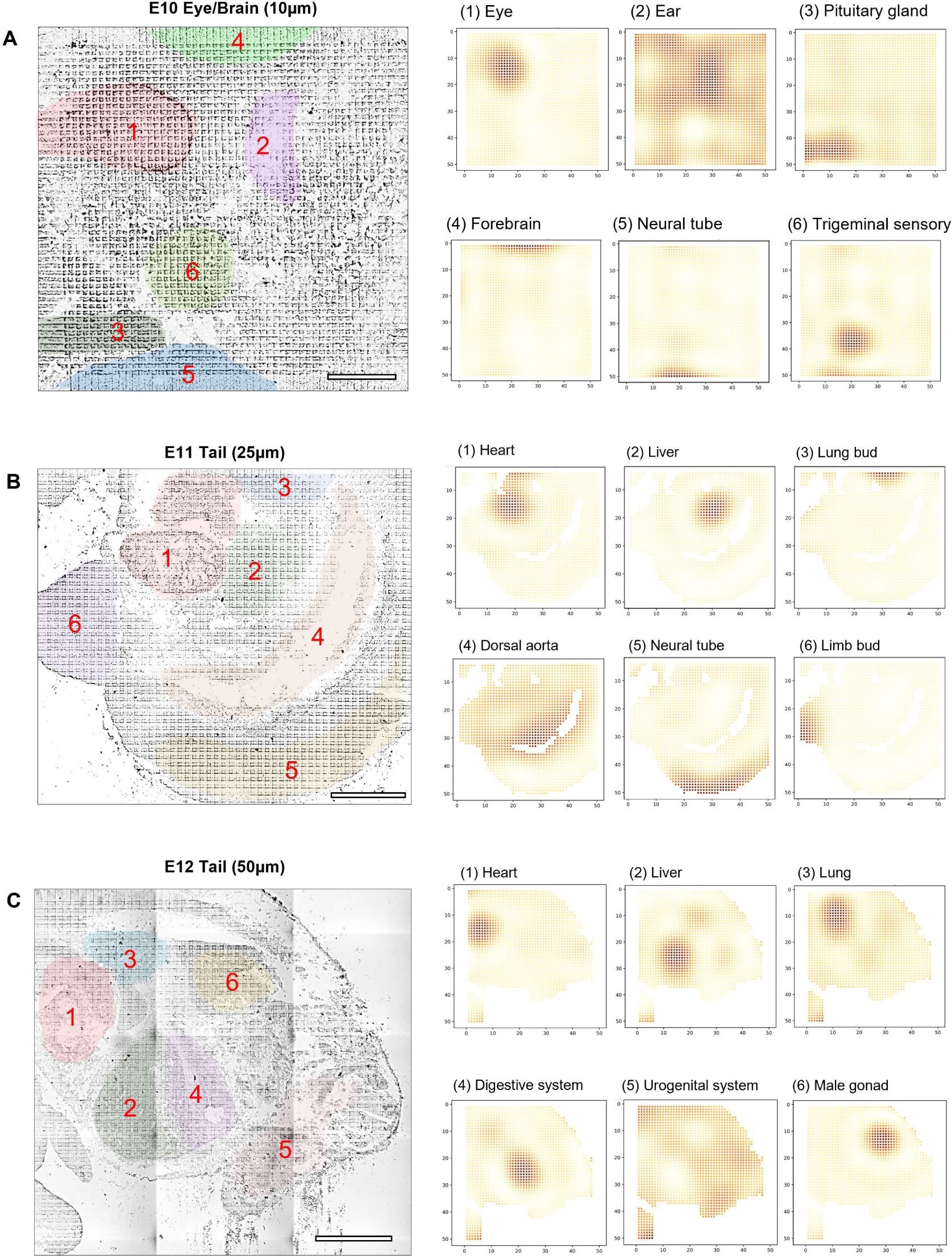
SpatialDE for automated feature identification. (A) Major features identified in the ultra-high-spatial-resolution data described in Figure 4. It revealed not only eye but numerous other tissue subtypes. (B) Major features identified in the lower body of a later-stage mouse embryo (Figure 6), which showed a wide variety of tissue types developed. (C) Major organs identified in the lower body of a E12 mouse embryo. It showed different tissue types that appeared at **this embryonic age.**

### Automated feature identification using spatialDE

Spatial differential expression (spatialDE) pipeline (Svensson et al., 2018a) previously developed for low-resolution spatial transcriptomics was evaluated in our study for automated discovery of spatial tissue features without using scRNA-seq for cell type annotation. In addition to the major pathways associated with eye development in Figure 4, spatialDE identified 20 features (see Figure S23 and Figure 7A) including eye, ear, muscle, forebrain, and epithelium, which are in agreement with scRNA-seq based cell type identification. In contrast, some features were hardly distinguishable in the corresponding tissue image such as ear (presumably due to too early stage in the developmental process) and forebrain (barely covered in the mapped tissue region). SpatialDE was applied to the data in Figure 6 and detected not only heart, liver, dorsal aorta, and neural tube as previously discussed but also a small fraction of lung bud covered in the mapped tissue region. Many internal organs begin to develop at the stage of E10 but barely distinguishable. To further evaluate the potential for spatialDE to detect more distinct organs or tissues, an E12 mouse embryo was analyzed using DBiT-seq. Interestingly, in only 1/3 of the whole embryo tissue section, spatialDE identified 40 distinct features including heart, lung, urogenital system, digestive system, and male gonad (testis) (see Figure S24 and Figure 6C). Many of these features were still too early to identify based on tissue morphology. We also revisited the E10 whole mouse embryo (Figure 2) and E11 lower body DBiT-seq data (Figure 6), and identified ∼20 and ∼25 distinct features, respectively (Figure S25&S26), which were less than that from the E12 sample, indicating that the features newly identified in E12 were associated with the developmental process and the emergence of internal organs at this stage.

### Combing immunofluorescence staining and DBiT-seq on the same tissue section

Lastly, we demonstrated DBiT-seq with immunofluorescence stained tissue sections. A E11 mouse embryo tissue slide was stained with DAPI (blue), phalloidin (Green) and red fluorescent labelled P2RY12 antibody (a G protein-coupled receptor) (Figure S27A). Then, we performed DBiT-seq. When the microfluidic chip was still on the tissue slide, we imaged the microfluidic channels and the tissue immunofluorescence. With DAPI staining for nucleus, we could conduct cell segmentation using ImageJ (Figure S27E). The immunostaining also enabled us to select the pixels of interest such as those containing single cells or those showing specific protein expression to study the association between morphological characteristics, protein expression, and transcriptome (Figure S27G&H). Immunofluorescence staining is widely used in tissue pathology to measure spatial protein expression at the cellular or sub-cellular level. Combining immunofluorescence with high-resolution (10µm) DBiT-seq on the same tissue slide could improve the mapping of spatial omics data to specific cell types.

## DISCUSSION

We developed a completely new technology for high-resolution (∼10um) spatial omics sequencing. Early attempts towards spatial transcriptomics were based on multiplexed fluorescent *in situ* hybridization(Chen et al., 2015; Eng et al., 2019; Lubeck et al., 2014; Perkel, 2019). Recently, a major breakthrough in the field arose from the use of high throughput NGS sequencing to reconstruct spatial transcriptome maps (Rodriques et al., 2019; Stahl et al., 2016), which were unbiased, genome-wide, and presumably easier to adopt by a wider range of scientists in the biological and biomedical research community. These NGS-based methods achieved spatial transcriptomics through a method called “barcoded solid-phase RNA capture”(Trcek et al., 2017), which used a DNA barcode spot array such as ST (Stahl et al., 2016) or a barcoded bead array such as Slide-seq (Rodriques et al., 2019) to capture mRNAs released from a freshly sectioned frozen tissue specimen carefully placed on top. These approaches are still technically demanding, requiring a lengthy and sophisticated step to decode the beads, while the mRNA capture efficiency and the number of detectable genes per pixel at the 10μm spot size level is markedly below optimal. Additionally, it is not obvious how to extend to other omics or multi-omics measurements. Herein, DBiT-seq is a fundamentally different approach which does not require the lysis of tissues to release mRNAs and is compatible with existing formaldehyde-fixed tissue slides. It obviates the need to conduct sophisticated sequential hybridization or SOLiD sequencing to decode beads. It is versatile and easy to operate with a simple PDMS slab clamped on the tissue slide and just a set of reagents. This standalone device is intuitive to use, requires no sophisticated fluidic handling, and thus can be readily adopted by researchers who have no training in microfluidics.

The versatility of our workflow further enabled combined spatial mapping of multiple omics such as whole mRNA transcriptome and a panel of 22 protein markers. It was applied to the study of whole mouse embryos and identified all major tissue types during early organogenesis. High-resolution DBiT-seq can readily resolve fine features such as brain microvasculature and a single-cell-layer of pigmentated epithelium lining around an optic vesicle. We demonstrated not only high spatial resolution but also excellent data quality with high genome coverage and large numbers of detectable genes per 10μm pixel as compared to Slide-seq or HDST. This improvement enabled us to visualize the spatial expression of individual marker genes to resolve fine features as thin as one cell layer. Integration of DBiT-seq and scRNA-seq data can readily identify the dominant cell type in each spatial pixel. We also demonstrated DBiT-seq on immunostained tissue slide, allowing for correlating cell morphology and spatial transcriptome at the cellular level.

Like any other emerging technologies, DBiT-seq has limitations. First, although it is close to single-cell level spatial mapping, it does not directly resolve single cells. Our approach potentially allows for high resolution immunofluorescence or FISH on the same tissue slide to facilitate cell segmentation and the deconvolution of spatial omics data to computationally derive single-cell spatial omics. Secondly, there is a theoretical resolution limit. Although the validation experiments in Figure 2 indicate the diffusion distance is ∼1μm and the theoretical resolution can be as fine as ∼2μm. However, because the tissue section thickness is >5μm and the tissue deformation may block the microchannel flow if the channels are small and shallow. According to our observations, the achievable smallest pixel size is approximately 5μm, in which most pixels contain one or a fraction of a cell, making computational convolution more feasible. Third, the number of flow channels in current DBiT-seq device is still limited such that the mappable area at the resolution of 10μm pixel size is 1mmx1mm. It can be increased by increasing the number of barcodes to 100×100 or 200×200. Alternatively, a serpentine microfluidic channel design can increase the mapping area without the need to increase the number of DNA barcodes.

In summary, we report on a versatile technology, microfluidic deterministic barcoding in tissue for spatial omics sequencing (DBiT-seq), to measure mRNA transcriptome and a panel of 22 proteins on a fixed tissue slide and at high spatial resolution (10μm pixel size). This NGS-based approach is unbiased and genome wide for mapping biomolecules in the tissue context. DBiT-seq differs fundamentally from other NGS-based spatial transcriptomics methods and only requires a set of reagents and a simple device to perform the experiments. The workflow is versatile and can be modified for the mapping of other biomolecular information. It may find applications in a wide range of fields in biological and biomedical research including developmental biology, neuroscience, cancer, immunobiology, and clinical pathology.

## SUPPLEMENTAL INFORMATION

Supplementary Information can be found online at [to be inserted, SI is also provided as part of the manuscript submission].

## ACKNOWLEDGEMENTS

We thank Drs. Haifan Lin, Andre Levchenko, Dianqing Wu and Jun Lu for helpful discussions. R.F. dedicates this work to his senior colleagues/mentors Drs. W. Mark Saltzman and Jay Humphrey on the occasions of their 60^th^ birthday celebration. This research was supported by Packard Fellowship for Science and Engineering (to R.F.), Stand-Up-to-Cancer (SU2C) Convergence 2.0 Award (to R.F.), and Yale Stem Cell Center Chen Innovation Award (to R.F.). Y.L. was supported by the Society for ImmunoTherapy of Cancer (SITC) Postdoctoral Fellowship. The molds for microfluidic chips were fabricated at the Yale University School of Engineering and Applied Science (SEAS) Nanofabrication Center. We used the service provided by the Genomics Core of Yale Cooperative Center of Excellence in Hematology (U54DK106857). This sequencing service was conducted at Yale Stem Cell Center Genomics Core Facility which was supported by the Connecticut Regenerative Medicine Research Fund and the Li Ka Shing Foundation. It was also conducted using the sequencing facility at the Yale Center for Genomic Analysis (YCGA).

## AUTHOR CONTRIBUTIONS

Conceptualization: R.F.; Methodology, Y.L., M.Y., Y.D., and G.S.; Experimental investigation, Y.L., Y.D., G.S., C.C.G., D.K., Z.B., Y.X., and D.Z.; Data Analysis, M.Y., Y.L., Y.D. and R.F.; Resources, D.K., Z.B., and Y.X.; Writing – Original Draft, R.F., Y.L., and M.Y.; Writing – Review and Editing, R.F., Y.L., M.Y., G.S., C.C.G.,T.T., A.E., and S.H.

## STAR METHODS

### KEY RESOURCES TABLE

| REAGENT or RESOURCE | SOURCE | IDENTIFIER |
| --- | --- | --- |
| Antibodies |  |  |
| Alexa Fluor® 647 anti-mouse CD326 (Ep-CAM) Antibody | Biolegend | 118212 |
| Alexa Fluor® 488 anti-mouse Panendothelial Cell Antigen Antibody | Biolegend | 120506 |
| PE anti-P2RY12 Antibody | Biolegend | 848004 |
| TotalSeq antibodies | Biolegend | See table S1 |
| Biological Samples |  |  |
| Mouse C57 Embryo Sagittal Frozen Sections, E10 | Zyagen | MF-104-10-C57 |
| Mouse C57 Embryo Sagittal Frozen Sections, E12 | Zyagen | MF-104-12-C57 |
| Chemicals, Peptides, and Recombinant Proteins |  |  |
| Maxima H Minus Reverse Transcriptase (200 U/L) | Thermo Fisher Scientific | EP0751 |
| dNTP mix | Thermo Fisher Scientific | R0192 |
| RNase Inhibitor | Enzymatics | Y9240L |
| SUPERase• In™ RNase Inhibitor | Thermo Fisher Scientific | AM2694 |
| T4 DNA Ligase | New England Biolabs | M0202L |
| Ampure XP beads | Beckman Coulter | A63880 |
| Dynabeads MyOne C1 | Thermo Fisher Scientific | 65001 |
| Proteinase K, recombinant, PCR grade | Thermo Fisher Scientific | EO0491 |
| Kapa Hotstart HiFi ReadyMix | Kapa Biosystems | KK2601 |
| Formaldehyde solution | Sigma | F8775-25ML |
| NEBuffer 3.1 | New England Biolabs | B7203S |
| T4 DNA Ligase Reaction Buffer | New England Biolabs | B0202S |
| PMSF | Sigma | 10837091001 |
| Evagreen Dye, 20X in water | Biotium | 31000-T |
| Critical Commercial Assays |  |  |
| Nextera XT DNA Preparation Kit | FC-131-1024 | Illumina |
| Deposited Data |  |  |
| Oligonucleotides |  |  |
| Primers, Ligation linkers, DNA barcodes | IDT | See Table S2 |
| Software and Algorithms |  |  |
| Other |  |  |

### LEAD CONTACT AND MATERIALS AVAILABILITY

Further information and requests for resources and reagents may be directed to the corresponding author Rong Fan.

### EXPERIMENTAL MODEL AND SUBJECT DETAILS

#### Animals

Mouse: C57BL/6NCrl (Charles River Laboratories)

This mouse is purchased from Charles River Laboratories, and the embryo was collected the day the mouse was received. For detailed information about mouse maintenance and care, refer to the manufacturer’s information sheet.

### METHOD DETAILS

#### Microfluidic device fabrication and assembly

The microfluidic device was fabricated with polydimethylsiloxane (PDMS) using soft lithography. The chrome photomasks with 10 µm, 25µm and 50 µm resolution were ordered from the company Front Range Photomasks (Lake Havasu City, AZ). The molds were fabricated using SU-8 negative photoresist according to the following microfabrication process. A thin layer of SU-8 resist (SU-8 2010, SU-8 2025 and SU-8 2050, Microchem) was spin-coated on a clean silicon wafer following manufacturer’s guidelines. The thickness of the resistant was ∼50 µm for the 50-µm-wide microfluidic channel device, ∼28 µm for 25-µm-wide device, and ∼20 µm for 10-µm-wide device. A protocol to perform SU-8 photo lithography, development, and hard baking was followed based on the manufacturer’s (MicroChem) recommendations to yield the silicon molds for PDMS replication.

PDMS microfluidic chips were then fabricated via a replication molding process. The PDMS precursor was prepared by combining GE RTV PDMS part A and part B at a 10:1 ratio. After stir mixing, degassing, this mixture was poured to the mold described above, degassed again for 30min, and cured at 75 °C for ∼2 hours or overnight. The solidified PDMS slab was cut out, peeled off, and the inlet and outlet holes were punched to complete the fabrication. The inlet holes were ∼2 mm in diameter, which can hold up to 13 µL of solution. A pair of microfluidic chips with the same location of inlets and outlets but orthogonal microfluidic channels in the center were fabricated as a complete set of devices for flow barcoding a tissue slide. To do that, the PDMS slab was attached to the tissue section glass slides and a custom-designed acrylic clamp was used to firmly hold the PDMS against the tissue specimen to prevent leakage across microfluidic channels without the need for harsh bonding processed such as thermal bonding or plasma bonding(Temiz et al., 2015).

#### DNA barcodes and other key reagents

Oligos used were listed in Table S1 Antibody-Oligo sequences and Table S2 DNA oligos and DNA barcodes. All other key reagents used were listed as Table S3.

#### Tissue Handling

Formaldehyde fixed tissue or frozen tissue slides were obtained from a commercial source Zyagen (San Diego, CA). The protocol Zyagen used to prepare the embryonic tissue slides is the following. The pregnant mice (C57BL/6NCrl) were bred and maintained by Charles River Laboratories. More information can be found in the information sheet. The time-pregnant mice (day 10 or day 12) were shipped to Zyagen (San Diego, CA) the same day. The mice were sacrificed at the day of arrival for embryos collection. The embryo sagittal frozen sections were prepared by Zyagen (San Diego, CA) as following: the freshly dissected embryos were immersed into OCT and snapped frozen with liquid nitrogen. Before sectioning, the frozen tissue block was warmed to the temperature of cryotome cryostat (-20°C). Tissue block was then sectioned into thickness of ∼7 µm and placed in the center of a poly-L-lysine coated glass slide (CatLog no. 63478-AS, electron microscopy sciences). The frozen slides were then fixed with 4% formaldehyde or directly kept at -80 °C if a long-time storage is needed.

#### Tissue slides and fixation

To thaw the tissue slides, they were taken out of the freezer, placed on a bench at room temperature for 10 minutes, and then cleaned with 1X phosphate buffer saline (PBS) supplemented with RNase inhibitor (0.05U/μL, Enzymatics). If the tissue slides were frozen sections, they were first fixed by immersing in 4% formaldehyde (Sigma) for 20 minutes. Afterwards, the tissue slides were dried with forced nitrogen air and then ready to use for spatial barcoding.

#### Tissue histology and H&E staining

An adjacent tissue section was also requested from the same commercial resource which could be used to perform tissue histology examination using H&E staining. Basically, the fixed tissue slide was first cleaned by DI water, and the nuclei were stained with the alum hematoxylin (Sigma) for 2 minutes. Afterwards, the slides were cleaned in DI water again and incubated in a bluing reagent resolution (0.3% acid alcohol, Sigma) for 45 seconds at room temperature. Finally, the slides were stained with eosin for 2 more minutes. The stained embryo slide was examined immediately or stored at -80 °C fridge for future analysis.

#### Immunofluorescence staining

Immunofluorescence staining was performed either on the same tissue slide or an adjacent slide to yield validation data. Three fluorescent-labelled antibodies listed below were used for visualizing the expression of three target proteins: Alexa Fluor 647 anti-mouse CD326 (Ep-CAM) Antibody, Alexa Fluor 488 anti-mouse Panendothelial Cell Antigen Antibody, PE anti-P2RY12 Antibody. The procedure to stain the mouse embryo tissue slide is as follows. (1) Fix the fresh frozen tissue sections with 4% Formaldehyde for 20 mins, wash three times with PBS. (2) Add 1% bovine serum albumin (BSA) in PBS to block the tissue and incubate for 30 mins at RT. (3) Wash the tissue with PBS for three times. (4) Add the mixture of three antibodies (final concentration 25 µg/mL in 1% BSA, PBS) to the tissue, need around 50 µL. Incubate for 1 hour in dark at RT. (5) Wash the tissue with PBS for three times, with 5 mins washing each time. (6) Dip the tissue in water shortly and air dry the tissue. (7) Image the tissue using EVOS (Thermo Fisher EVOS fl), at a magnification of 10 x. Filters used are Cy5, RFP and GFP.

#### Application of DNA-antibody conjugates to the tissue slide

In order to obtain spatial proteomic information, we incubated the fixed tissue slide with a cocktail of DNA-antibody conjugates prior to microfluidic spatial barcoding. The cocktail was prepared by combining 0.1 µg of each DNA-antibody conjugates (see Table S1). The tissue slide was first blocked with 1% BSA/PBS plus RNase inhibitor, and then incubated with the cocktail for 30 minutes at 4°C. Afterwards, the tissue slide was washed 3 times with a washing buffer containing 1% BSA + 0.01% Tween 20 in 1X PBS and one time with DI water prior to attaching the first PDMS microfluidic chip.

#### Adding the first set of barcodes and reverse transcription

To perform spatial barcoding of mRNAs for transcriptomic mapping, the slides were blocked by 1% BSA plus RNase inhibitor (0.05U/μL, Enzymatics) for 30 minutes at room temperature. After cleaning with 1x PBS and quickly with DI water, the first PDMS microfluidic chip was roughly aligned and placed on the tissue glass slide such that the center of the flow barcoding region covered the tissue of interest. This tissue section was then permeabilized by loading 0.5% Triton X-100 in PBS into each of the 50 channels followed by incubation for 20 minutes and finally were cleaned thoroughly by flowing through 20μL of 1X PBS. A vial of RT mix was made from 50 μL of RT buffer (5X, Maxima H Minus kit), 32.8 µL of RNase free water, 1.6 µL of RNase Inhibitor (Enzymatics), 3.1 µL of SuperaseIn RNase Inhibitor (Ambion), 12.5 µL of dNTPs (10 mM, Thermo Fisher), 25 µL of Reverse Transcriptase (Thermo Fisher), 100 µL of 0.5X PBS with Inhibitor (0.05U/μL, Enzymatics). To perform the 1^st^ microfluidic flow barcoding, we added to each inset a 5 µL of solution containing 4.5 µL of the RT mix described and 0.5 µL of one of the 50 DNA barcodes (A1-A50) solution (25 µM), and then pulled in using a house vacuum for <3 minutes depending on channel width. Afterwards, the binding of DNA oligomers to mRNAs fixed in tissue was allowed to occur at room temperature for 30 minutes and then incubated at 42 °C for 1.5 hours for *in situ* reverse transcription. To prevent the evaporation of solution inside the channels, the whole device was kept inside a sealed wet chamber(Gervais and Delamarche, 2009). Finally, the channels were rinsed by flowing NEB buffer 3.1(1X, New England Biolabs) supplemented with 1% RNase inhibitor (Enzymatics) continuously for 10 minutes. During the flow barcoding step, optical images could be taken to record the exact positions of these microfluidic channels in relation to the tissue section subjected to spatial barcoding. It was done using an EVOS microscope (Thermo Fisher EVOS fl) in a light or dark field mode. Then the clamp was removed and the PDMS chip was detached from the tissue slide, which was subsequently dipped into a 50 mL Eppendorf tube containing RNase free water to rinse off remaining salts.

#### Adding the second set of barcodes and ligation

After drying the tissue slides, the second PDMS chip with the microfluidic channels perpendicular to the direction of the first PDMS chip in the tissue barcoding region was carefully aligned and attached to the tissue slide such that the microfluidic channels cover the tissue region of interest. The ligation mix was prepared as follows: 69.5 µL of RNase free water, 27 µL of T4 DNA ligase buffer (10X, New England Biolabs), 11 µL T4 DNA ligase (400 U/µL, New England Biolabs), 2.2 µL RNase inhibitor (40 U/µL, Enzymatics), 0.7 µL SuperaseIn RNase Inhibitor (20 U/µL, Ambion), 5.4 µL of Triton X-100 (5%). To perform the second flow barcoding, we added to each channel a total of 5 µL of solution consisting of 2 µL of the aforementioned ligation mix, 2 µL of NEB buffer 3.1(1X, New England Biolabs) and 1 µL of DNA barcode B (25 µM). Reaction was allowed to occur at 37 °C for 30 minutes and then the microfluidic channels were washed by flowing 1X PBS supplemented with 0.1% Triton X-100 and 0.25% SUPERase In RNase Inhibitor for 10 minutes. Again, the images showing the location of the microfluidic channels on the tissue slide could be taken during the flow step under the light or dark field optical microscope (Thermo Fisher EVOS fl) before peeling off the second PDMS chip.

#### cDNA collection and purification

We devised a square well PDMS gasket, which could be aligned and placed on the tissue slide, creating an open reservoir to load lysis buffer specifically to the flow barcoded tissue region to collect cDNAs of interest. Depending on the area of this region, the typical amount of buffer is 10 - 100 µL of Proteinase K lysis solution, which contains 2 mg/mL proteinase K (Thermo Fisher), 10 mM Tris (pH = 8.0), 200 mM NaCl, 50 mM EDTA and 2% SDS. Lysis was carried out at 55 °C for 2 hours. The lysate was then collected and stored at -80 °C prior to use. The cDNAs in the lysate were purified using streptavidin beads (Dynabeads MyOne Streptavidin C1 beads, Thermo Fisher). The beads (40 µL) were first washed three times with 1X B&W buffer (Ref to manufacturer’s manual) with 0.05% Tween-20, and then stored in 100 µL of 2X B&W buffer (with 2 μL of SUPERase In Rnase Inhibitor). To perform purification from stored tissue lysate, it was allowed to thaw, and the volume was brought up to 100 µL by RNase free water. Then, 5 µL of PMSF (100 µM, Sigma) was added to the lysate and incubated for 10 minutes at room temperature to inhibit the activity of Proteinase K. Next, 100 µL of the cleaned streptavidin bead suspension was added to the lysate and incubated for 60 minutes with gentle rotating. The beads with cDNA were further cleaned with 1X B&W buffer for two times and then with 1X Tris buffer (with 0.1% Tween-20) once.

#### Template switch and PCR amplification

The cDNAs bound to beads were cleaned and resuspended into the template switch solution. The template switch reaction mix contains 44 μL of 5X Maxima RT buffer (Thermo Fisher), 44 μL of 20% Ficoll PM-400 solution (Sigma), 22 μL of 10 mM dNTPs each (Thermo Fisher), 5.5 μL of RNase Inhibitor (Enzymatics), 11 μL of Maxima H Minus Reverse Transcriptase (Thermo Fisher), and 5.5 μL of a template switch primer (100 μM). The reaction was conducted at room temperature for 30 minutes followed by an additional incubation at 42 °C for 90 minutes. The beads were rinsed once with a buffer containing 10 mM Tris and 0.1% Tween-20 and then rinsed again with RNase free water using a magnetic separation process. PCR was conducted following these two steps. In the first step, a mixture of 110 µL Kapa HiFi HotStart Master Mix (Kapa Biosystems), 8.8 μL of 10 μM stocks of primers 1 and 2, and 92.4 μL of water was added to the cleaned beads. If the protein detection was conducted in conjunction using a process similar to CITE-seq, a primer 3 solution (1.1 μL, 10 μM) was also added at this step. PCR reaction was then done using the following conditions: first incubate at 95°C for 3 mins, then cycle five times at 98°C for 20 seconds, 65°C for 45 seconds, 72°C for 3 minutes and then the beads were removed from the solution by magnet. Evagreen (20X, Biotium) was added to the supernatant with 1:20 ratio, and a vial of the resultant solution was loaded into a qPCR machine (BioRad) to perform a second PCR step with an initial incubation at 95°C for 3 minutes, then cycled at 98°C for 20 seconds, 65°C for 20 seconds, and finally 72°C for 3 minutes. The reaction was stopped when the fluorescence signal just reached the plateau.

#### Amplicon purification, sequencing library preparation and quality assessment

The PCR product was then purified by Ampure XP beads (Beckman Coulter) at 0.6×ratio. The mRNA-derived cDNAs (>300 bp) were then collected from the beads. If the cDNAs were less than 300 bp, they remained in the supernatant fraction. If the protein detection was conducted like CITE-seq, this fraction was used instead. For sequencing antibody-DNA conjugate-derived cDNAs, we further purified the supernatant using 2X Ampure XP beads. The purified cDNA was then amplified using a PCR reaction mix containing 45 µL purified cDNA fraction, 50 µL 2x KAPA Hifi PCR Master Mix(Kapa Biosystems), 2.5 µl P7 primer of 10 µM and 2.5 µL P5 cite primer at 10 µM. PCR was performed in the following conditions: first incubated at 95°C for 3 minutes, then cycled at 95°C for 20 seconds, 60°C for 30 seconds and 72 °C for 20 seconds, for 10 cycles, lastly 72 °C for 5 minutes. The PCR product was further purified by 1.6X Ampure XP beads. For sequencing mRNA-derived cDNAs, the quality of amplicon was analyzed firstly using Qubit (Life Technologies) and then using an Agilent Bioanalyzer High Sensitivity Chip. The sequencing library was then built with a Nextera XT kit (Illumina) and sequenced using a HiSeq 4000 sequencer using a pair-end 100x100 mode. To conduct joint profiling of proteins and mRNAs, the DNA-antibody conjugate-derived sequencing library was combined with mRNA-derived cDNA library at a 1:9 ratio, which is sufficient to detect the finite set of proteins and minimally affects the sequencing depth required for mRNAs.

#### Tissue fluorescent staining before DBiT-seq

Fluorescent staining of tissue sections with either common nucleus staining dyes or fluorescent labelled antibodies can be performed before the DBiT-seq to facilitate the identification of tissue region of interest. After the DBiT-seq fixation procedure with formaldehyde, the whole tissue was permeabilized with 0.5% Triton X-100 in PBS for 20 minutes and cleaned with 1X PBS for three times. Working solution mixture of DAPI and phalloidin (FITC labelled) were added on top of the tissue and then incubate at room temperature for 20 minutes. After washing thrice with 1X PBS, tissue sections were blocked with 1% BSA for 30 minutes. Finally, antibody with fluorescent labels (here we use P2RY12) were added and incubated at room temperature for 1 hour. Images of the tissue were taken using EVOS microscope (Thermo Fisher EVOS fl), with a resolution of 10 x. Filters used were DAPI, GFP and RFP. DBiT-seq barcoding procedure could be continued after staining.

#### smFISH and comparison with DBiT-seq

Single molecular fish (smFISH) was performed using HCR v3.0 kit (Molecular Instruments, Inc) following manufacture protocols. Probes used in current study included Ttn, sfrp2, Trf and Dlk1. smFISH z-stack images were taken using a ZEISS LSM 880 confocal microscope with a 60x oil immersion objective. The smFISH quantitation was performed using FISH-quant(https://biii.eu/fish-quant). mRNA transcript count was an average of three fields of view with each having a size of 306 x 306 µm. The sum of DBiT-seq transcript counts in the same locations were also calculated and compared side by side with smFISH counts.

#### Cell number counting in each pixel

Cell numbers for each pixel were counted manually using DAPI and ethidium homodimer-1 stained tissue images (Figure S1C). The total cell counts were obtained by summing the nucleus numbers in each of the pixels. If a nucleus appeared at the edge of a pixel, we would count it as 1 if more than half of the nucleus lied within the pixel and as 0 if otherwise. A total of 50 pixels were counted and the averaged numbers were reported.

### QUANTIFICATION AND STATISTICAL ANALYSIS

#### Sequence alignment and generation of gene expression matrix

To obtain transcriptomics data, the Read 2 was processed by extracting the UMI, Barcode A and Barcode B. The processed read 1 was trimmed, mapped against the mouse genome(GRCh38), demultiplexed and annotated (Gencode release M11) using the ST pipeline v1.7.2 (Navarro et al., 2017), which generated the digital gene expression matrix for down-stream analysis. The rows of the gene matrix correspond to pixels, defined by their location info (barcode A x barcode B) and columns correspond to genes. Normalization of gene data was completed through either Scran (V3.11) or Seurat’s (V3.2) “SCTransform normalization”.

For proteomics data, the Read 2 was processed by extracting the antibody-derived barcode, spatial Barcode A and Barcode B. The processed read was trimmed, demultiplexed using the ST pipeline v1.7.2 (Navarro et al., 2017), which generated the gene protein matrix for down-stream analysis. Similar to the gene expression matrix, the rows correspond to pixels, defined by (barcode A x barcode B) and columns correspond to proteins.

The pan-mRNA and pan-protein heatmap plots in Figure 2A were generated using raw UMI counts without normalization.

#### Clustering whole mouse embryo data

Spatially variable genes were identified by SpatialDE(Svensson et al., 2018b). The resulting list list of differentially expressed genes was submitted to ToppGene(Chen et al., 2009) for GO and Pathway enrichment analysis. Spatially variable genes generated by SpatialDE were used to conduct the clustering analysis. Non-negative matrix factorization(NMF) was performed using the NNLM pacakges in R, after the raw expression values were log-transformed. We chose k of 11 for the mouse embryo DBiT-seq transcriptome data obtained at a 50μm pixel size. For each pixel, the largest factor loading from NMF was used to assign cluster membership. NMF clustering of pixels was plotted by tSNE using the package “Rtsne” in R.

#### Comparison with ENCODE bulk sequencing data

Public bulk RNA-Seq datasets were downloaded from ENCODE (liver, heart and neural tube from mouse embryo E11.5) and the raw expression counts were normalized with FPKM. For DBiT-seq data, “pseudo-bulk” gene expression profiles were obtained by summing counts for each gene in each tissue region and divided by the sum of total UMI counts in this specific region, and further multiplied by 1 million. The scatter plots were plotted using log10(FPKM+1) value for bulk data and log10(pseudo gene expression+1)) for DBiT-seq data. Pairwise Pearson correlation coefficients were calculated. Good correlations (r >0.784) were observed between the two different sets of data.

#### Gene length bias analysis

Gene length bias is well understood in bulk RNA-seq data. We further analyzed our DBiT-seq data and ST data using reference package GeneLengthBias for RNAseq data (Phipson et al., 2019) following standard protocols.

#### Data analysis with single-cell RNA-seq analysis workflow

The data analysis of E10-E12 tissue sections was carried out with Seurat V3.2 (Butler et al., 2018; Stuart et al., 2019) following standard procedures. In short, data normalization, transformation, and selection of variable genes were performed using the SCTransform function with default settings. Principal component analysis (PCA) was performed on the top 3,000 variable genes using the RunPCA function, and the first 30 principal components were used for Shared Nearest Neighbor (SNN) graph construction using the FindNeighbors function. Clusters were then identified using the FindClusters function. We used Uniform Manifold Approximation and Projection (UMAP) to visualize DBiT-seq data in a reduced two-dimensional space (McInnes et al., 2018). To identify differentially expressed genes for every cluster, pair-wise comparisons of cells in individual clusters against all remaining cells were performed using the FindAllMarkers function (settings: min.pct = 0.25, logfc.threshold = 0.25). Expression heatmap was then generated using top 10 differentially expressed genes in each cluster.

#### Cell type identification

Automatic cell type identification for E11 mouse tail region (Figure 6) was achieved with SingleR (version 1.2.3) (Aran et al., 2019) following standard procedure. Single cell RNA-seq data E10.5 from (Cao et al., 2019) was used as the reference. The 12 most frequent cell types were shown in the UMAP, and cell types with small size were shown as “other”.

Cell type identification for E10 Eye region (Figure 4) was performed through integration with scRNA-seq reference data. We combined DBiT-seq data with scRNA-seq data of mouse embryo E9.5 and E10.5 (Cao et al., 2019) using Seurat V3.2 and did the clustering after “SCTransform” procedure. DBiT-seq data showed a similar distribution as scRNA-seq reference data. We then assign each cluster with a cell type using cell type information from the reference data (if two cell types presented in one cluster, the major cell types were assigned). The cell type of each pixel was then assigned by their cluster number.

### DATA AND CODE AVAILABILITY

#### Data availability

The accession number for the sequencing data reported in this paper is submitted to GEO: https://www.ncbi.nlm.nih.gov/geo/query/acc.cgi?acc=GSE137986 The reviewer access code is: ubmbcwkgvhunzqp

#### Code availability

Code for sequencing data analysis is available (https://github.com/MingyuYang-Yale/DBiT-seq).

## SUPPLEMENTAL INFORMATION

### Supplementary Figures

**Figure S1.**
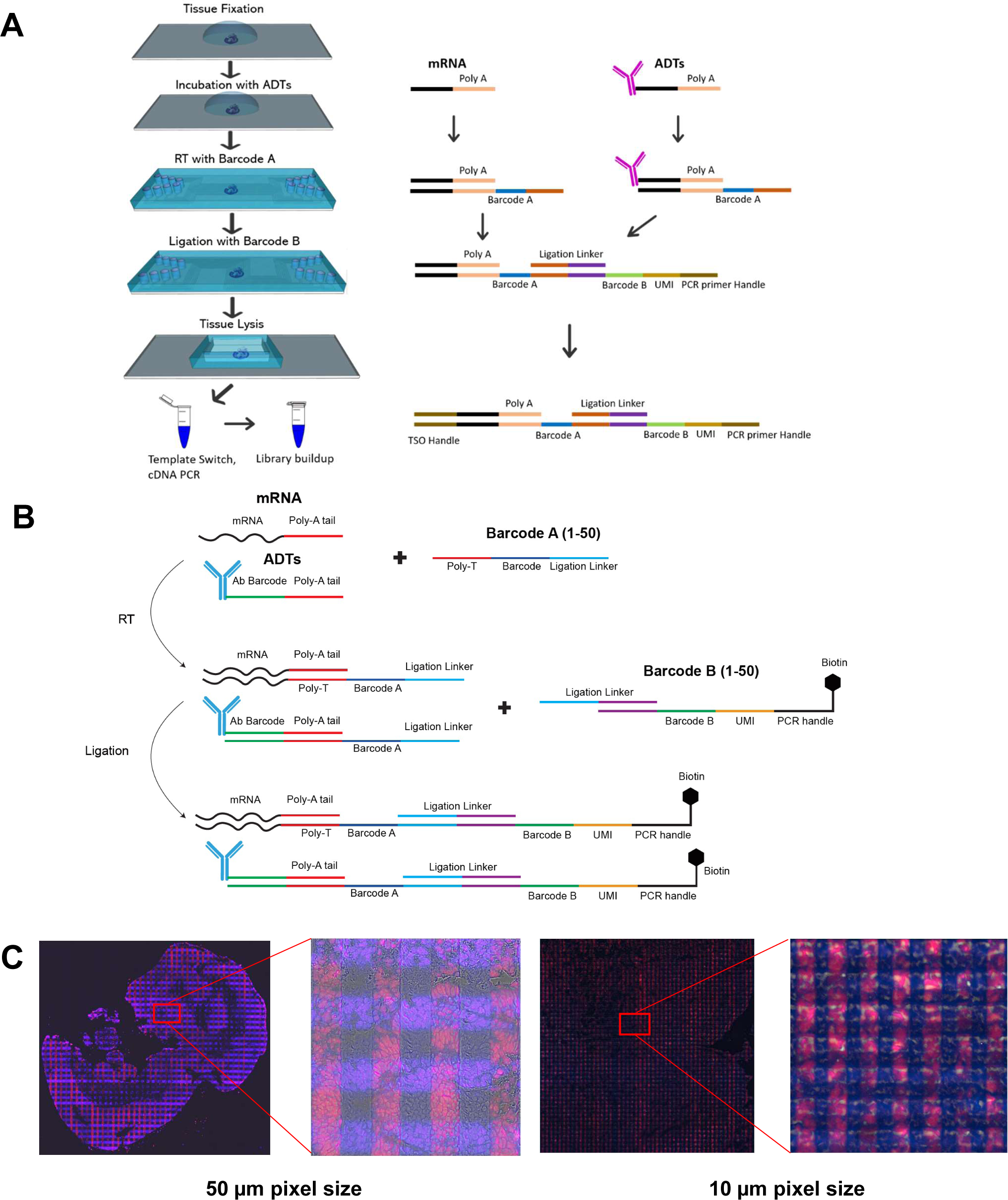
Experimental design of DBiT-seq and fluorescent staining-based verification. (A) Detailed microfluidic device design (left panel) and barcoding chemistry protocol (right panel). Left panel: fresh frozen tissue sections were first allowed to warm to room temperature for 10 minutes. Then, 4% Formaldehyde was added, and tissue was fixed for 20 minutes at room temperature. After fixation, a cocktail of 22 antibody-DNA tags (ADTs) were added and incubated at 4°C for 30 minutes. After washing three times with PBS, 1^st^ PDMS chip was attached to the glass slide. Barcode A (A1-A50) along with reverse transcription mixture was flowed through each channel. After reverse transcription, the 1^st^ PDMS chip was removed and a 2^nd^ PDMS was attached. Ligation solution along with Barcode B (B1-B50) was flowed into each channel. When finished, the 2^nd^ PDMS chip was removed and a PDMS gasket was attached to the glass slide. Lysis solution was added into the gasket and the lysate was collected. cDNA and ADT derived cDNA were extracted using streptavidin coated magnetic beads. Template switch and PCR were then performed. The sequencing library was finally built with standard tagmentation. Right panel: DNA barcode A consists of a poly T region, a barcode region and a ligation region. The poly T region will recognize the poly A tail of mRNA and ADTs. DNA Barcode B consists of a ligation region, a barcode region, a UMI region and a PCR primer handle region. During ligation process, the ligation region will be ligated to the ligation region of barcode A. The cDNA product will then be template-switched. The final product is further amplified by PCR. (B) Schematic of the biochemistry protocol to add spatial barcodes to a tissue slide. Proteins of interest are labeled with antibody DNA tags (ADTs), each of which consists of a unique antibody barcode (15mer, see Table S1) and a poly-A tail. Barcode A1-A50 contains a ligation linker(15mer), a unique spatial barcode Ai (i=1:50, 8mer, see Table S2), and a poly-T sequence(16mer), which detects mRNAs and proteins through binding to poly-A tails. After introducing barcodes A1-A50 to the tissue slide, reverse transcription is conducted *in situ* to generate cDNAs from mRNAs as well as antibody barcodes. Barcode B1-B50 consists of a ligation linker(15mer), a unique spatial barcode Bj(j=1:50, 8mer, see Table S2), a unique molecular identifier (UMI)(10mer), and a PCR handle (22mer) terminally functionalized with biotin, which facilitates the purification in the later steps using streptavidin-coated magnetic beads. When the barcodes B1-B50 are introduced to the tissue sample that is already barcoded with A1-A50 using an orthogonal microfluidic delivery, a complementary ligation linker is also introduced and initiates the covalent ligation of barcodes A and B, giving rise to a 2D array of spatially distinct barcodes AiBj (i=1:50 and j=1:50). (C) DAPI and ethidium homodimer-1 (both are nuclei staining dyes) staining of mouse embryo (E10) tissue section. After attaching the 1^st^ PDMS chip, DAPI (purple color) solution was flowed through each channel and incubated for 20 minutes at room temperature. 1^st^ PDMS was removed and a 2^nd^ PDMS was attached. Ethidium homodimer-1 (Red) was then flowed through. The resulted tissue will form a mosaic-like structure, with the intersections to be stained by both nuclei dyes.

**Figure S2.**
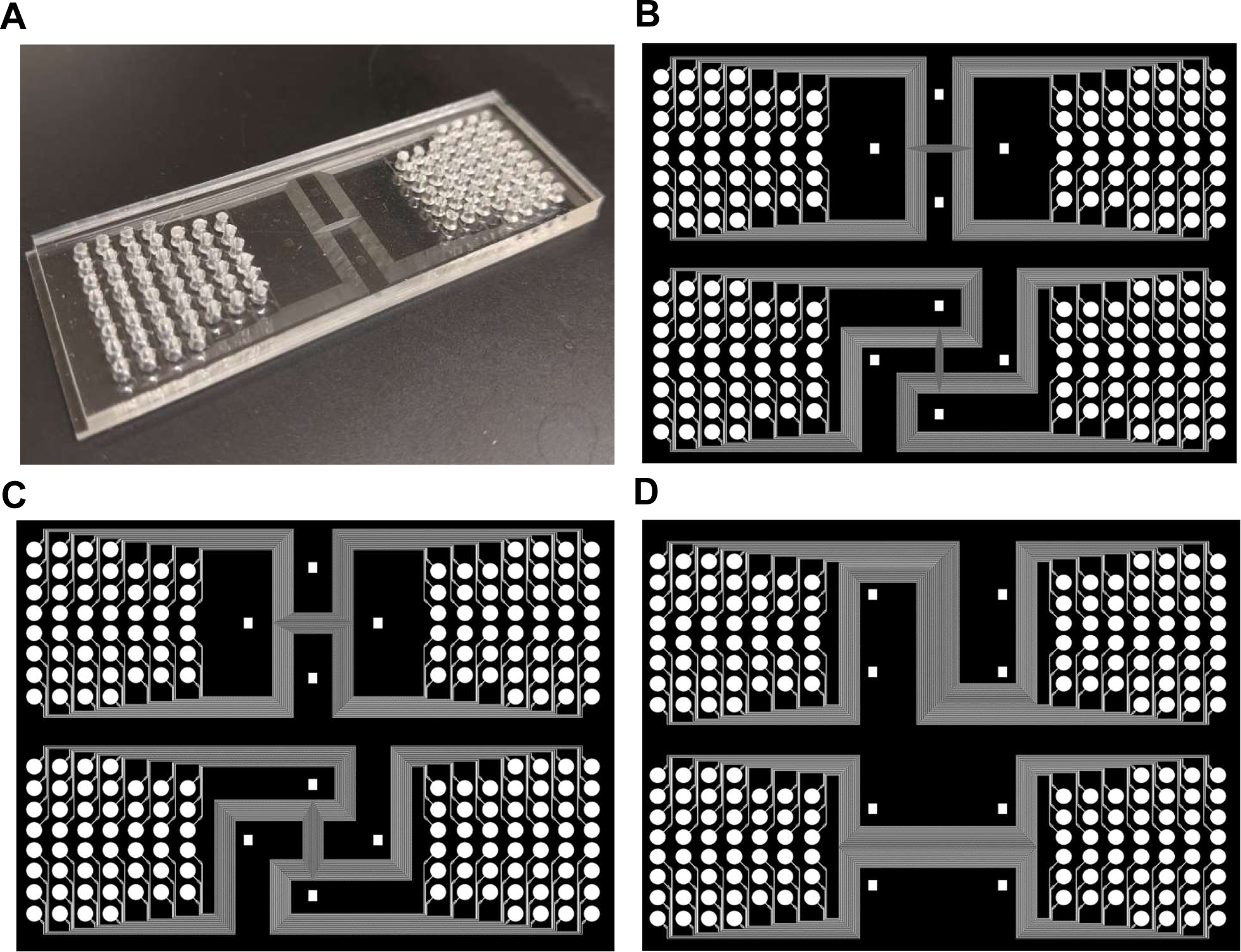
Device design with different channel widths. (A) A photograph of PDMS chip (1^st^ direction) with fifty 10µm-width channels in the center. Holes were punched with a 2mm diameter puncher. (B) AutoCAD design of PDMS chip with 10µm width channels. (C) AutoCAD design of PDMS chip with 25 µm width channels. (D) AutoCAD design of PDMS chip with 50 µm width channels.

**Figure S3.**
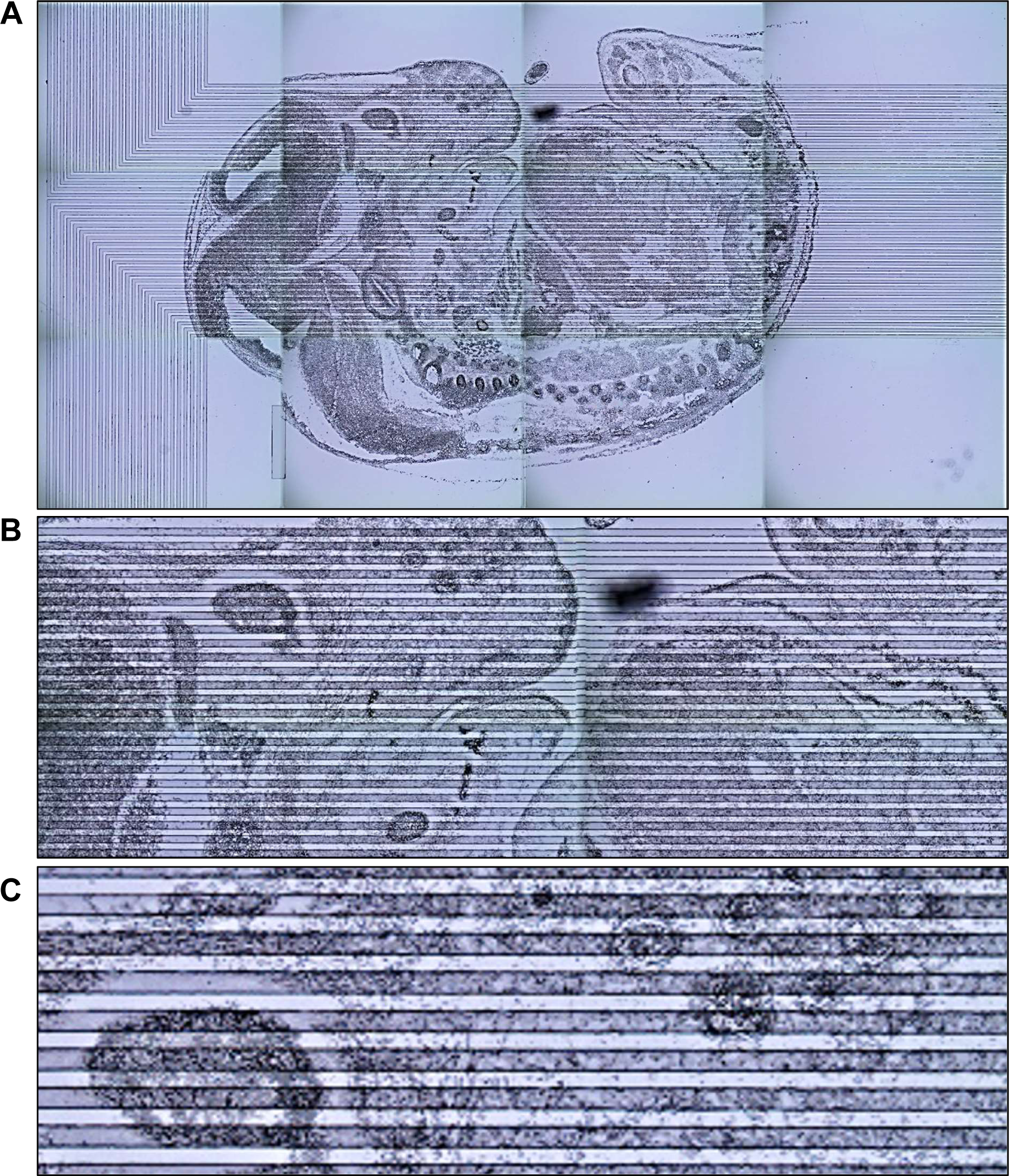
Optical images showing the microfluidic channels placed on an embryonic tissue surface. These pictures are from a PDMS chip (50 µm channel width) covering mouse embryo (E14) tissue slides. (A) Whole embryonic tissue; (B) Enlarged view of front head and a section of the lower body. (C) Enlarged view of a region at the front head.

**Figure S4.**
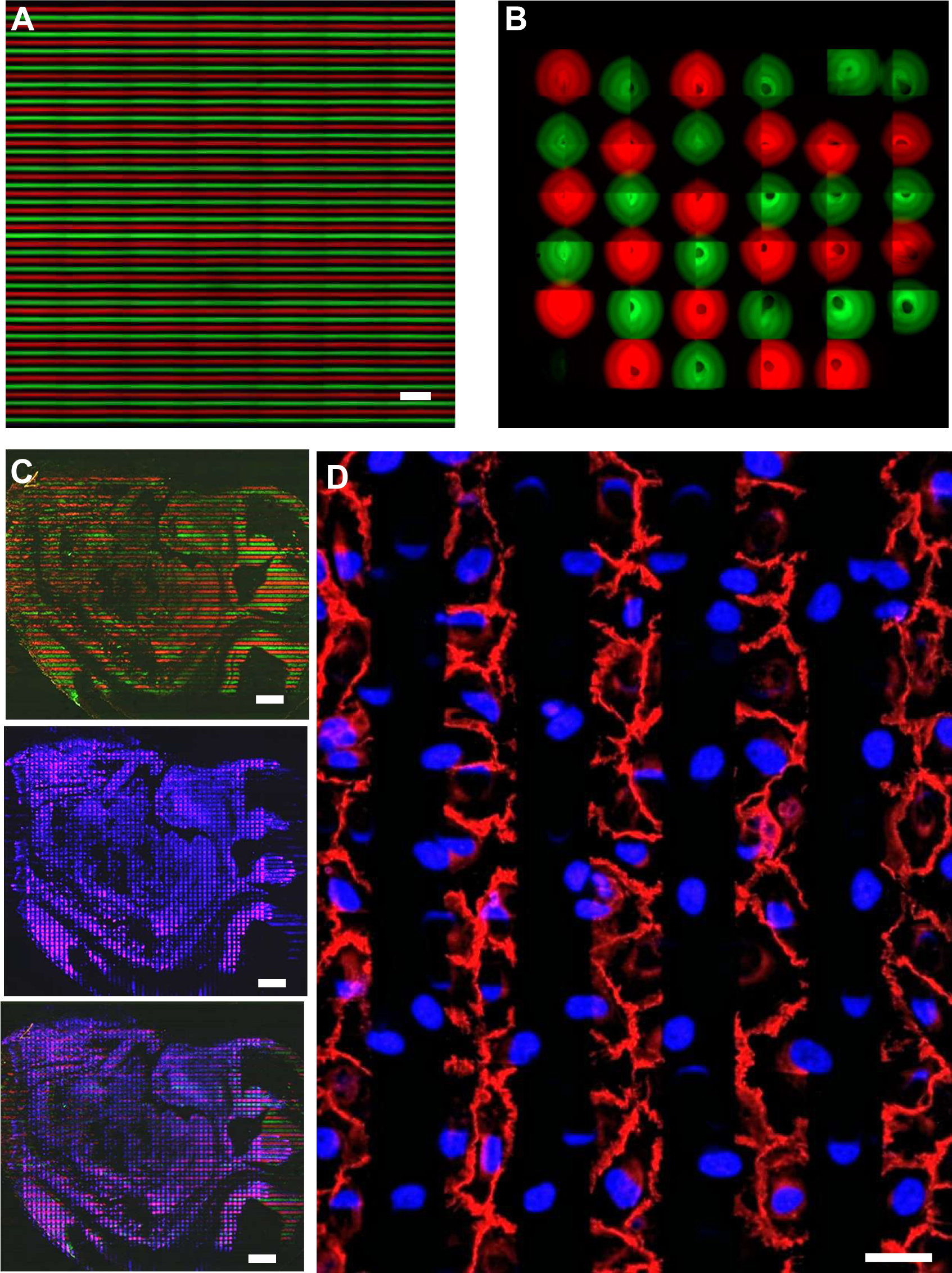
Flow barcoding evaluation with a 50 -tm channel width microfluidic system. (A) Testing possible crosstalk between two neighboring channels by alternately flowing an FITC (Green) labelled Barcode A and a Cy3 (Red) labelled Barcode A. Scale bar= 400 -tm. (B) Analyzing possible leakage at the injection area. Each circle represents an inset hole. Green colored circles are loaded with FITC labelled Barcode A and red colored circles are loaded with Cy3 labelled Barcode A. (C) Staining mouse embryo (E14) mRNAs with FITC (Green) labelled Barcode A and Cy3 (Red) labelled Barcode A (upper panel). Further staining with Cy5 labelled Barcode B (Middle panel). Overlay of two previous panels (lower panel). Scale bar = 600 µm. (D) Staining a layer of fixed NIH 3T3 cells using 50µm microchannels. The first flow (horizonal) introduced DAPI to stain nuclei (blue) and the second flow (vertical) introduced phalloidin to stain for actin fiber cytoskeleton (red). Scale bar = 50 µm.

**Figure S5.**
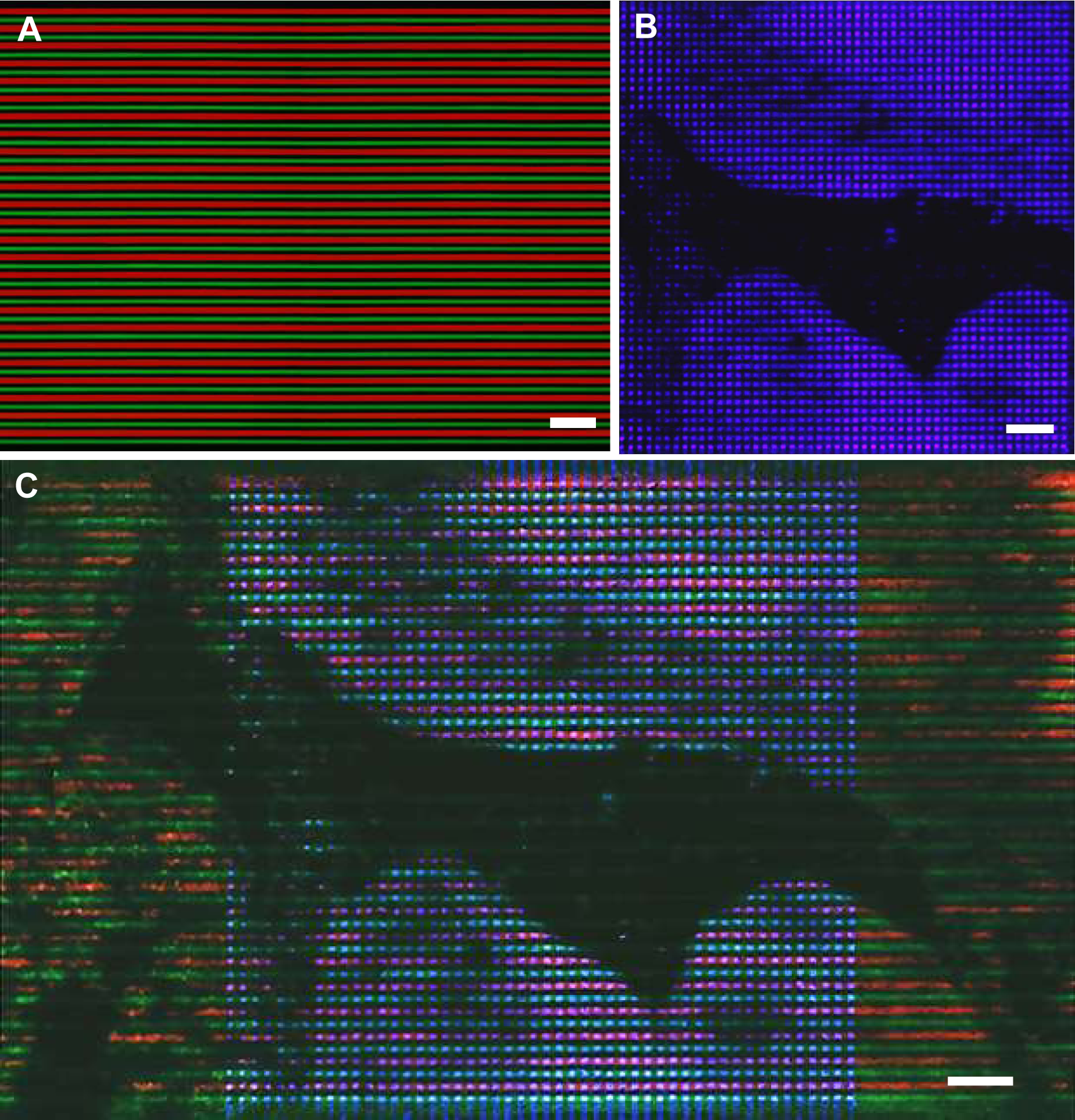
Flow barcoding evaluation with a 10 µm channel width microfluidic system. (A) Measuring possible crosstalk between two neighboring channels by alternately flowing an FITC (Green) labelled Barcode A and a Cy3 (Red) labelled Barcode A. Scale bar = 100µm. (B) Staining part of mouse embryo (E14) mRNAs with FITC (Green) labelled Barcode A and Cy3 (Red) labelled Barcode A (Fluorescence not shown). Further staining with Cy5 (purple) labelled Barcode B. Scale bar = 100µm. (C) Overlay of three fluorescent channels of Figure S5(B). Scale bar = 100µm.

**Figure S6.**
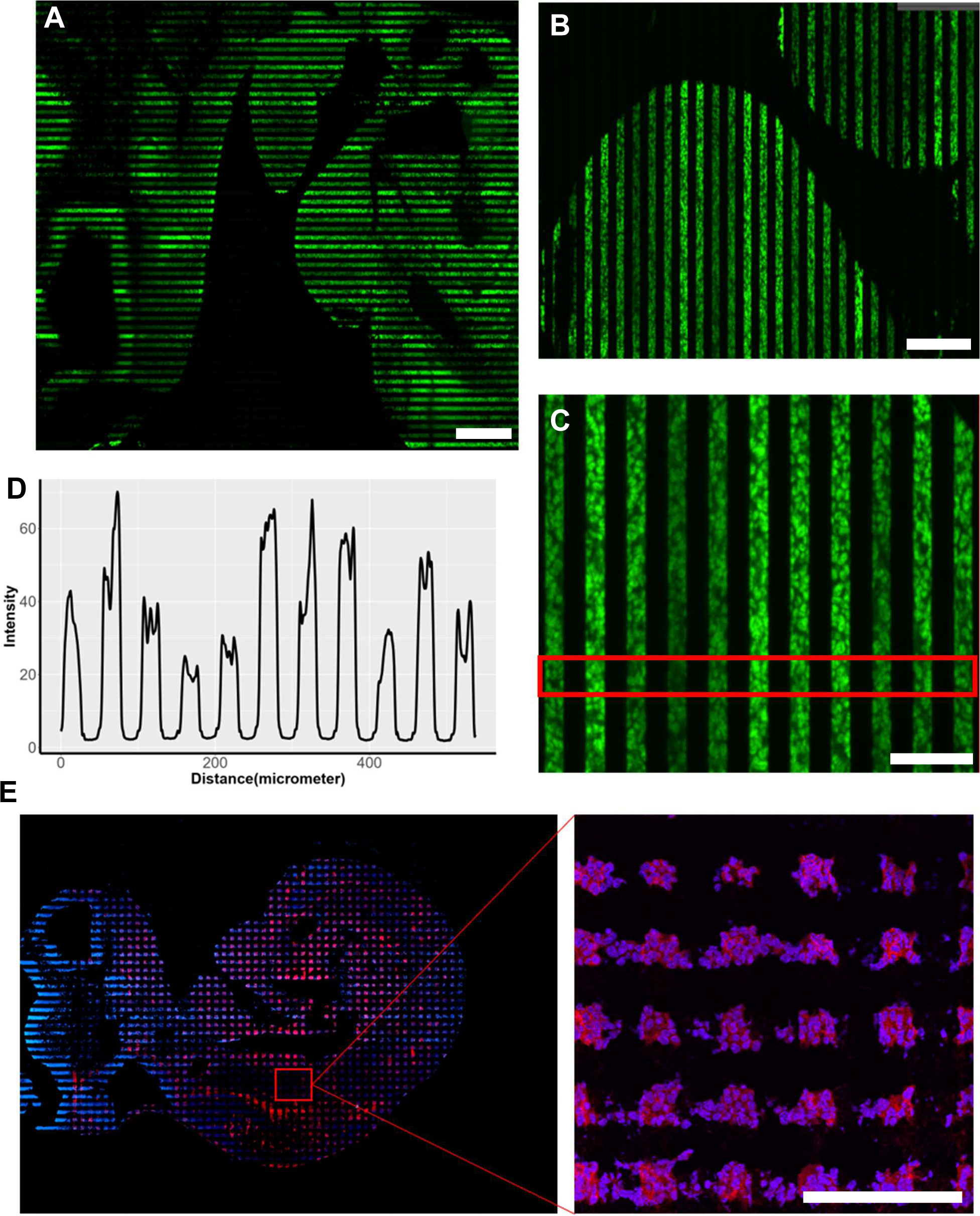
Flow barcoding evaluation with a 25 -tm channel width microfluidic system. (A) Measuring possible crosstalk between two neighboring channels by alternately flowing an I-lTC (Green) labelled Barcode A. Scale bar= 300 -tm. (B) Zoomed-in picture of picture A. Scale bar = 200 µm. (C) Zoomed-in picture of picture B. Scale bar = 100 µm. (D) Quantification of fluorescence signals along the red box in C. (E) left panel: Staining mouse embryo (E10) mRNAs with DAPI (Blue) and ethidium homodimer-1 (Red); Right panel: enlarged picture of E. Scale bar = 100 µm.

**Figure S7.**
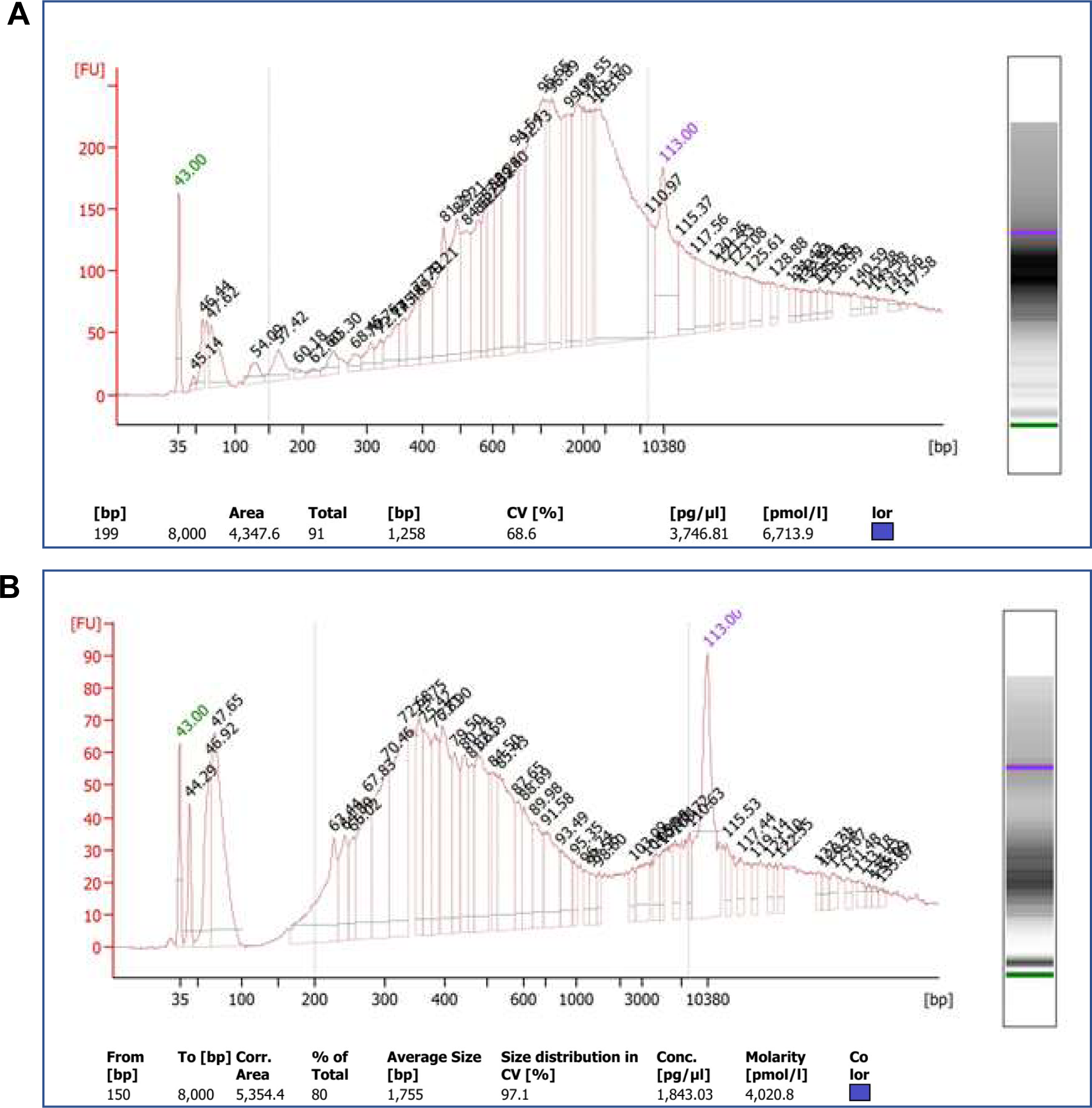
Size distribution of cDNA amplicons from mouse embryo samples. (A) Fresh frozen tissue sample tested with DBiT-seq immediately. (B) Fresh frozen sample left at room temperature for 24 hours before testing with DBiT-seq. Due to degradation, shorter cDNA was observed when left at room temperature for an extended time.

**Figure S8.**
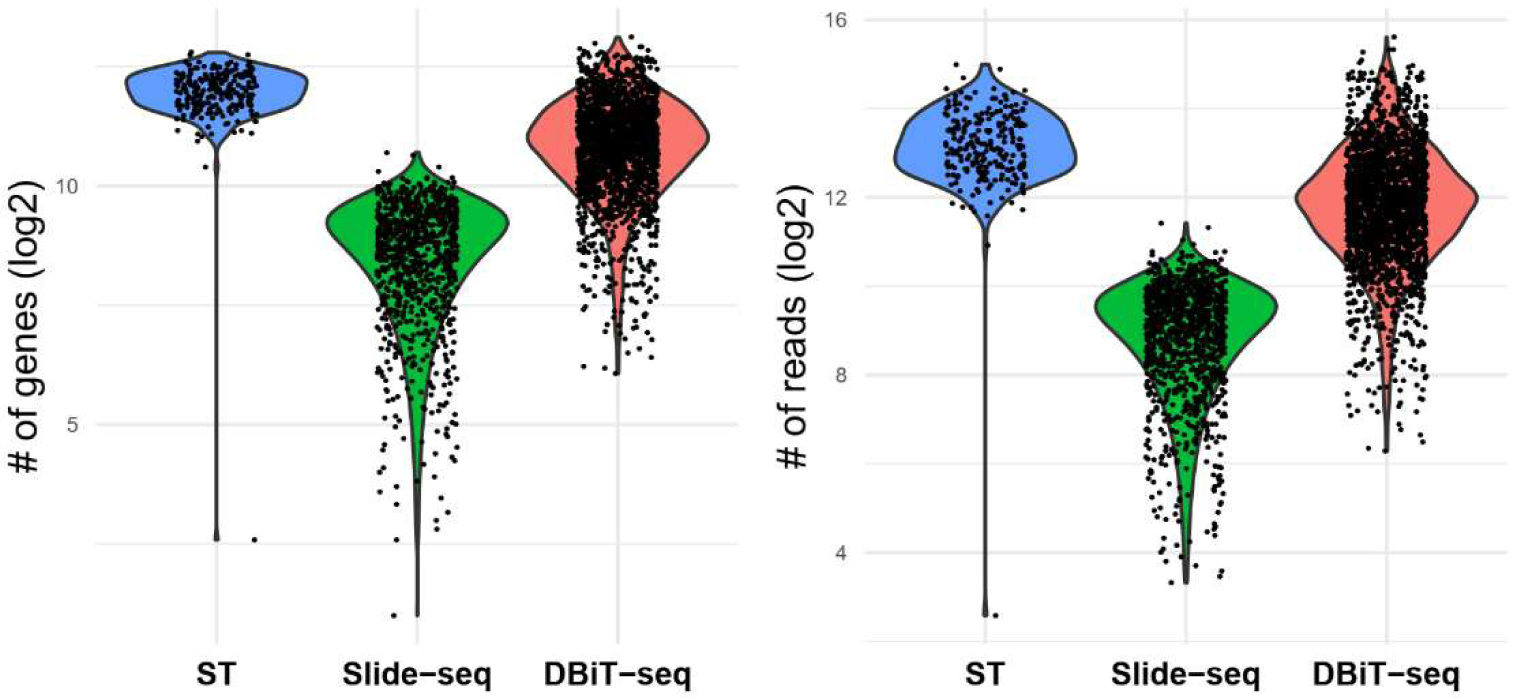
Comparison of the # of genes and UMI reads per pixel with other methods. Our sequencing data was obtained using a 10μm microfluidic barcoding approach. The number of reads per pixel was close to 5,000. Slide-seq only had ∼200 reads per pixel (10μm and the low-resolution ST method yielded a similar number of reads per pixel, but the pixel size was ∼100-150μm, about 100 times larger than ours.

**Figure S9.**
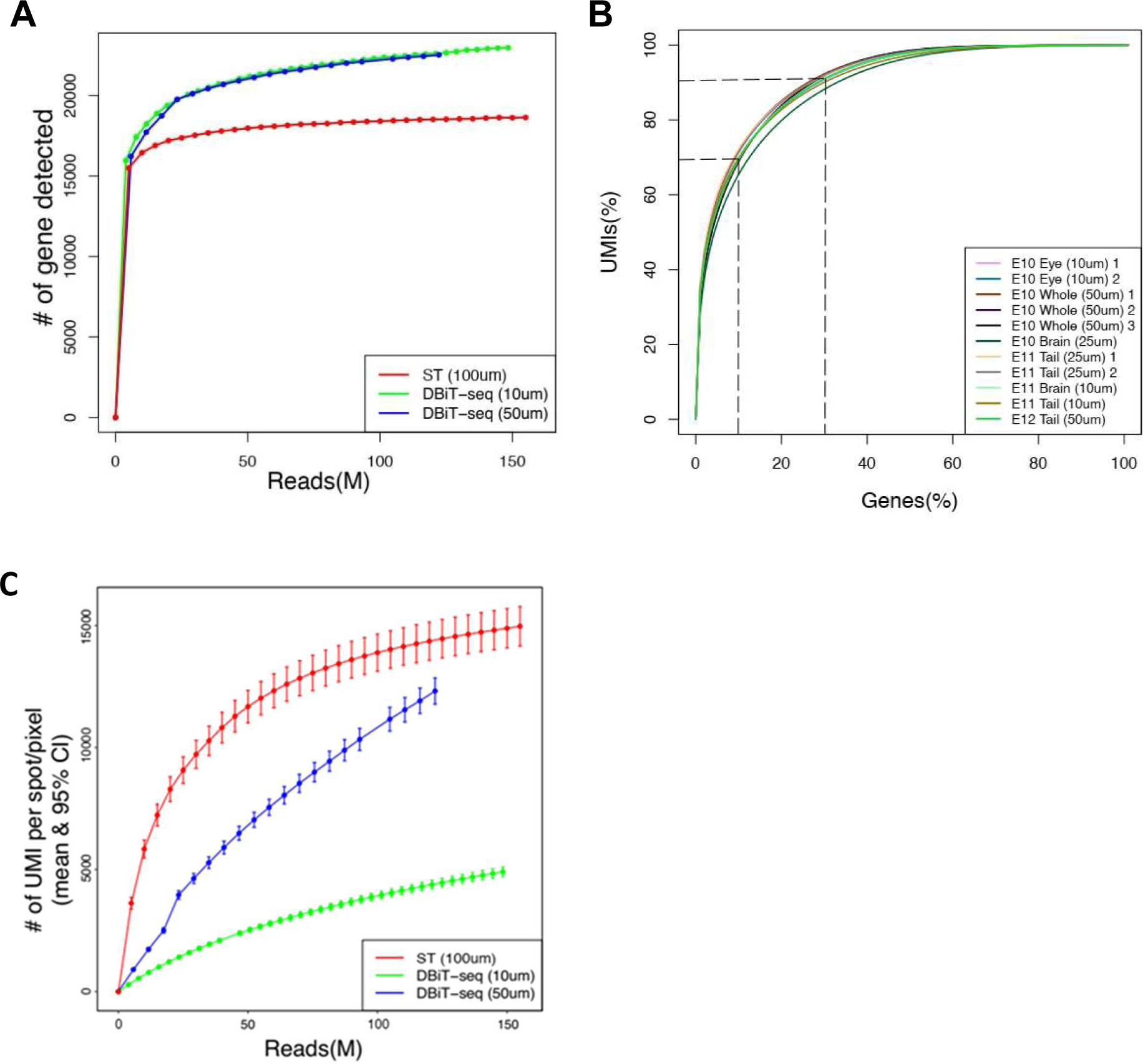
Saturation curve of DBiT-seq and cumulative number of genes vs. UMIs. (A) The saturation curves of DBiT-seq with different resolutions (10µm and 50µm) are nearly identical. DBiT-seq had more genes detected, but this could be due to sample differences. Overall, the DBiT-seq curves are quite similar to ST method. (B) Cumulative number of genes vs. UMI. It is observed that top 10% genes account for 70% of the UMIs and top 30% genes account for 90% of UMIs. (C) Number of UMIs per pixel vs. raw reads.

**Figure S10.**
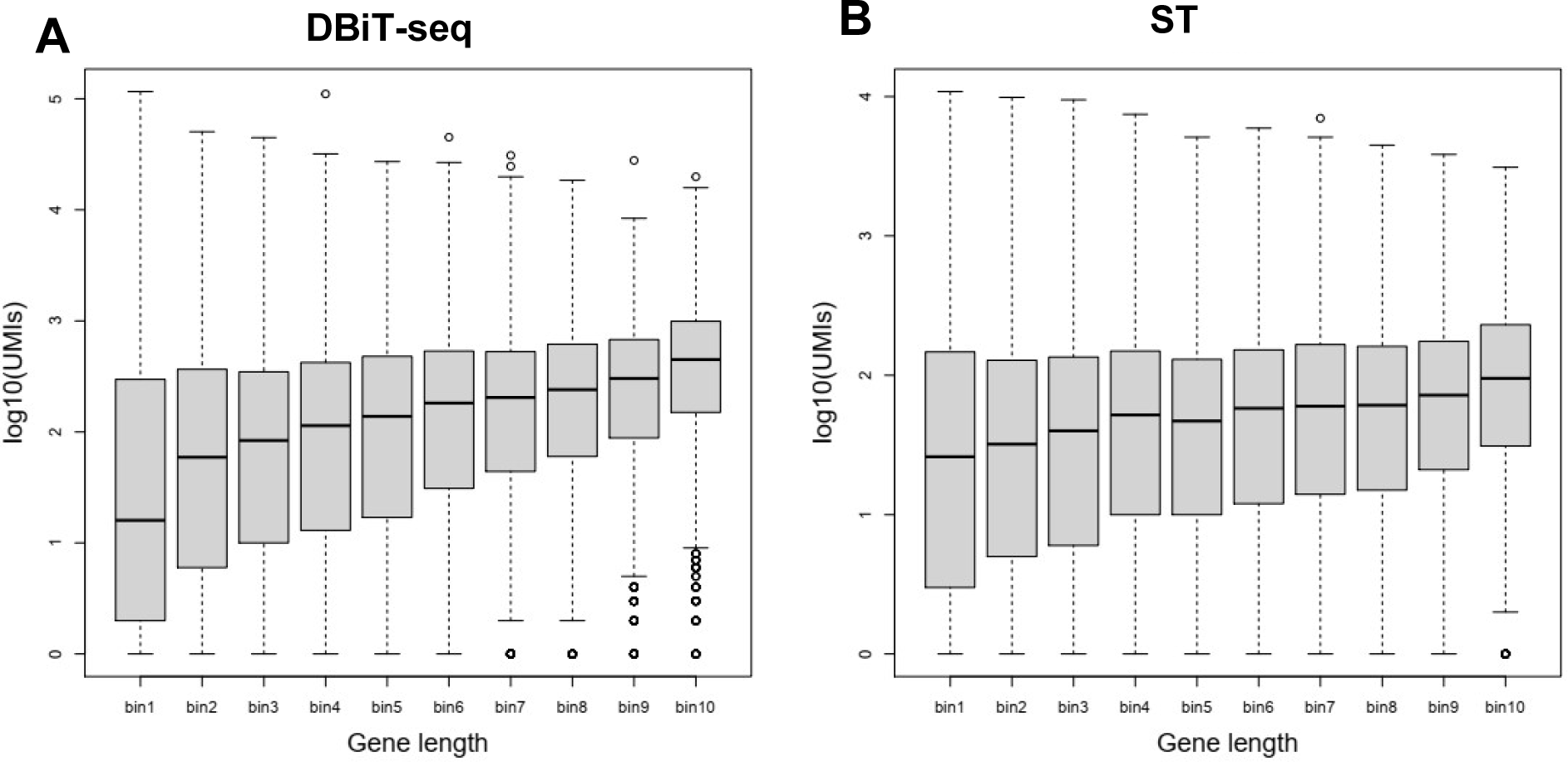
Gene length bias analysis of DBiT-seq and ST data. (A) Gene length bias analysis of DBiT-seq sample E10 with 50 um resolution. (B) Gene length bias analysis of ST sample “Rep9”. It is observed that gene length bias existed in both methods. The bias shows that shorter genes have lower UMI counts as compared to longer genes.

**Figure S11.**
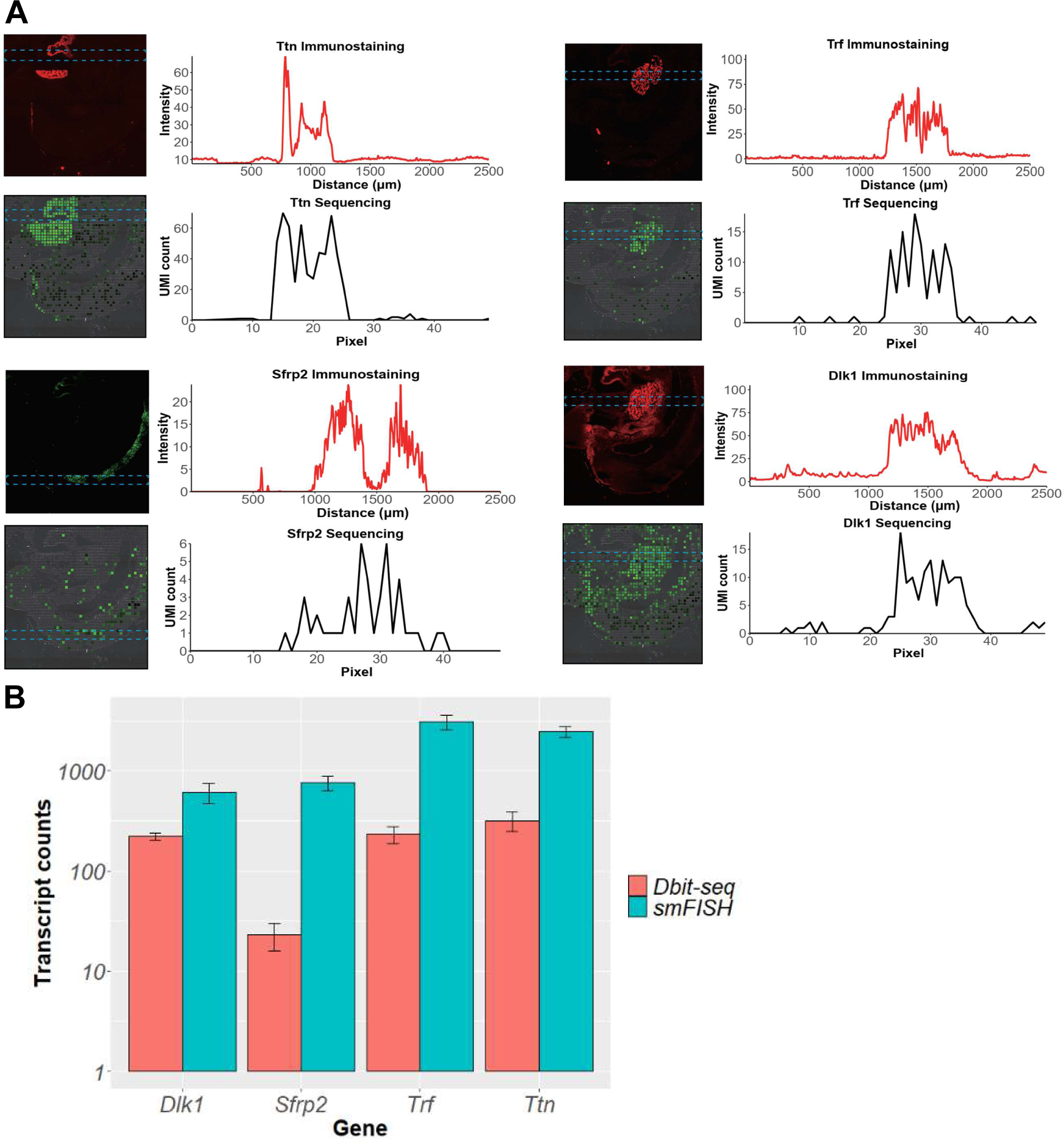
Comparison of DBiT-seq with smFISH. (A) Comparison of the DBiT-seq results of four genes (*Dlk1, Sfrp2, Trf, Ttn*) with smFISH. (B) Comparison of the transcript counts of four genes between DBit-seq and smFISH. The count was collected for an area of 306 µm x 306 µm.

**Figure S12.**
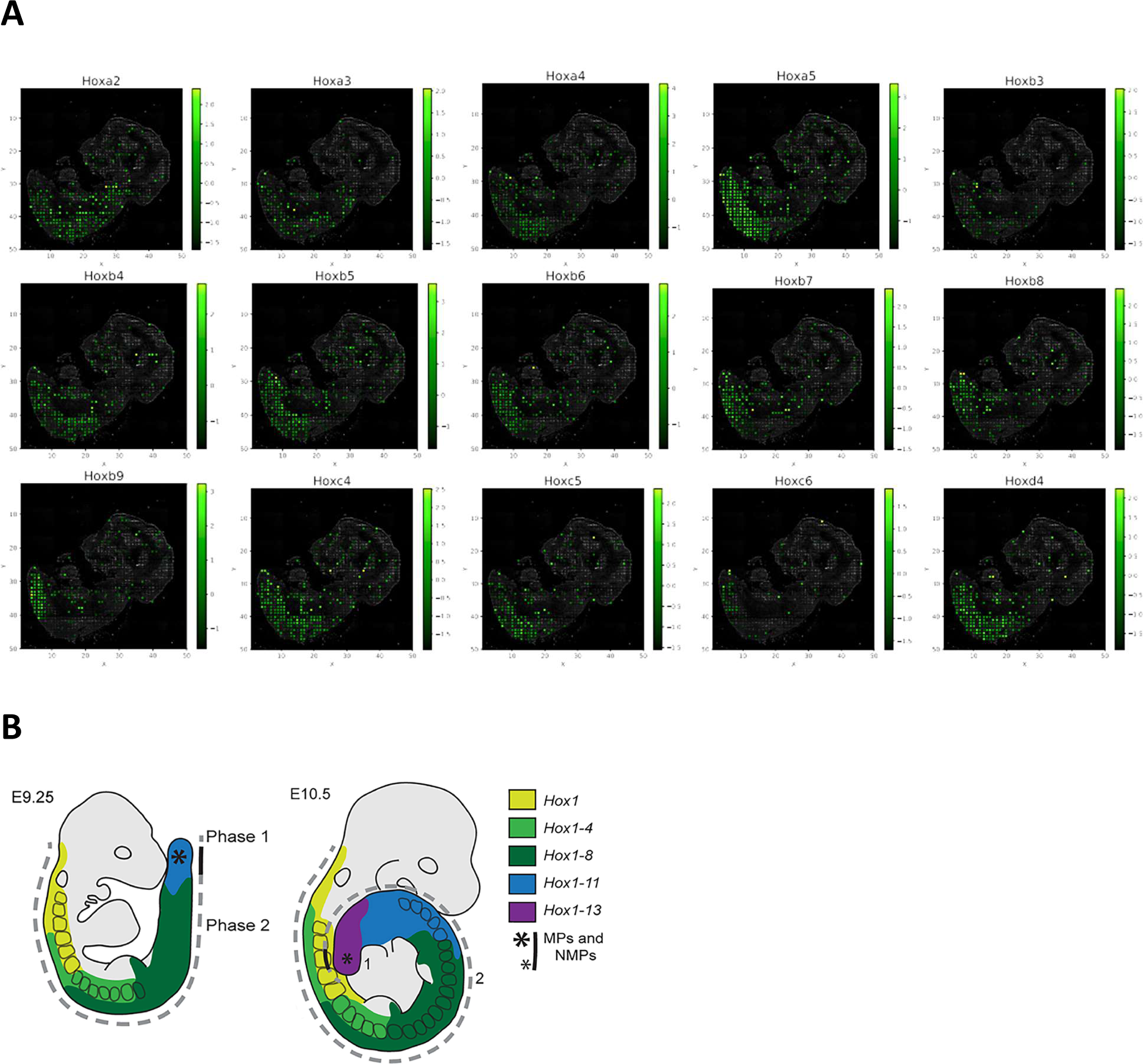
*Hox* gene patterns in a E10 mouse embryo. (A) Visualization of *Hox* gene expression. They are a family of homeobox genes that specify regions of the body plan along the anterior-posterior (head-tail) embryonic axis. We searched *Hoxa5* in eMouseAtlas website(http://www.emouseatlas.org/gxdb/dbImage/segment6/26565/detail_26565.html), which showed it was strongly associated with the tail development at mouse embryo day 10.5, which was consistent with DBiT-seq data – *Hoxa5* expression pattern. (B) Schematic illustration of the *Hox* gene codes in mouse embryos (E9-E10.5). (Deschamps J., et al., GENES & DEVELOPMENT 31:1406–1416). This review article summarized the *Hox* gene codes in mouse embryos (E9-E10.5). Higher number *Hox* genes (*Hox9-13*) are expressed in the tail region and the lower number *Hox* genes are expressed throughout the entire neutral tube extending to the hindbrain, which is consistent with the results in (A), for example, in *Hoxa2* vs *Hoxb9*. Other *Hox* genes also follow a similar trend.

**Figure S13.**
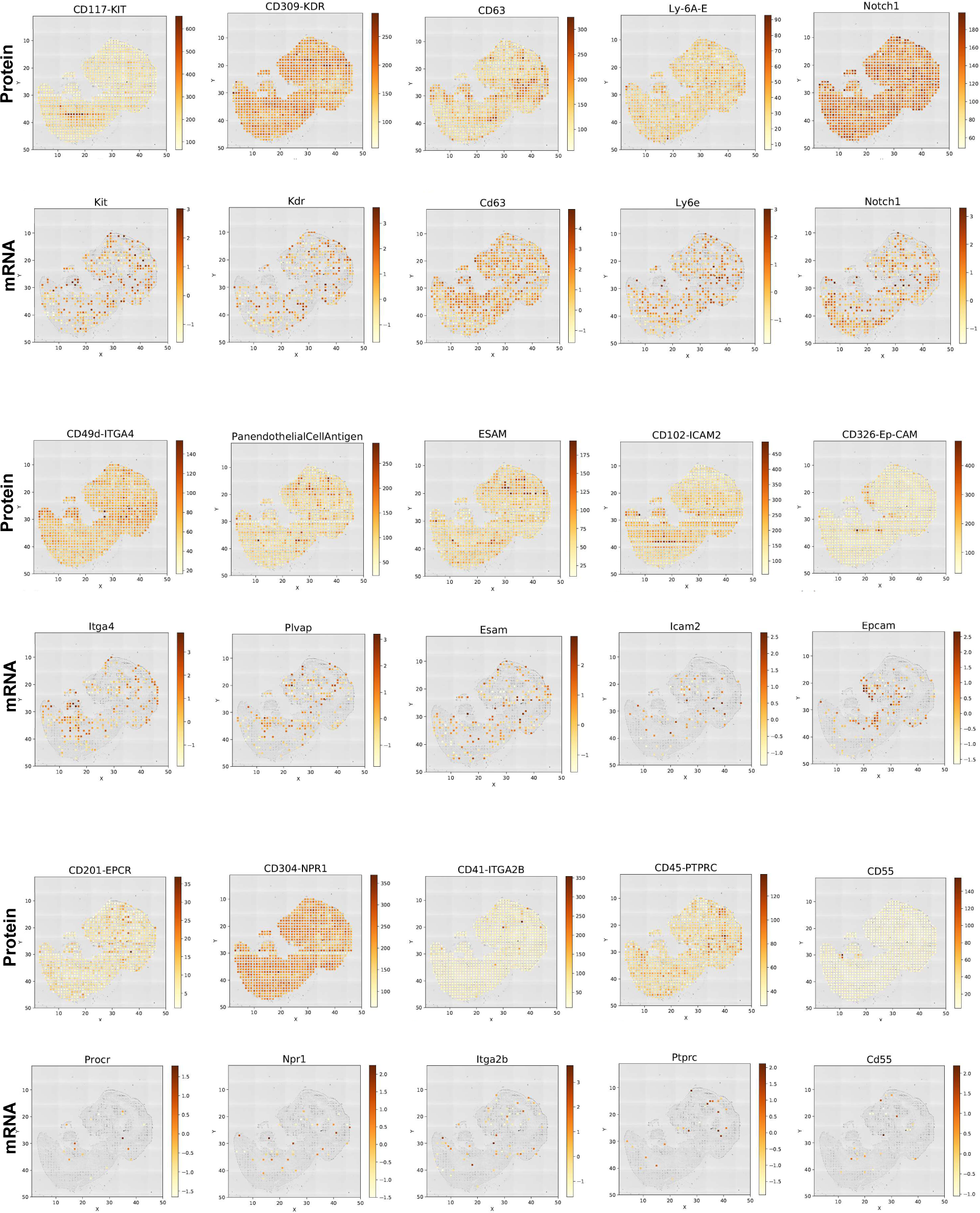
Protein and mRNA pairs as shown in Figure 2.

**Figure S14.**
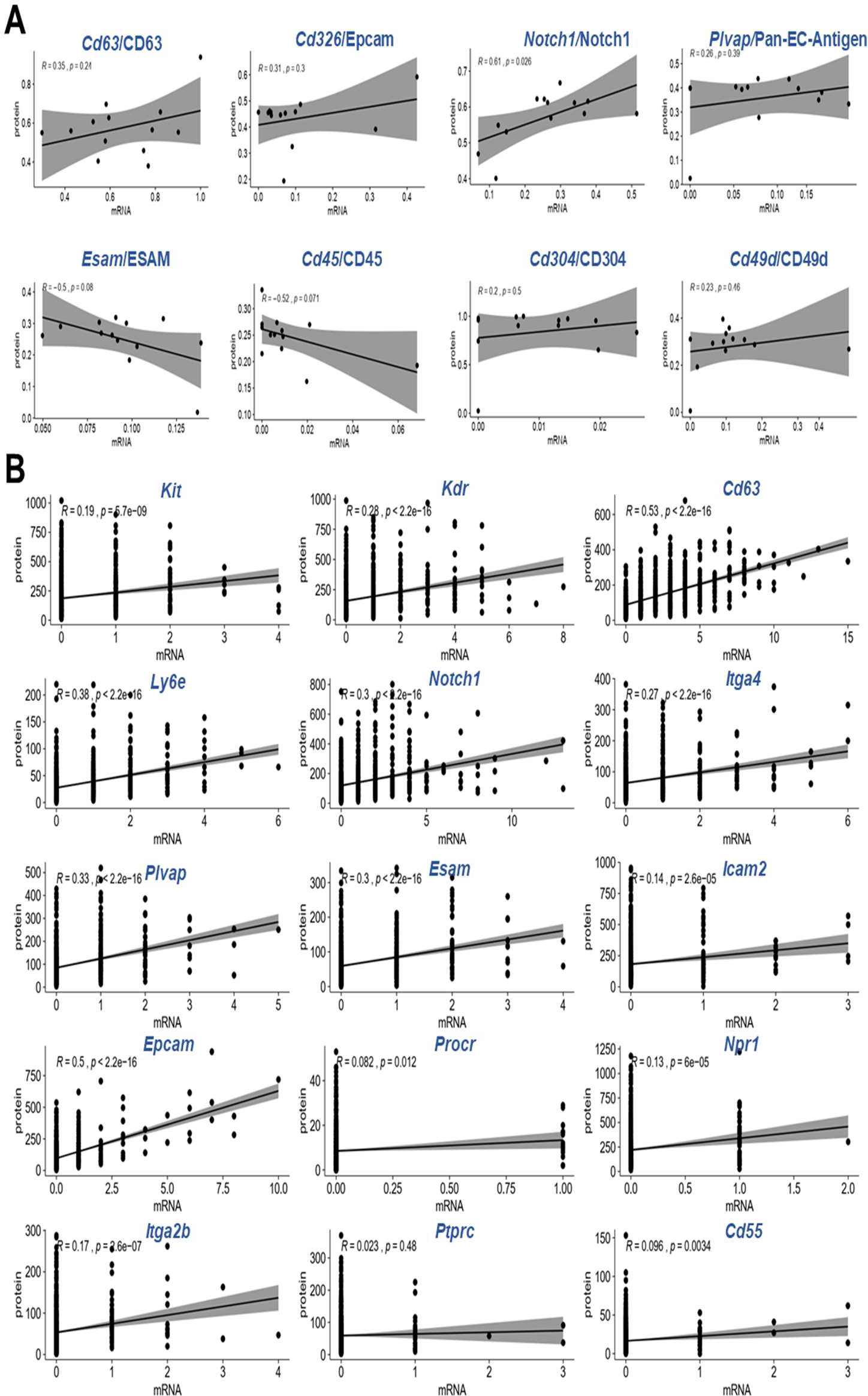
Pearson correlation analysis between mRNA and proteins pairs. (A) Pearson correlation between mRNA and proteins from all13 different tissue areas in Figure 2F. (B) Individual pixel level Pearson correlation between mRNA and proteins from Figure S8.

**Figure S15.**
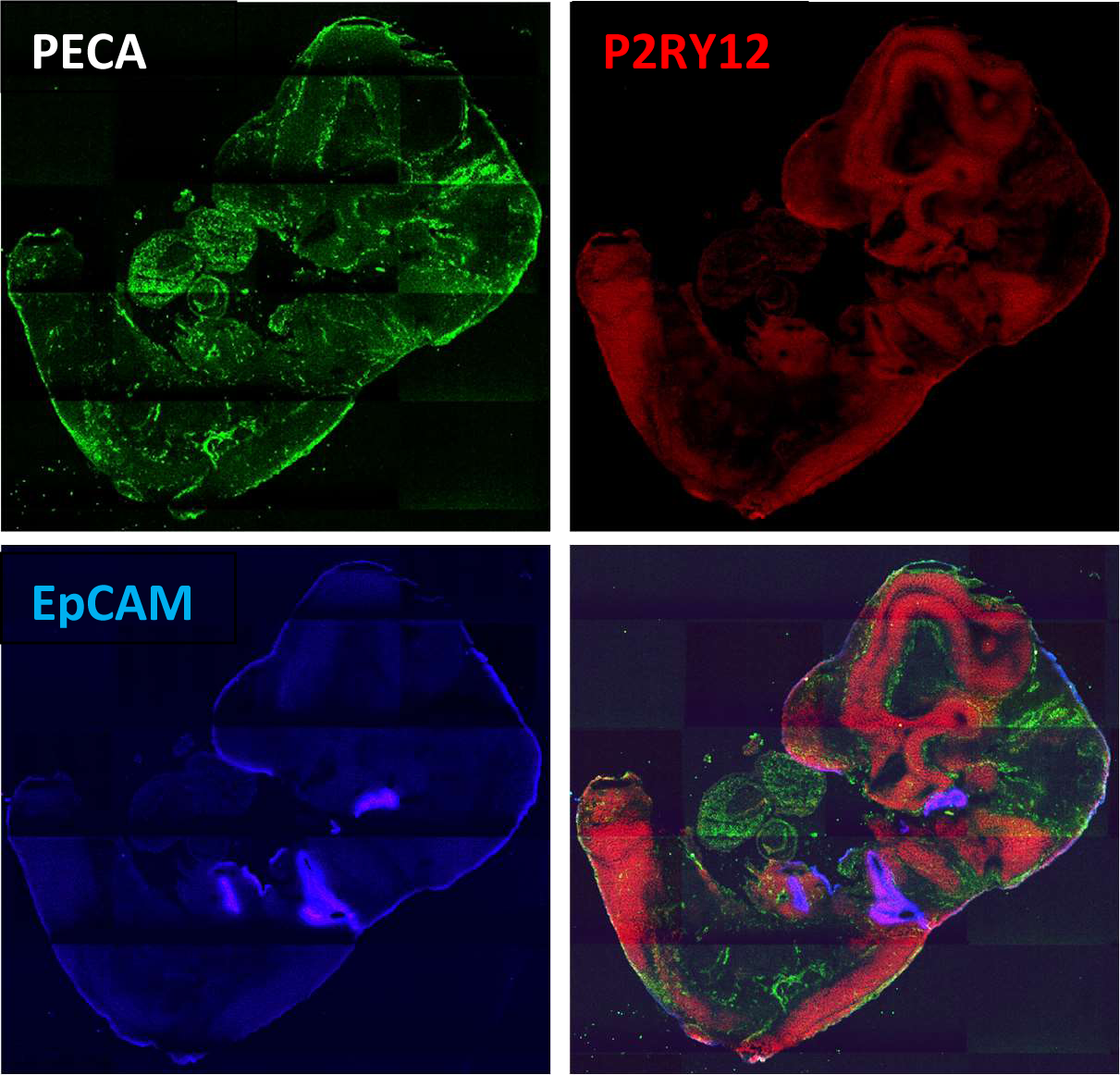
Immunofluorescence tissue staining of a neighboring tissue slide as shown in Figure 2. Pan-endothelial antigen (PECA), which marks the formation of embryonic vasculature, is expressed extensively at this stage (E10), consistent with the protein and mRNA expression revealed by DBiT-seq. EpCAM, an epithelial marker, already show up but in several highly localized regions, which were also identified by DBiT-seq (both mRNA and protein). P2RY12 (Red) is a marker for microglia in CNS, which depicts the spatial distribution of the neural system.

**Figure S16.**
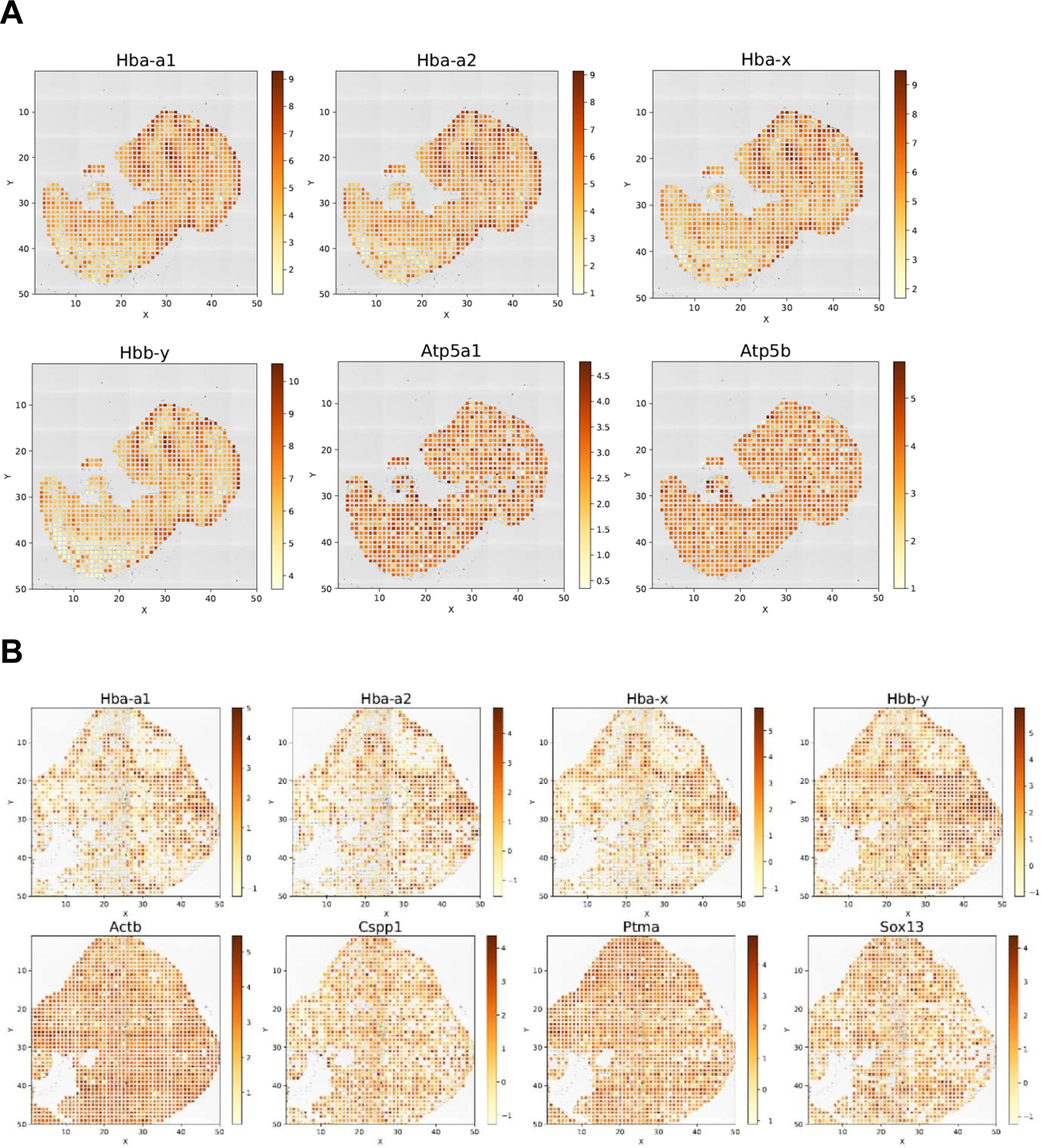
Effect of normalization on spatial mRNA expression pattern. Related to Figures 2 and 3. Pan-mRNA UMI count maps in Figures 2 and 3 correspond to the raw data without normalization and preprocessing. As expected, the variability of microfluidic flow across different channels resulted in the “banding” effect, which was similar to that observed in DNA oligomer density variability in bead-based single-cell RNA-sequencing and can be normalized to correct this effect. Because of limited sensitivity in detecting low-abundance mRNAs, the correction of “banding” effect was better evaluated by visualizing the spatial expression patterns of highly abundant genes. (A) Spatial expression maps of six high abundance genes in the 50µm E10 whole embryo DBiT-seq data visualized after standard normalization. Related to Figure 2. (B) Spatial expression maps of eight high abundance genes in the 25µm E10 embryonic brain DBiT-seq data visualized after standard normalization. Related to Figure 3. It is worth noting that these hemoglobin genes mainly in red blood cells correlated strongly with the expression pattern of Pan-Endotherlial-Cell Antigen (PECA) protein, which delineated brain microvasculature.

**Figure S17.**
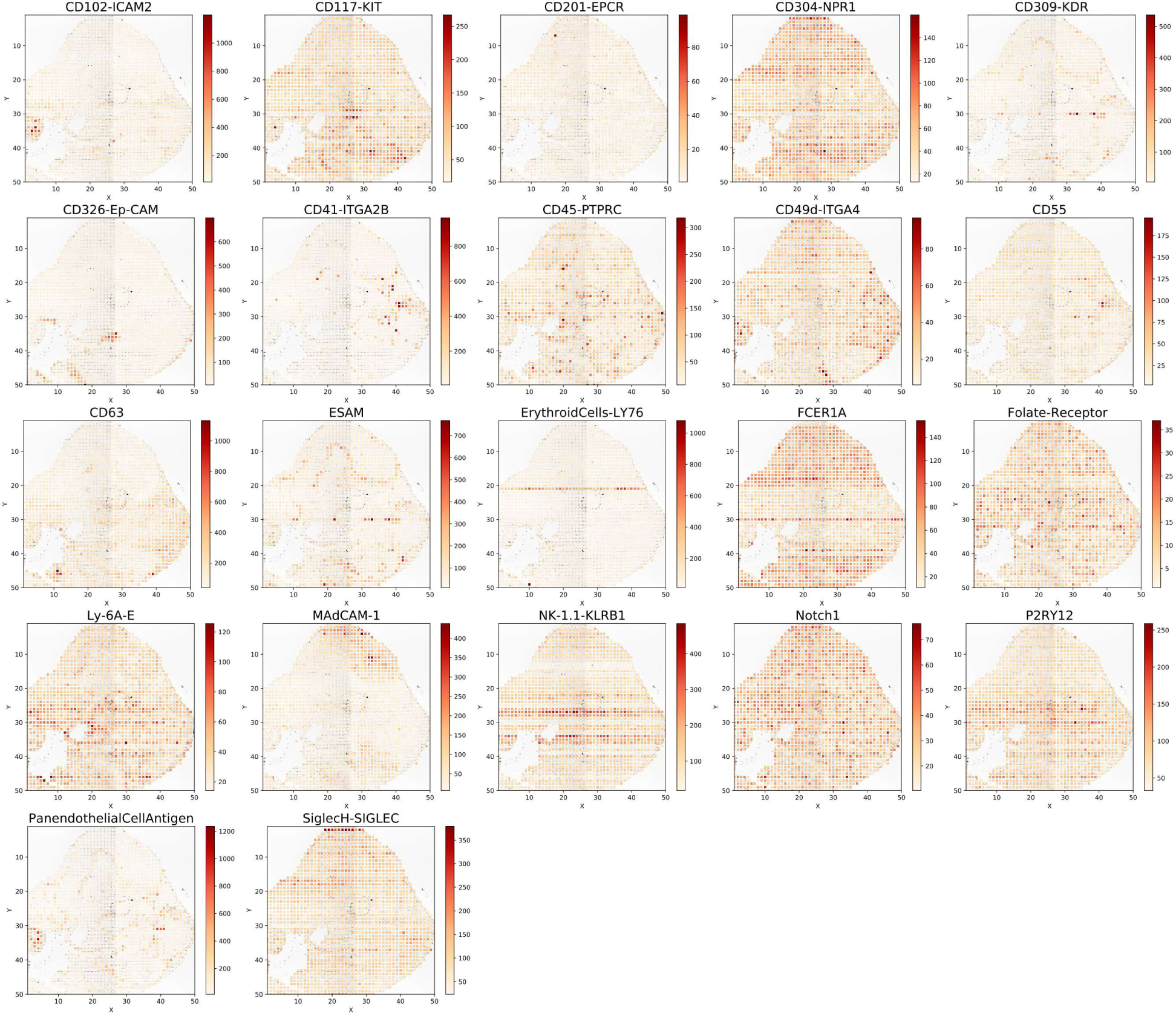
Embryonic brain protein atlas. Related to Figure 3. Protein expression of all 22 individual proteins in embryonic mouse brain (E10, 25 µm pixel size).

**Figure S18.**
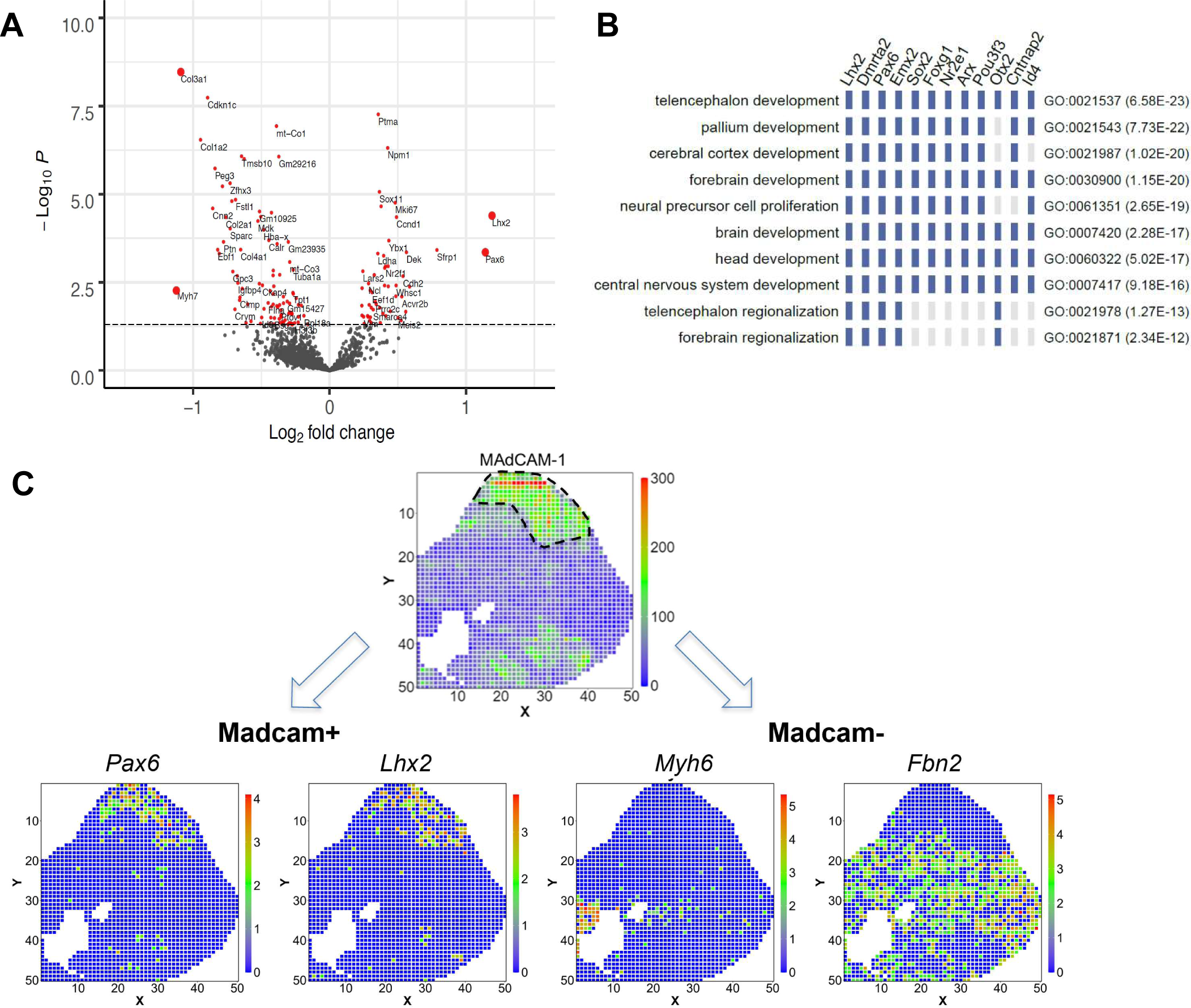
Volcano plot and representative genes from MadCAM-1+ and MadCAM-1-regions of mouse embryo brain. (A) Volcano plot showing spatial proteomics-guided differential mRNA expression analysis to search for region-specific transcriptomic signature. (B) Pathway analysis of MAdCAM1+ forebrain region. (C) Visualization of four transcripts identified by the analysis in (B). Pax6 and Lhx2 are enriched in the MAdCAM-1+ region of the forehead. *Myh6* and *Pbn2* were enriched in MAdCAM-1-region.

**Figure S19.**
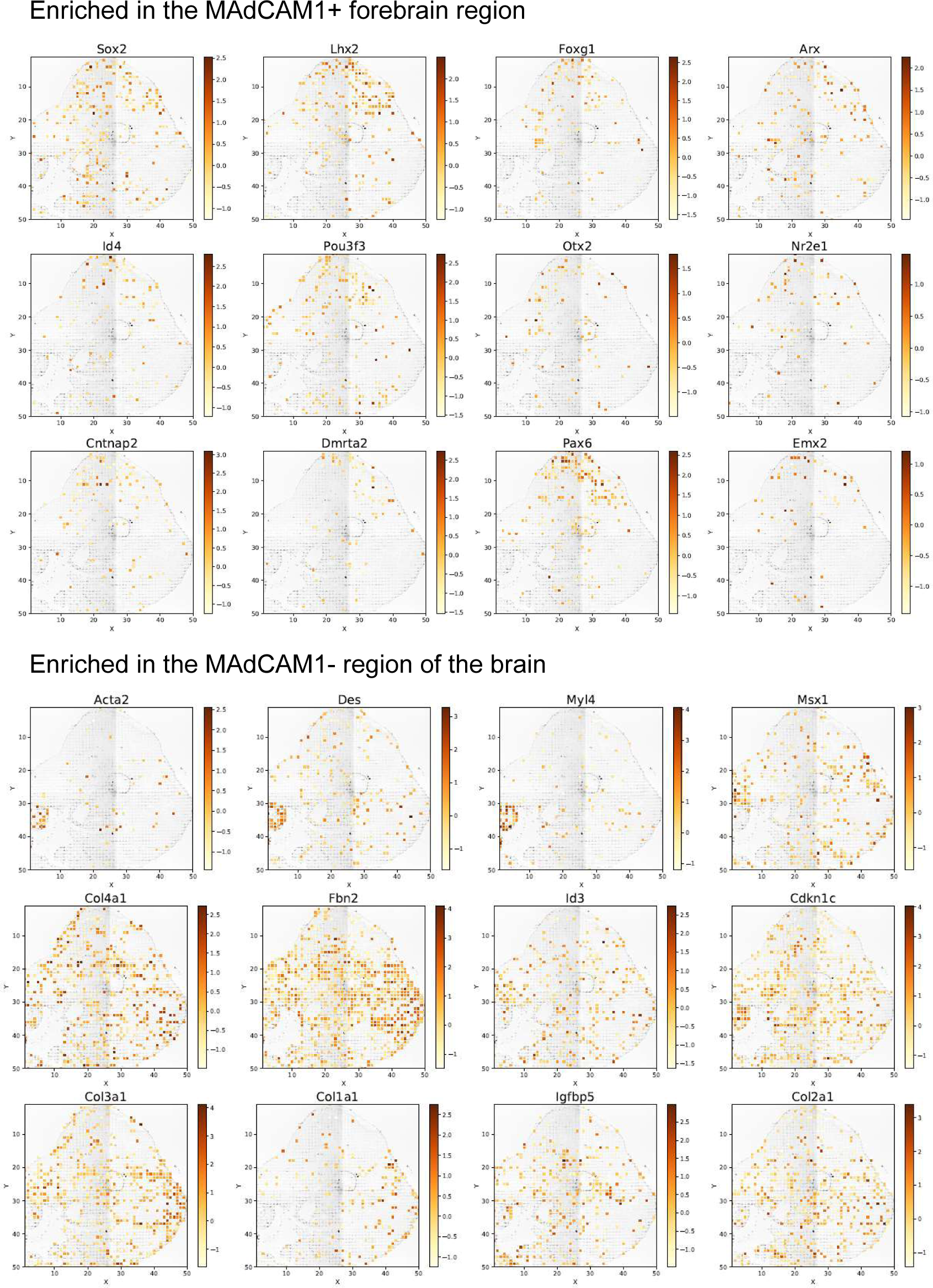
Differential gene expression in MAdCAM-1+ vs-regions. Related to Figure 4. Spatial expression of mRNAs enriched in MAdCAMpl rotein+ or-regions identified by spatial proteomics-guided differential gene expression analysis.

**Figure S20.**
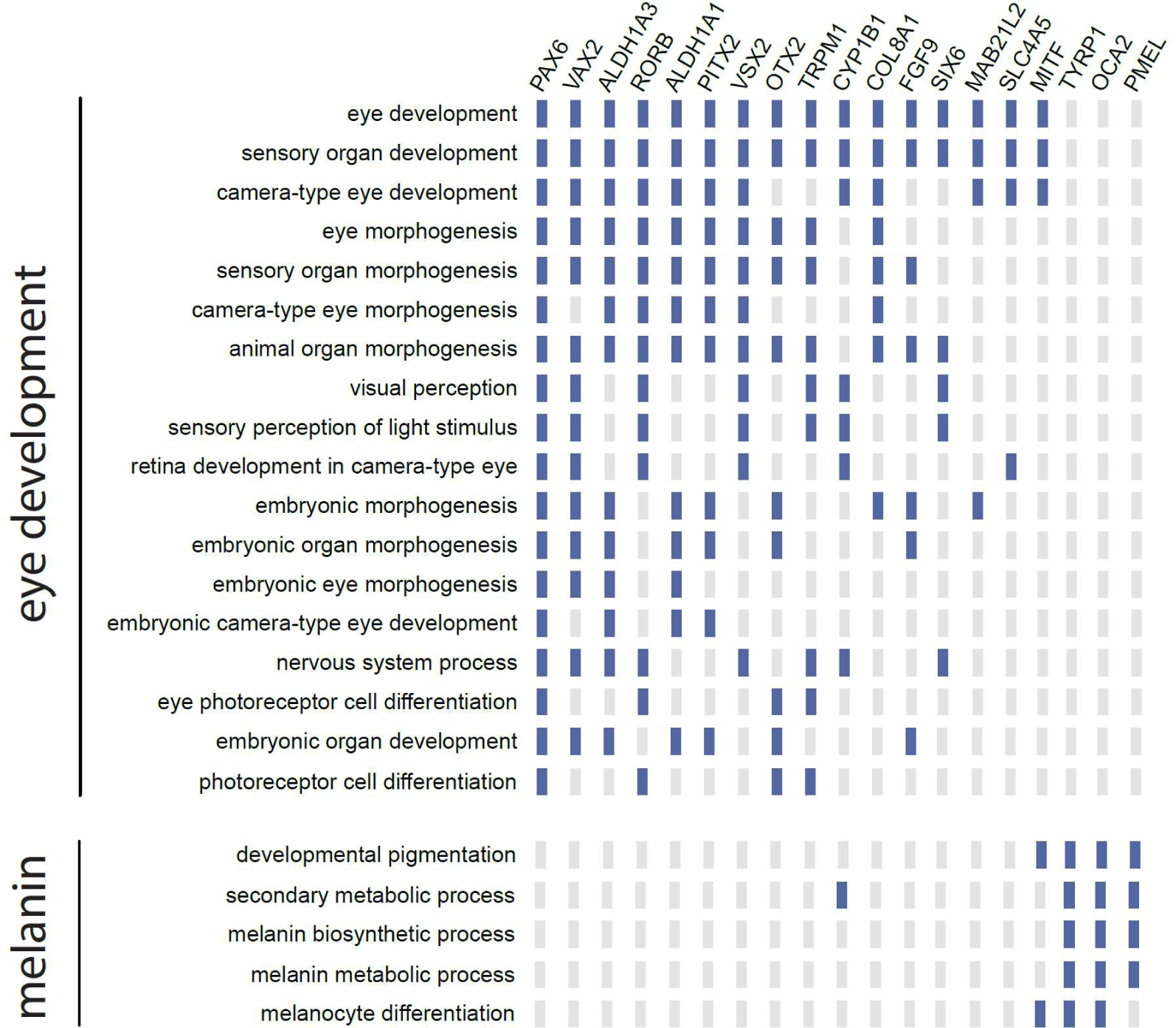
Pathway analysis of data shown in Figure 4. Pathway analysis and differentially expressed genes. The most profound pathways identified are eye development including the associated sensory nerves and melanin.

**Figure S21.**
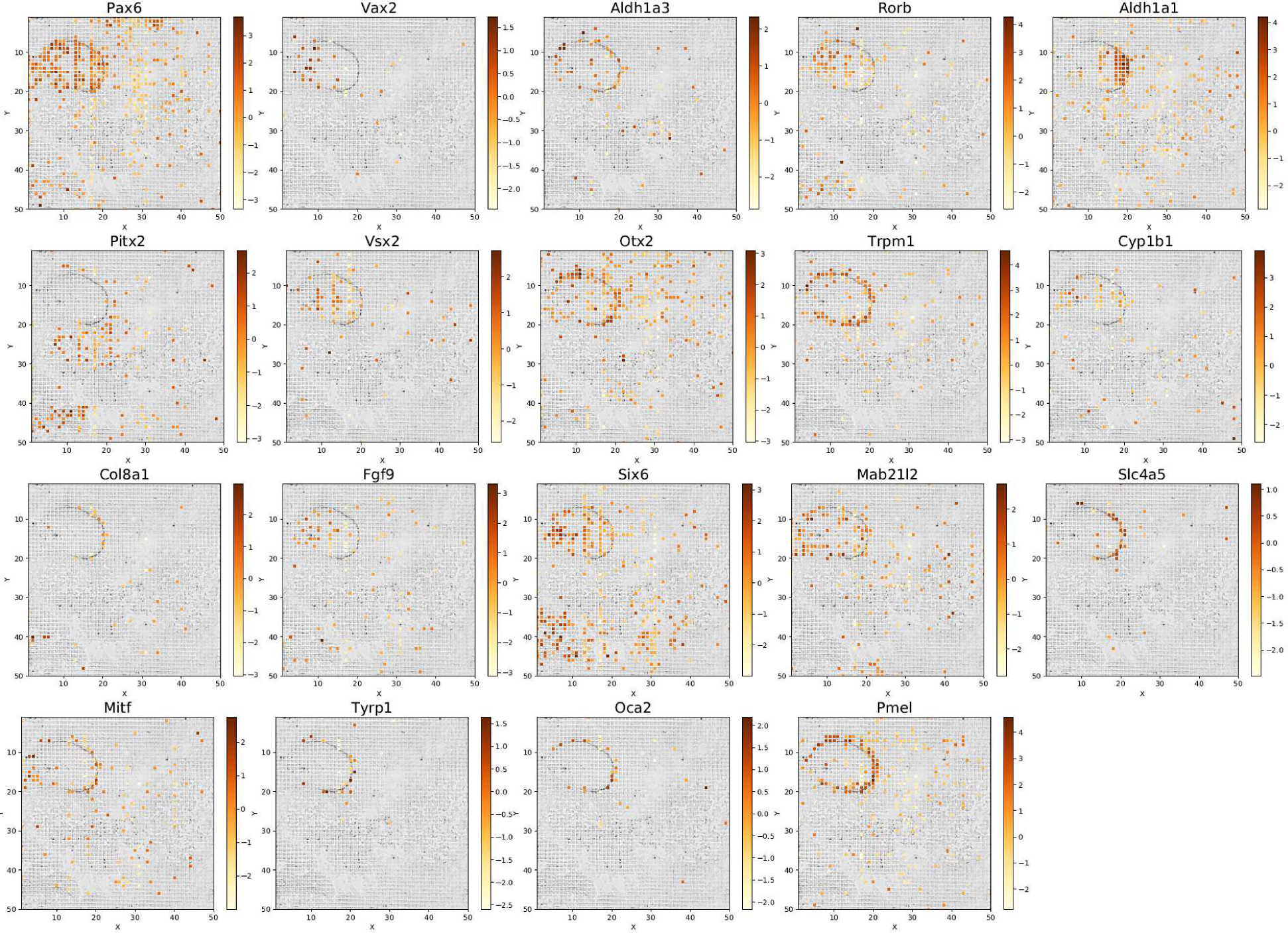
Spatial mRNA expression patterns around the eye. Related to Figure 5. Spatial expression patterns of top ranked genes identified in the eye development region (10 µm pixel).

**Figure S22.**
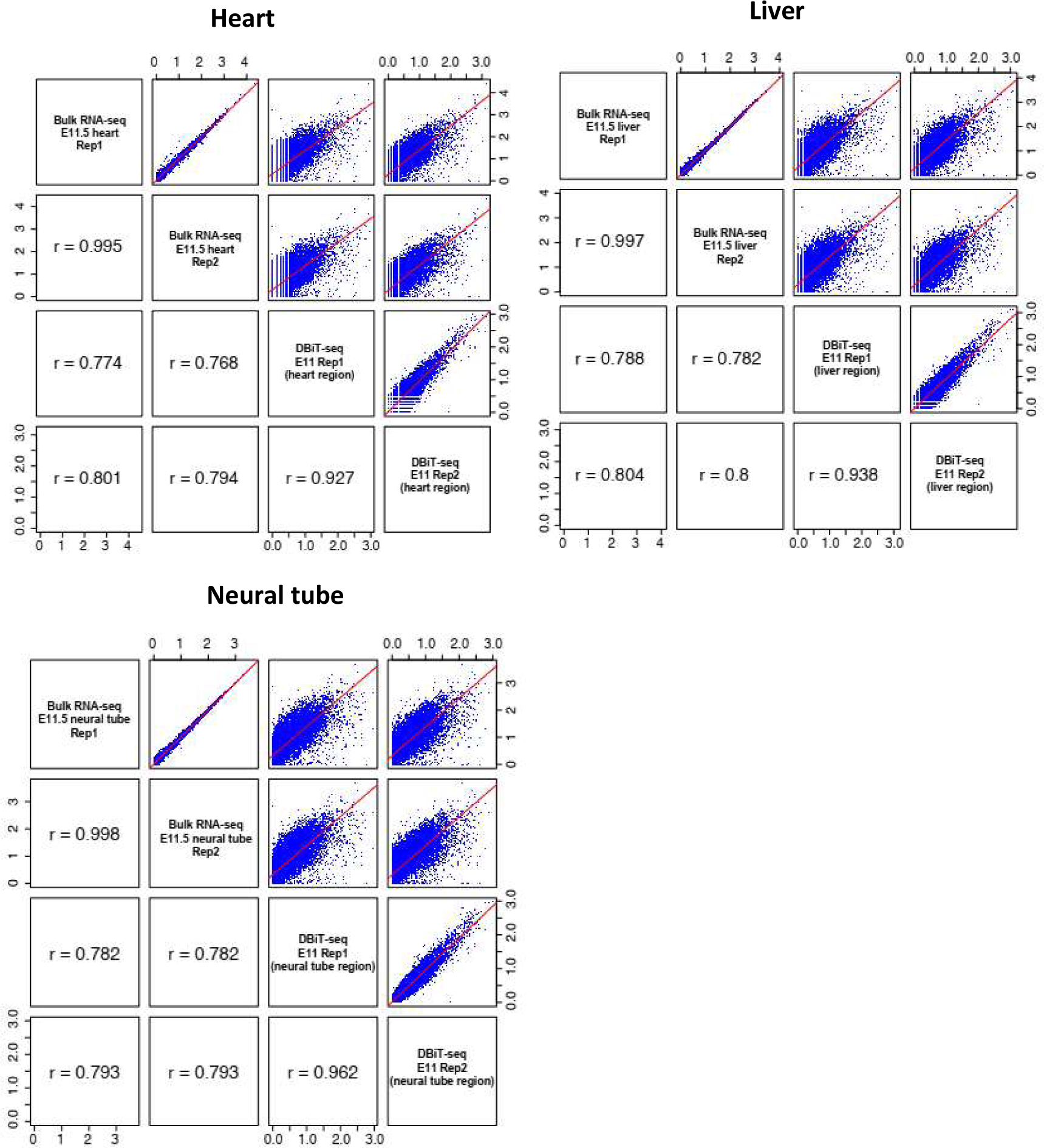
Pearson correlation between pseudo-bulk expression of our embryo data and ENCODE bulk RNA-seq datasets. Three featured regions (heart, liver, and neural tube) of mouse embryo (E11 and E11.5) were manually selected and pseudo-bulk expression was generated and compared to ENCODE bulk RNA-seq datasets. Average correlation coefficients for each tissue regions were: 0.74±0.02 for heart, 0.79±0.01 for liver and 0.79±0.01 for neural tube. A strong agreement between the two datasets were observed.

**Figure S23.**
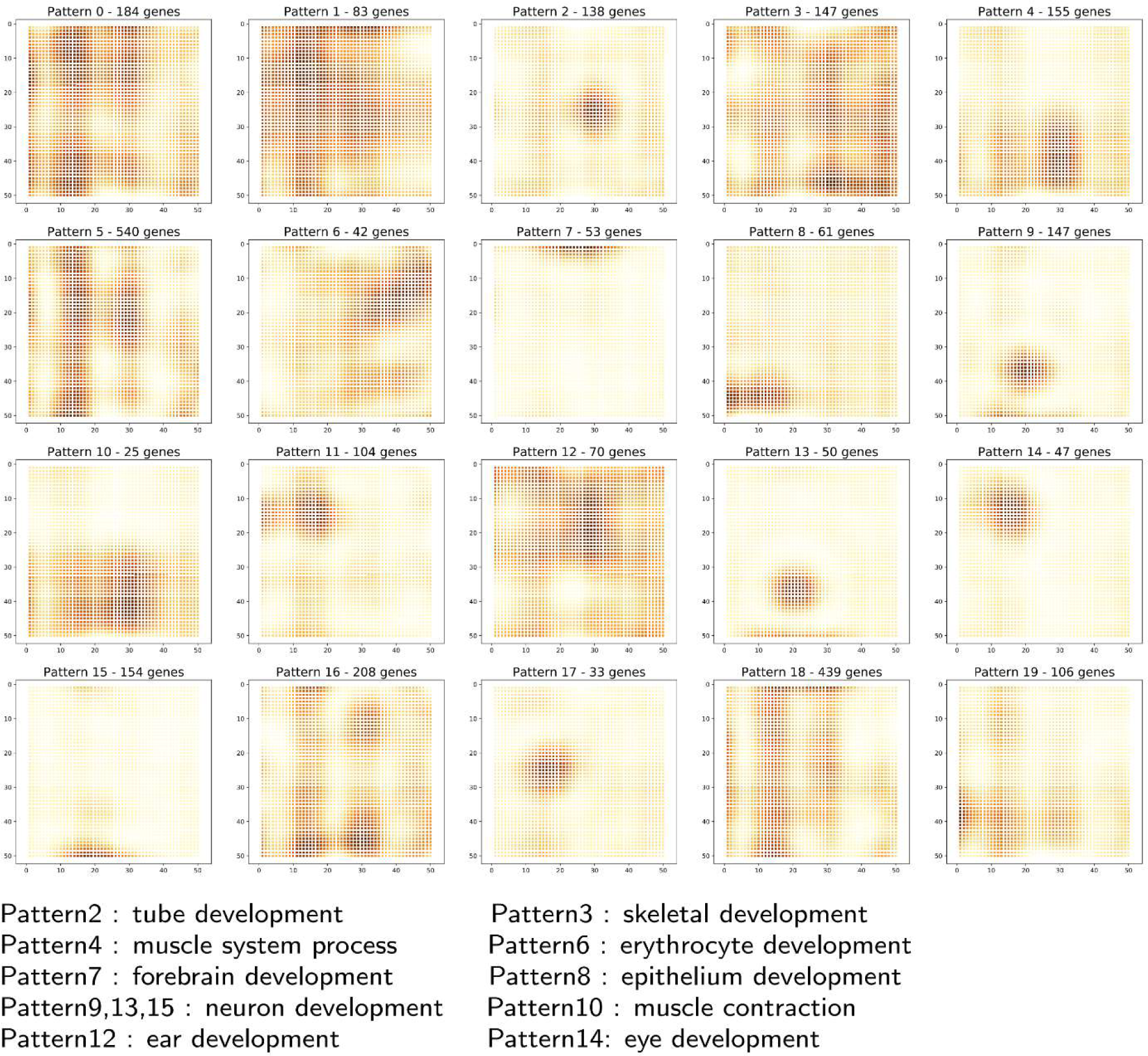
All features identified around the eye. Related to Figure 4. Features identified using SpatialDE in region around the eye field from the spatial transcriptome data. Pixel size = 10 µm.

**Figure S24.**
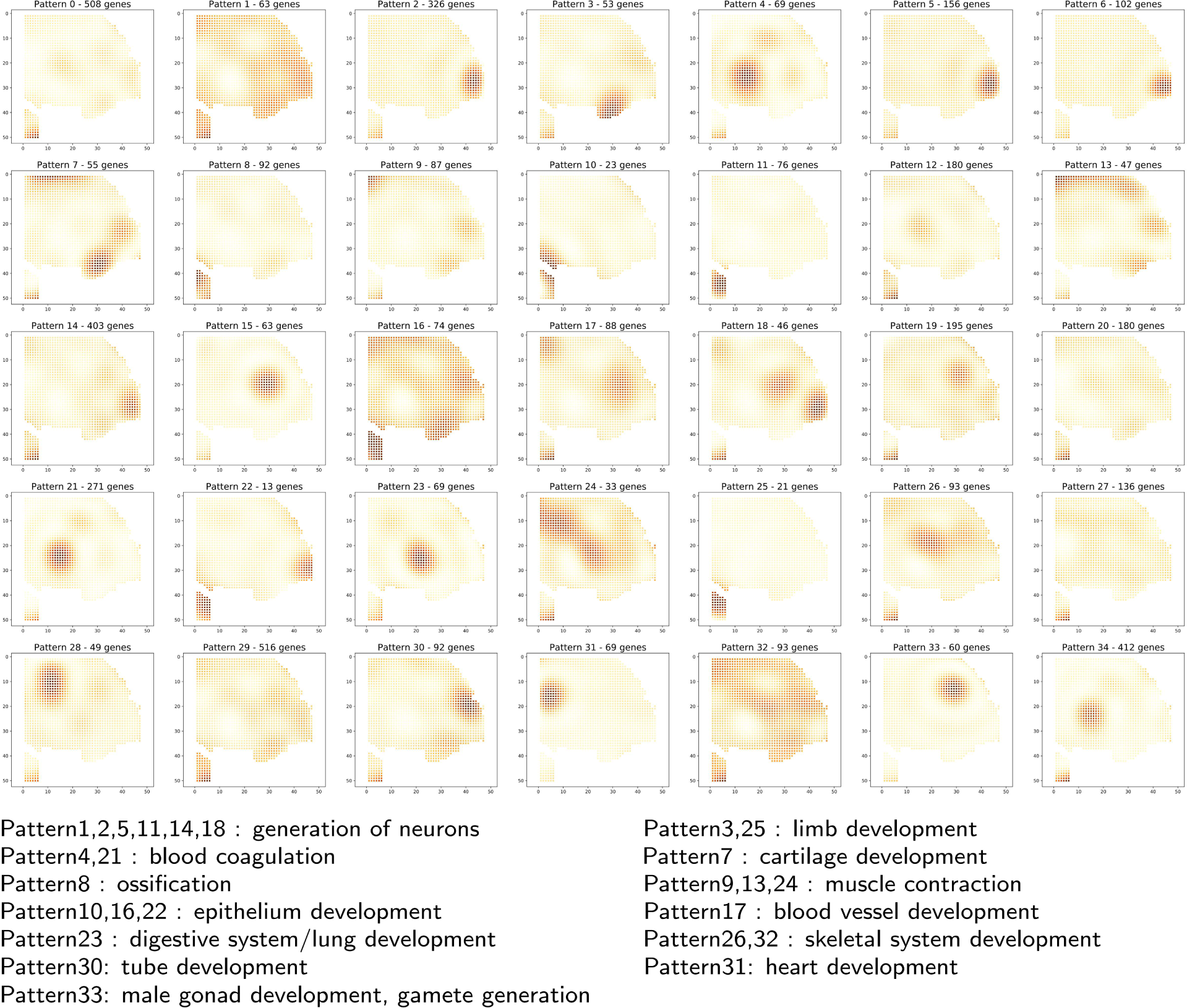
All features identified in an E12 mouse embryo DBiT-seq data. Related to Figure 6. Feature identified in an E12 mouse embryo tissue slide using SpatialDE. Pixel size = 50 µm.

**Figure S25.**
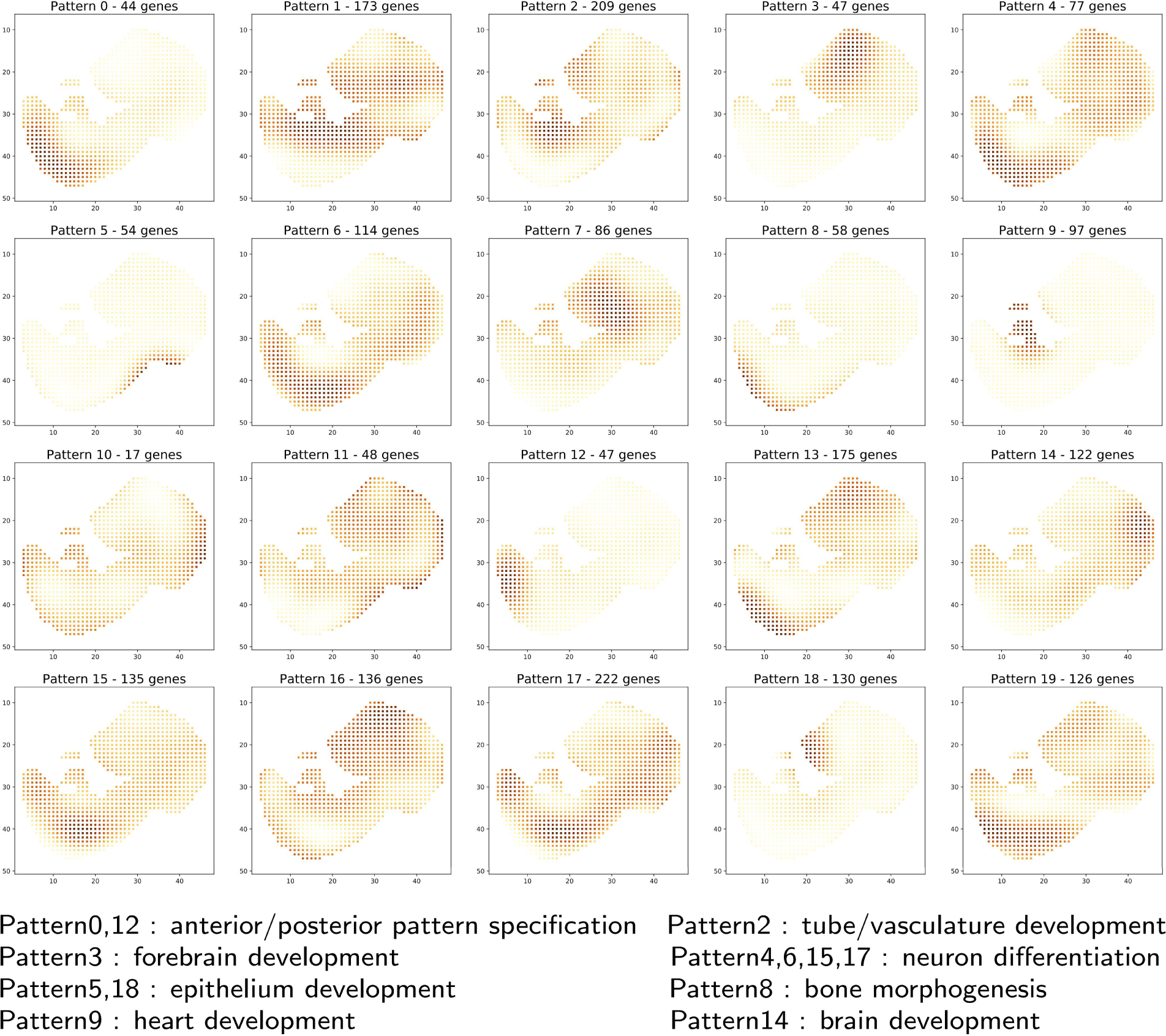
All features identified in a E10 whole mouse embryo DBiT-seq. Features identified in the whole mouse embryo (E10) tissue slide using SpatialDE. Pixel size = 50 µm.

**Figure S26.**
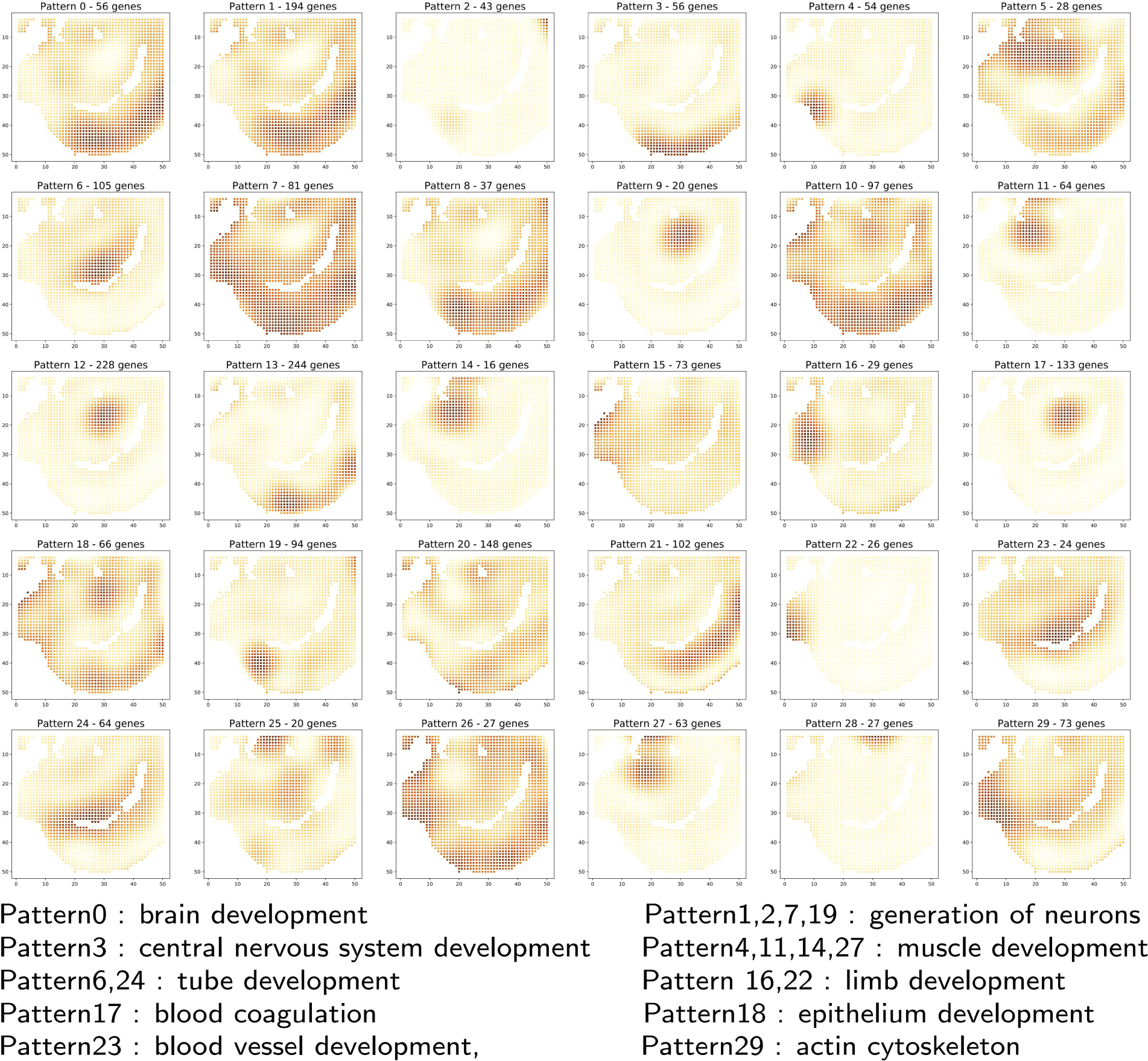
All features identified in the E11 mouse embryo as shown in Figure 6. Features identified in the whole mouse embryo (E11) tissue slide using SpatialDE. Pixel size = 25 µm.

**Figure S27.**
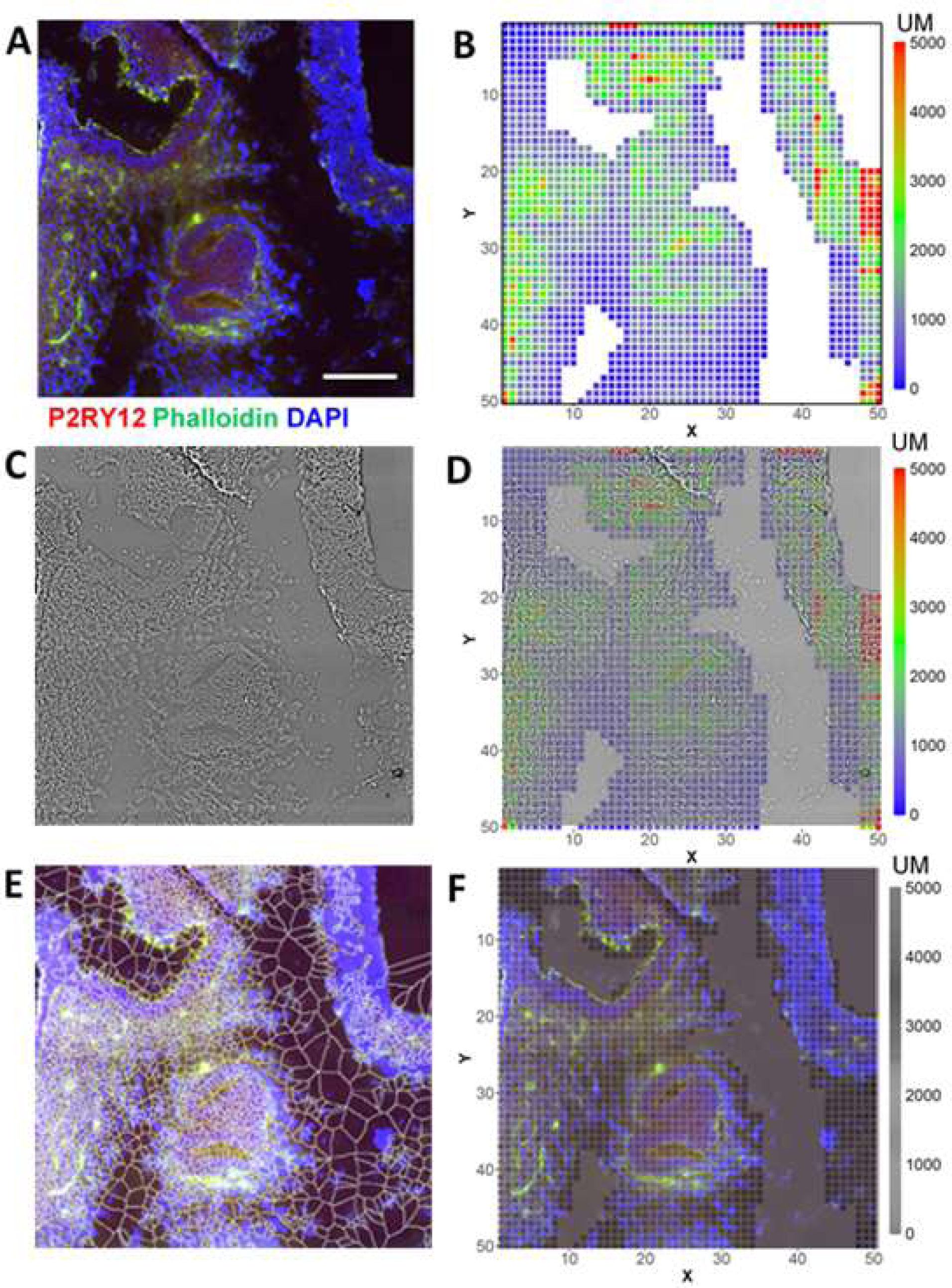
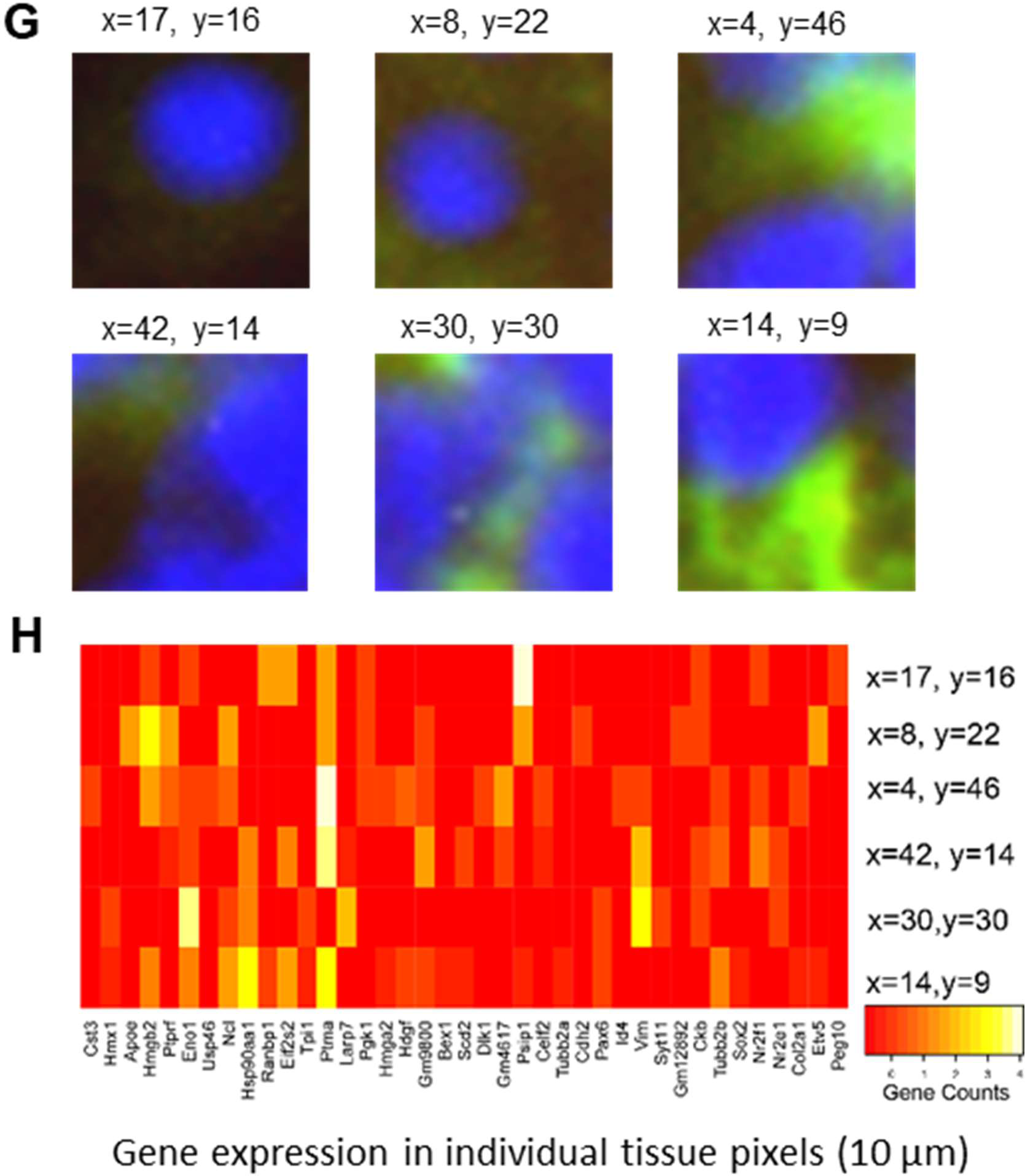
Transcriptome analysis of fluorescent stained mouse embryo (E11). (A) Fluorescent image of pre-stained mouse embryo tissue section. Tissue sections were first stained with DAPI, Phalloidin and P2RY12. (B) UMI heatmap of the same tissue section. (C) Bright filed image of mouse embryo tissue section. (D) Overlap of bright filed image with UMI heatmap. (E) Cell segmentation of the fluorescent stained section. Segmentation was carried out using ImageJ. Four representatives of differentially expressed genes that matches well with whole transcriptome clustering. (F) Pixelized fluorescent images. (G) Representative fluorescent image of individual pixels (10µm). (H) Gene expression profiles of selected pixels from (G).

### Supplementary Tables

**Table S1.** List of antibody-derived DNA tags (ADTs). (See the Excel file attached)

**Table S2.** List of PCR oligo and DNA barcode sequences. (See the Excel file attached)

**Table S3.** Summary of buffers, chemicals, and reagents. (See the Excel file attached).

**Table S4.** Summary of samples and quality evaluation.

| Sample ID | Annotation | # of Pixels under tissue | # of unique genes present | Average # of unique molecules per pixel | Average # of unique genes per pixel |
| --- | --- | --- | --- | --- | --- |
| 10 μm | E10 Eye (10 μm) 1 | 2,500 | 22,969 | 4,909 | 2,068 |
| 50 μm | E10 Whole (50 μm) 1 | 902 | 22,243 | 9,427 | 3,478 |
| 0628cL | E12 Tail (50 μm) | 1,667 | 22,571 | 3,942 | 2,000 |
| 0702cL | E10 Whole (50 μm) 2 | 901 | 22,846 | 11,937 | 4,132 |
| 0713cL-P | E10 Brain (25 μm) | 1,789 | 18,787 | 876 | 646 |
| 0719cL | E10 Eye (10 μm) 2 | 2,500 | 20,268 | 2,151 | 1,116 |
| 0725e10cL | E10 Whole (50 μm) 3 | 936 | 22,802 | 12,314 | 4,170 |
| E11-1L | E11 Tail (25 μm) 1 | 1,837 | 20,849 | 2,788 | 1,375 |
| E11-2L | E11 Tail (25 μm) 2 | 1,662 | 21,890 | 4,689 | 2,101 |
| E11-FL-1L | E11 Brain (10 μm) | 1,840 | 21,337 | 1,421 | 827 |
| E11-FL-2L | E11 Tail (10 μm) | 2,109 | 20,257 | 1,320 | 763 |

